# A scalable CRISPR-Cas9 gene editing system facilitates CRISPR screens in the malaria parasite *Plasmodium berghei*

**DOI:** 10.1101/2024.04.20.590404

**Authors:** Thorey K. Jonsdottir, Martina S. Paoletta, Takahiro Ishizaki, Sophia Hernandez, Maria Ivanova, Alicia Herrera Curbelo, Paulina A. Saiki, Martin Selinger, Debojyoti Das, Johan Henriksson, Ellen S.C. Bushell

**Author notes:** Corresponding authors: Ellen S.C. Bushell, Thorey K. Jonsdottir, Martina S. Paoletta, Umeå University, Umeå, Sweden. These authors contributed equally to this work (Thorey K. Jonsdottir, Martina S. Paoletta).

## Abstract

Many *Plasmodium* genes remain uncharacterised due to low genetic tractability. Previous large scale knockout screens have only been able to target about half of the genome in the more genetically tractable rodent malaria parasite *Plasmodium berghei*. To overcome this limitation, we have developed a scalable CRISPR system called PbHiT, which uses a single cloning step to generate targeting vectors with 100 bp homology arms physically linked to a guide RNA (gRNA) that effectively integrate into the target locus. We show that PbHiT coupled with gRNA sequencing robustly recapitulates known knockout mutant phenotypes in pooled transfections. Furthermore, we provide vector designs and sequences to target the entire *P. berghei* genome and scale-up vector production using a pooled ligation approach. This work presents for the first time a tool for high-throughput CRISPR screens in *Plasmodium* for studying the parasite’s biology at scale.

## Introduction

Malaria is a mosquito-borne disease caused by apicomplexan *Plasmodium* parasites with over 249 million cases and 608,000 deaths reported annually^1^. Rational design of new and urgently needed interventions requires the identification of novel parasite targets involved in parasite infection or transmission. Reverse genetics is a powerful tool to identify essential genes and determine gene function in genetically tractable organisms, including those of the Apicomplexa phylum^2^. The murine species *Plasmodium berghei* is widely used as an *in vivo* malaria model that facilitates studies of the entire parasite life cycle and benefits from a higher transfection efficiency compared to the human parasite *Plasmodium falciparum*^3^.

Targeted reverse genetic screens have been conducted in *Plasmodium* including the systematic knockout of exported proteins in *P. falciparum*^4^ and all predicted protein kinases and phosphatases in *P. berghei* ^5,6^, as well as the conditional inactivation of unstudied genes encoding non-secreted proteins on chromosome 3 in *P. falciparum* ^7^. These gene-by-gene studies are laborious and not scalable to the entire genome. The *<u>Plasmo</u>dium* <u>Ge</u>netic <u>M</u>odification (*Plasmo*GEM) project provides knockout vectors for more than half of the ∼5,000 *P. berghei* genes. The *Plasmo*GEM vectors owe their efficiency to long homology arms and are equipped with molecular barcodes^8,9^. *Plasmo*GEM gene knockout screens using barcode sequencing have identified parasite genes essential for *in vivo* asexual blood stage growth, and the sexual and liver developmental stages required for mosquito transmission^9–11^. However, there are substantial limitations to the genetic toolbox available to query gene function at scale and, as a result, our knowledge of *Plasmodium* gene function.

In the last decade, CRISPR-Cas9 has transformed experimental genetics by providing an efficient and versatile way of manipulating genomes and was rapidly adapted to the study of *Plasmodium*^12,13^. Using Cas9 to precisely engineer a double-strand break (DSB) enhances the efficiency of gene editing in *Plasmodium* when using a standard length (≤1000 bp) homology region (HR)^14^. Since *Plasmodium* lacks the pathway for canonical non-homologous end-joining (c-NHEJ), any CRISPR-Cas9 mediated edit requires a homology directed repair (HDR) template to facilitate DSB repair^15^. This prohibits the adoption of standard CRISPR-Cas9 disruption screens that rely on c-NHEJ, which introduces insertion and deletion mutations during repair. The HDR template must instead be delivered into the parasite together with the corresponding guide RNA (gRNA) and can be supplied in the same genetic vector that carries the gRNA or on a separate linear or circular DNA molecule^12,13,16–19^. The development of CRISPR-Cas9 screens in *Plasmodium* requires a scalable system where the gRNA and the HR are physically linked to ensure that each parasite receives a matched gRNA and HR during pooled transfections.

In the related parasite *Toxoplasma gondii*, NHEJ-dependent CRISPR-Cas9 gene disruption screens have greatly accelerated functional annotation of its genome^20–24^. Recently, the development of a *T. gondii* high-throughput tagging CRISPR-Cas9 system, which physically couples the gRNA and HR in a single scalable vector, has paved the way for HDR-mediated CRISPR-Cas9 screens^25^. However, to implement this system in *Plasmodium* parasites, several technical roadblocks must be overcome.

Here, we present an optimised CRISPR-Cas9 system for the malaria parasite *P. berghei*. We demonstrate robust and efficient gene editing using a HDR-template with short 100 bp homology arms and adopt this to develop the *P. berghei* high-throughput (PbHiT) CRISPR-Cas9 vector system. The synthetic fragment containing the gene-specific gRNA and HR region can be cloned in a single, scalable step. The PbHiT system is versatile and can be used for gene knockout and epitope tagging, with transgenic parasites typically appearing within four to six days post-transfection. We provide a protocol for producing pools of targeting vectors, gRNA and HR sequences to target the entire *P. berghei* protein-coding genome. Furthermore, we couple the PbHiT system with gRNA sequencing and deploy it in CRISPR screens, where we use next-generation sequencing (NGS) to monitor the growth of knockout mutant pools within the bloodstream of infected mice. In summary, the PbHiT is an agile and effective system that is equally well-appointed to generate single-gene tagged or knockout mutant lines as it is to facilitate pooled transfections for CRISPR screens. To our knowledge, this represents the first genetic system that enables high-throughput CRISPR screens in malaria parasites.

## Results

### Efficient CRISPR-Cas9 editing using 100 bp homology arms

To develop an improved *P. berghei* CRISPR-Cas9 editing system, we first modified the existing *Plasmodium yoelii* pYCm CRISPR-Cas9 vector^26^ by replacing the *P. yoelii* U6 promoter with the endogenous *P. berghei* U6 promoter to drive gRNA expression. The resulting vector, pPbU6-hdhfr/yfcu-Cas9, encodes *Streptococcus pyogenes* Cas9 (spCas9) together with the dual positive/negative selection marker human dihydrofolate reductase/yeast cytosine deaminase and uridyl phosphoribosyl transferase (*hdhfr/yfcu*). To evaluate the effect of using the *P. berghei* specific U6 promoter on gene editing efficacy, we constructed both pYCm and pPbU6-hdhfr/yfcu-Cas9 vectors expressing the same gRNA to introduce a 3x haemagglutinin (3xHA) epitope tag at the 3’ end of the *P. berghei* gene encoding rhoptry-associated protein 2/3 (*rap2/3*, PBANKA_1101400). We simultaneously tested two methods for delivery of a HDR-template carrying ∼500 bp homology arms. In the one-plasmid approach, the repair template was cloned into the vector that carries the gRNA and Cas9. In the PCR-template approach, the repair template was amplified by PCR, and the amplicon was co-transfected with the pPbU6-hdhfr/yfcu-Cas9 or pYCm vector (**Fig. 1A i**), thus circumventing the subcloning of the repair template. To compare the transfection efficiency between methods, we used the time it took the parasites to reach a parasitemia of >0.5% in the bloodstream of pyrimethamine-treated mice following transfection. This is the lowest parasitemia at which we can accurately enumerate parasites, and it was used as a proxy measurement for the number of parasites initially edited. There was only a marginal difference in efficacy between parasites edited using the PbU6 compared to the PyU6 promoter when using the one-plasmid approach. In contrast, delivering the repair template using the one-plasmid compared to the PCR-template approach had a significant positive impact (p = 0.0002) on transfection efficacy, where parasitemia reached >0.5% 1-3 days earlier for the one-plasmid approach (**Fig. 1A i**). Genotyping PCRs of the *rap2/3* target locus confirmed successful integration of the 3xHA tag under all conditions tested (**Fig. S1A**).

**Figure 1.**
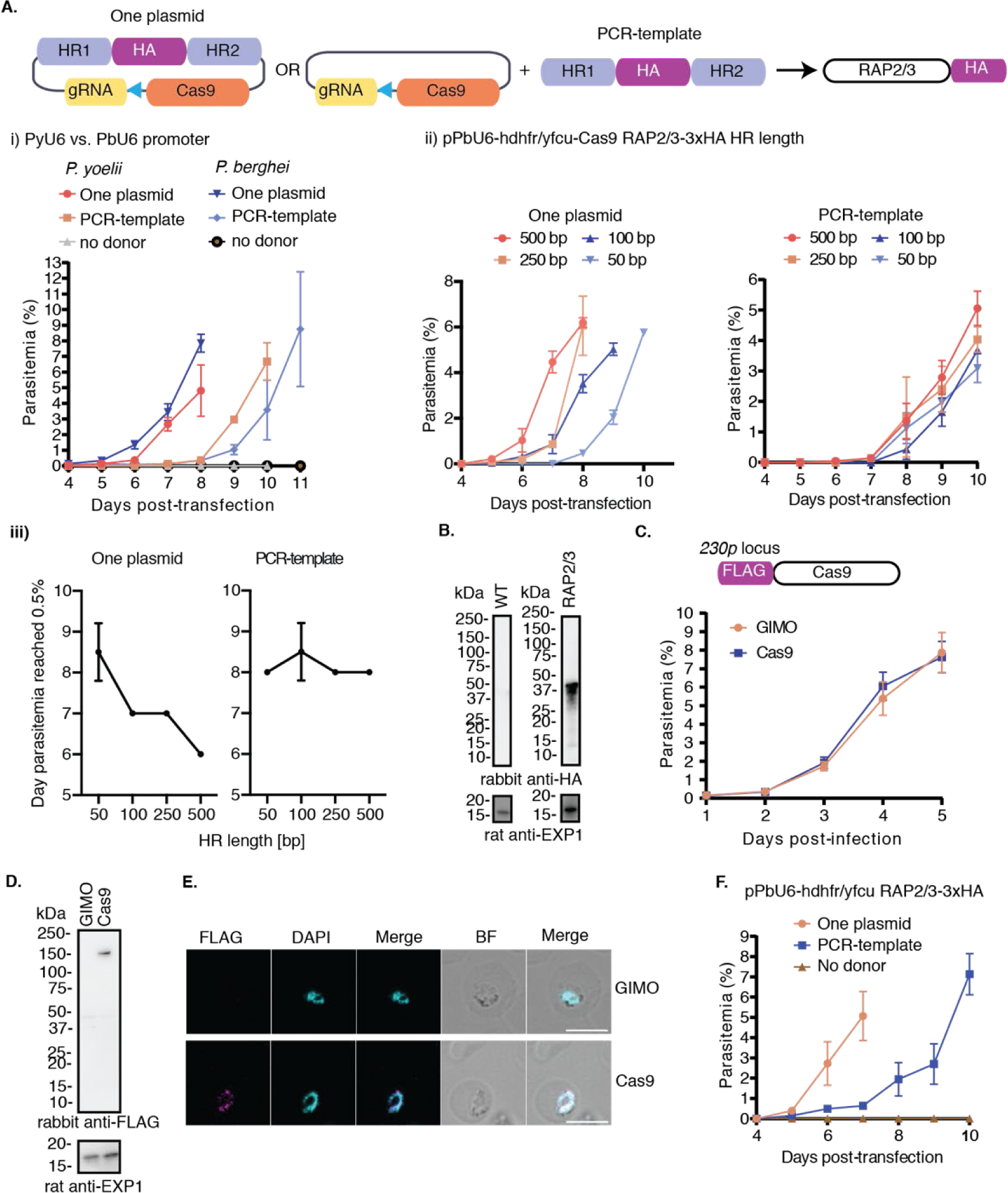
Characterisation of pPbU6-hdhfr/yfcu vector systems and the PbCas9 mother line. (**A**) Schematic of pPbU6-hdhfr/yfcu-Cas9 with the homology regions (HR) of the repair template delivered on the same plasmid (one-plasmid approach) or on a PCR product (PCR-template approach) to generate the *rap2/3*-3xHA parasite line. The effect on transfection efficiency was compared by both (**i**) substituting the *P. yoelii* U6 for the *P. berghei* U6 promoter (not significant) and (**ii**) different homology arm length (p = 0.0068). (**iii**) The relationship between homology arm length and transfection efficiency was examined by plotting homology arm length against the day parasitemia reached >0.5%. One or two-way ANOVAs were performed to compare transfection efficiencies from two biological replicates per condition. All parasitemia counts can be found in **Table S2**. (**B**) The expression of RAP2/3-3xHA was confirmed by Western blot following transfection with the pPbU6-hdhfr/yfcu-Cas9 vector using ∼500 bp HR. Wild type parasites were used as a negative control for the HA tag and the anti-EXP1 was used as a loading control. Full-length Western blots are shown in **Fig. S2**. (**C**) Growth assay of the PbCas9 line that constitutively expresses FLAG-Cas9 under the *hsp70* promoter with the PbGIMO mother line used as a control. Error bars = SD for three biological replicates. (**D**) FLAG-Cas9 expression was confirmed using Western blot, where FLAG detects Cas9 and anti-EXP1 was used as loading control. PbGIMO was used as negative control for the FLAG tag. (**E**) Immunofluorescence assay demonstrates that FLAG-Cas9 is expressed and localised to the nucleus. FLAG detects Cas9 and 4′,6-diamidino-2-phenylindole (DAPI) stains the nucleus. BF = Bright-field. The PbGIMO mother line was used as a negative control. Scale bar = 5 µm. (**F**) Transfection efficiency of the pPbU6-hdhfr/yfcu *rap2/3*-3xHA vector into the PbCas9 line was assessed using the one-plasmid versus the PCR-template approach. Error bars = SD for two biological replicates.

Next, we determined the minimum homology arm length required for efficient gene modification. We used the pPbU6-hdhfr/yfcu-Cas9 vector to target the *rap2/3* locus with both the one-plasmid and PCR-template approaches using varying length (∼50 to ∼500 bp) homology arms (**Fig. 1A ii**). Both the length of homology arms (p = 0.0068) and the use of a one-plasmid versus PCR-template approach (p = 0.0005) have a significant contribution to transfection efficiency, where the use of a one-plasmid system on its own is the most important determinant of efficiency. We further investigated the relationship between homology arm length and transfection efficacy in the two different systems and found that homology arm length has a significant effect on transfection efficiency for the one-plasmid (p = 0.0097) but not the PCR-template (p = 0.4789) approach (**Fig. 1A iii**). The most efficient editing was thus achieved using the one-plasmid approach with ∼500 bp homology arms, reaching a parasitemia of >0.5% on day six (**Fig. 1A ii-iii**). However, robust editing can be achieved with short ∼100 bp homology arms using both systems. Genotyping PCRs confirmed that the 3xHA epitope tag had been reliably integrated with both the one-plasmid and the PCR-template approach using homology arms from ∼100 to ∼500 bp. For very short ∼50 bp homology arms, the outcome was more variable (**Fig. S1B**). Western blot shows that RAP2/3-3xHA is expressed and of the expected size (**Fig. 1B**). Despite its lower efficacy, the PCR-template approach is attractive since it only requires cloning of the gRNA. We tested its broader applicability by tagging three other genes with 3xHA, steroid dehydrogenase (*sdg,* PBANKA_0522400), parasite-infected erythrocyte surface protein 1 (*piesp1,* PBANKA_0408500) and membrane associated histidine-rich protein 1a (*mahrp1a*, PBANKA_1145800) using ∼100 bp homology arms. The transgenic parasites came up on day 10 (*sdg*), 12 (*piesp1*) and 11 (*mahrp1a*) post-transfection. The correct integration and expression of the 3xHA tag was confirmed by PCR and Western blot. The expression of SDG and PIESP1 was also confirmed by immunofluorescence assays (IFA) (**Fig. S3**).

We hypothesised that endogenously expressing the Cas9 nuclease from the parasite’s genome would increase editing efficiency. We thereby generated a Cas9-expressing background line (PbCas9) in which Cas9 is under the control of the *P. berghei* constitutive promoter of heat shock protein 70 (*hsp70)*. The Cas9 expression cassette was inserted into the Gene In Marker Out (GIMO) locus within the dispensable *230p* gene of the GIMO *P. berghei* ANKA line^27^. Successful integration was confirmed by PCR (**Fig. S1C**) and did not affect parasite blood stage growth (**Fig. 1C**). Furthermore, Cas9 expression and expected nuclear localisation was confirmed by Western blot and IFA (**Fig. 1D-E**). We then generated a vector lacking Cas9, pPbU6-hdhfr/yfcu and evaluated it in combination with the PbCas9 parasite line. Again, we introduced a 3xHA tag into the 3’ of *rap2/3*. The Cas9 expressed from the genome in the new PbCas9 background line facilitated integration of the 3xHA tag using both the one-plasmid and PCR-template approaches (with ∼500 bp homology arms) and integration was confirmed by PCR (**Fig. S1D**). Consistent with previous observations, the one-plasmid approach was more efficient, with mice reaching a parasitemia above 0.5%, six days post-transfection (**Fig. 1F**).

### The pPbHiT vector containing the gRNA barcode and 100 bp homology arms efficiently integrates into the target locus

Scaling-up CRISPR-Cas9 in organisms lacking the c-NHEJ pathway requires the gene-specific gRNA and HR to be physically linked in the same plasmid to enable pooled transfections. Having established that the *P. berghei* genome can be effectively modified using short homology arms in a one-plasmid approach enabled us to adopt a high-throughput tagging strategy previously used in *T. gondii* ^25^. The CRISPR-Cas9 *P. berghei* High-Throughput strategy (PbHiT) developed here, relies on single-step cloning of a 320 bp synthetic fragment carrying the gRNA and homology arms into the pPbU6-hdhfr/yfcu-HiT (referred to as pPbHiT) vector, which we generated by modifying the pPbU6-hdhfr/yfcu vector (**Fig. 2**). Before transfection into Cas9-expressing parasites, the final pPbHiT vector containing the synthetic fragment is linearised, resulting in the homology arm sequences that drive integration flanking the entire plasmid. To reduce the likelihood of unintegrated plasmids maintained as episomes, the linearised vector was gel extracted prior to transfection (**Fig. 2**). When a Cas9-mediated DSB is repaired through HDR, the entire vector is inserted into the target locus and facilitates the editing of the target gene. The gene-specific gRNA is thereby stably integrated into the genome and serves as a molecular barcode to identify the edited parasites by NGS. The pPbHiT vector can be used for epitope tagging and gene knockout by adapting the position of the homology regions and gRNA (**Fig. 2)**. For epitope tagging, the endogenous 3’ untranslated region (UTR) of the target gene is replaced by the 3’UTR of the constitutively expressed gene *hsp*70. The pPbHiT vector has a modular design that enables easy modification to add or replace epitope tags or regulatory elements.

**Figure 2.**
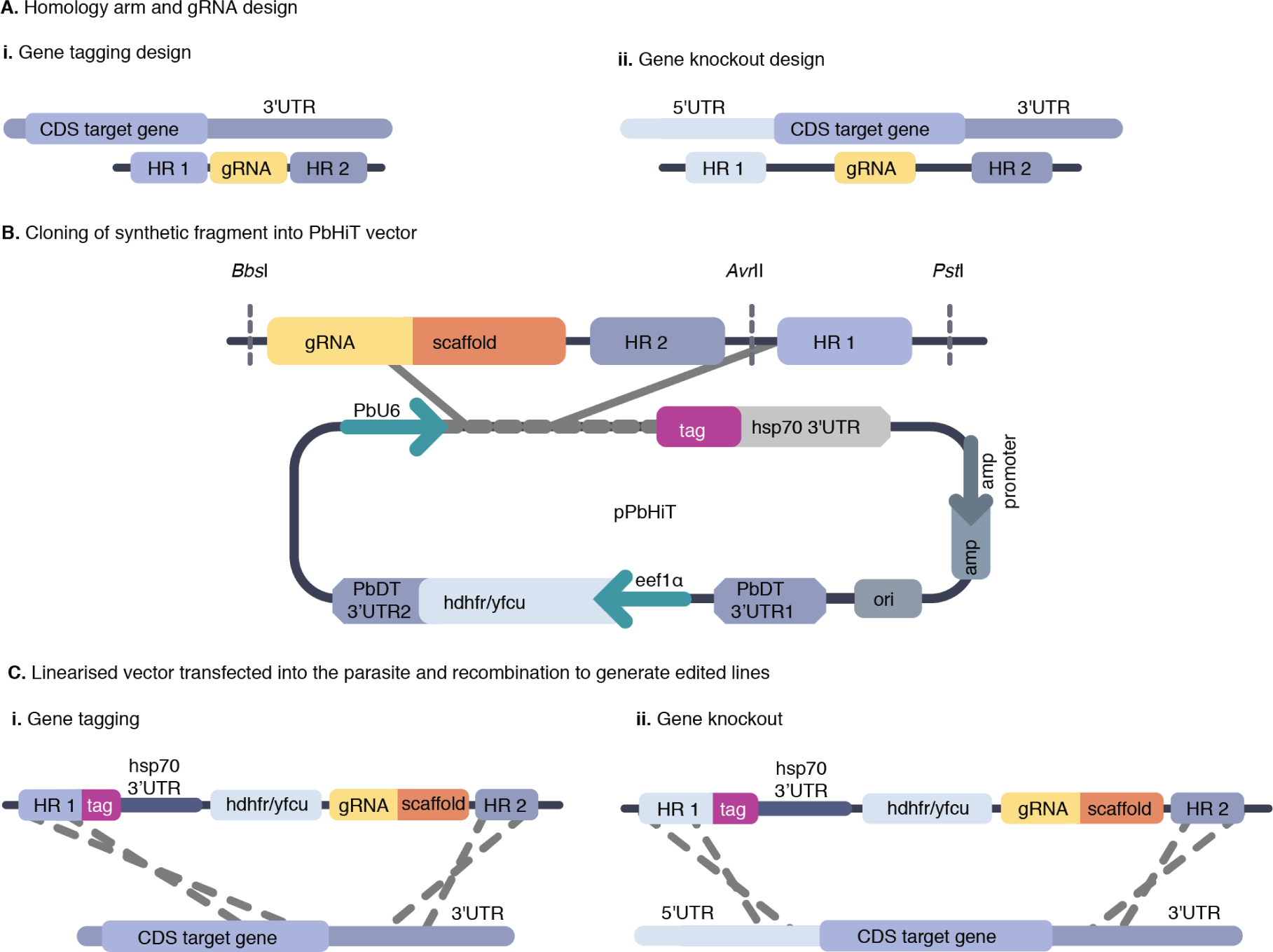
Schematic of pPbHiT vector design. (A) The placement of homology regions (HR) and gRNA in relation to the coding DNA sequence (CDS) of the target gene. (B) The gRNA, the gRNA scaffold, and the homology arms are ordered as a synthetic fragment in a pUC vector. The fragment is then cut out of the vector using *Bbs*I and *Pst*I restriction enzymes and cloned into the pPbHiT vector in a single step, where the pPbHiT vector has been linearised using *Bsm*BI and *Pst*I. The PbU6 promoter drives the gRNA expression. For tagging designs the 3’ end of the CDS ends up in-frame with a cMyc tag and the mRNA is stabilised by the *hsp70* 3’UTR. Expression of the positive-negative dual selection marker human dihydrofolate reductase (hdhfr) fused to yeast cytosine deaminase/uridyl phosphoribosyl transferase (*yfcu*) is driven by elongation factor 1 alpha (*eef1α)* promoter and is recyclable by the repeated terminator sequence of dhfr-thymidylate synthase (PbDT 3’UTR) flanking *hdfr-yfcu.* (C) The final pPbHiT vector with the synthetic fragment is linearised using *Avr*II before transfection, exposing the homology arms at each end of the vector to drive the integration into the target locus. The gene specific part of the gRNA then acts as a molecular barcode for the resulting mutant.

To test this strategy, we used the pPbHiT vector containing a triple cMyc (3x-cMyc) epitope tag and evaluated the efficiency of tagging genes using 50 and 100 bp homology arms. We targeted both *rap2/3* and PBANKA_1224200, a gene predicted to be localised along the secretory pathway. For both target genes, edited parasites were obtained with 50 and 100 bp long homology arms (**Fig. S4A**). Consistent with what we previously observed for the one-plasmid approach, 100 bp homology arms resulted in more efficient editing for both targets **(Fig. 3A i).** Furthermore, the wild type target loci were undetectable by PCR for both targets when using 100 bp homology arms **(Fig. S4A)**.

**Figure 3.**
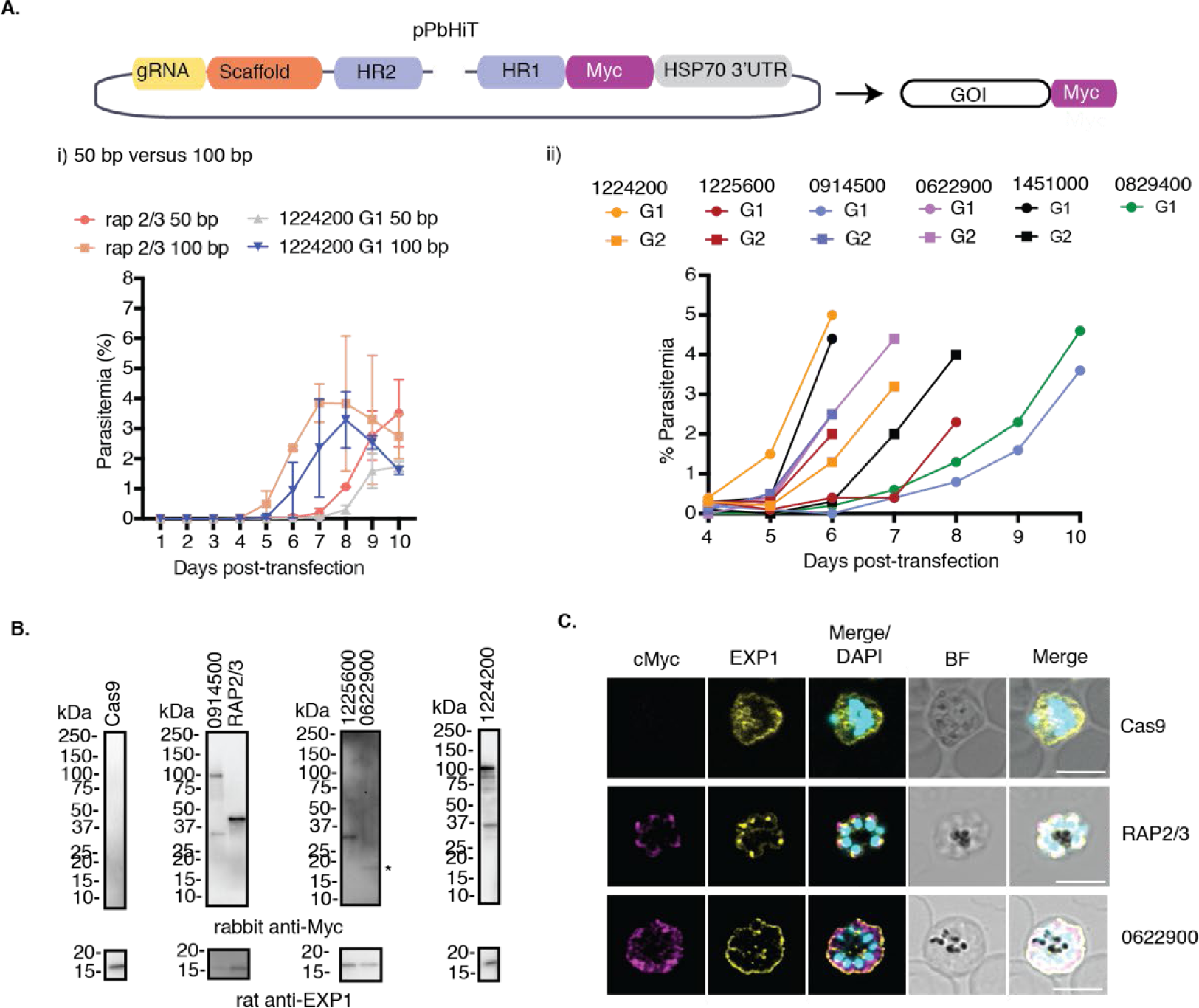
Efficient epitope tagging using the PbHiT system. (A) Schematic overview of pPbHiT plasmid used for cMyc epitope tagging of the gene of interest (GOI). (i) Daily parasitemia counts post-transfection comparing 50 bp and 100 bp homology arms for *rap2/3* and PBANKA_1224200 tagging vectors using a single gRNA. For both genes, the 100 bp homology region results in faster recovery of transgenic parasites. Error bars = SD for two biological replicates. (ii) Daily parasitemia counts post-transfection comparing transfection of six individual genes targeted (PBANKA_1224200, PBANKA_1225600, PBANKA_0914500, PBANKA_0622900, PBANKA_1451000 and PBANKA_0829400) using 100 bp homology arms and two gRNAs (G1 and G2) per target except for PBANKA_0829400. Both the gene targeted and the gRNA used affect transfection efficiency. All parasitemia counts can be found in Table S2. (B) The expression of tagged proteins was confirmed using Western blot using anti-HA antibody, where all proteins showed expected sizes. The PbCas9 background line was used as a negative control and anti-EXP1 was used as a loading control. Full-length Western blots are shown in Fig. S2. (C) Immunofluorescence assays of both RAP2/3-3xcMyc and PBANKA_0622900-3xcMyc showed that both proteins are expressed during the schizont stage as expected. Anti-EXP1 was used as a marker for the parasitophorous vacuole membrane/dense granules and DAPI to stain the nucleus. Scale bars = 5µm.

To further evaluate the performance of the PbHiT system, we tagged five other genes (PBANKA_1225600, PBANKA_0914500, PBANKA_0622900, PBANKA_1451000, and PBANKA_0829400) using 100 bp homology arms and two guides per gene. For PBANKA_0829400 we only obtained one gRNA within the accepted distance from the editing site, which is restricted to the 3’UTR for epitope tagging. Parasites emerged at different days post-transfection (days four to six), with a marked difference between the same gene targeted by the different gRNA, likely reflecting gRNA efficiency (**Fig. 3A ii**). However, this was not directly correlated to either the proximity of the guide to the editing site or the on/off gRNA target score. Genotyping PCRs confirmed the correct integration of the epitope tag in the 3’ UTR of the target gene (**Fig. S4B**). One gene (PBANKA_0914500) did not show any 3’ integration product when edited by guide one, however the 5’ integration was confirmed.

The expression of all but two of the tagged proteins (PBANKA_1451000 and PBANKA_0829400) was confirmed by Western blot for guide one (**Fig. 3B**). PBANKA_0914500-cMyc was detectable by Western blot, despite that it did not show a positive 3’ integration band by PCR. We also confirmed by IFAs that RAP2/3 and PBANKA_0622900 are expressed in schizonts, in agreement with transcriptomic data^28^. This shows that changing the gene’s 3’UTR does not affect mRNA stability and facilitates protein expression at the expected stage. In the case of RAP2/3, the 3x-cMyc tagged protein was observed in the apical end of the parasites, which is consistent with rhoptry localisation^29^ and demonstrates that the 3x-cMyc tag is not altering protein localisation (**Fig. 3C**). We also tested if transfections could be done without gel extracting the final pPbHiT linearised vector and saw no evidence of episomes by PCR (**Fig. S4C, D**). Guide RNA sequences and HR used for each tagging constructs can be found in **Table S3**.

In summary, the PbHiT system facilitates robust genetic modification of *P. berghei* using short 100 bp homology arms. Importantly, gene specific PbHiT vectors can be generated by a single cloning step that simultaneously introduces gRNA and HR sequences.

### PbHiT enables pooled vector transfections that recapitulate published knockout phenotypes

Having established the efficiency of the pPbHiT vector for editing single genes, we assessed the performance of the PbHiT system in pooled knockout vector transfections. We selected 12 target genes with *in vivo* blood-stage growth knockout phenotypes assigned with high confidence in the *Plasmo*GEM screen and classified as essential (n = 4), dispensable (n = 4), or slow growers (n = 4) (**Table S4**)^9^. The *rap2/3* gene was included since we have successfully modified it in single gene targeting experiments. Most genes were targeted by two gRNAs except for PBANKA_0515000 (ookinete surface protein P25, *p25*) and PBANKA_0933700 (mitogen-activated protein kinase 2, *map2k*), which had one guide each. A total of 22X knockout vectors were individually generated before pooling together in equal amounts, linearised, and transfected into the PbCas9 parasite line. Blood samples were taken at days four to eight post-transfection, genomic DNA was extracted and NGS sequencing libraries were prepared by nested PCR (**Fig. 4A**) (adapted from ^9^).

**Figure 4.**
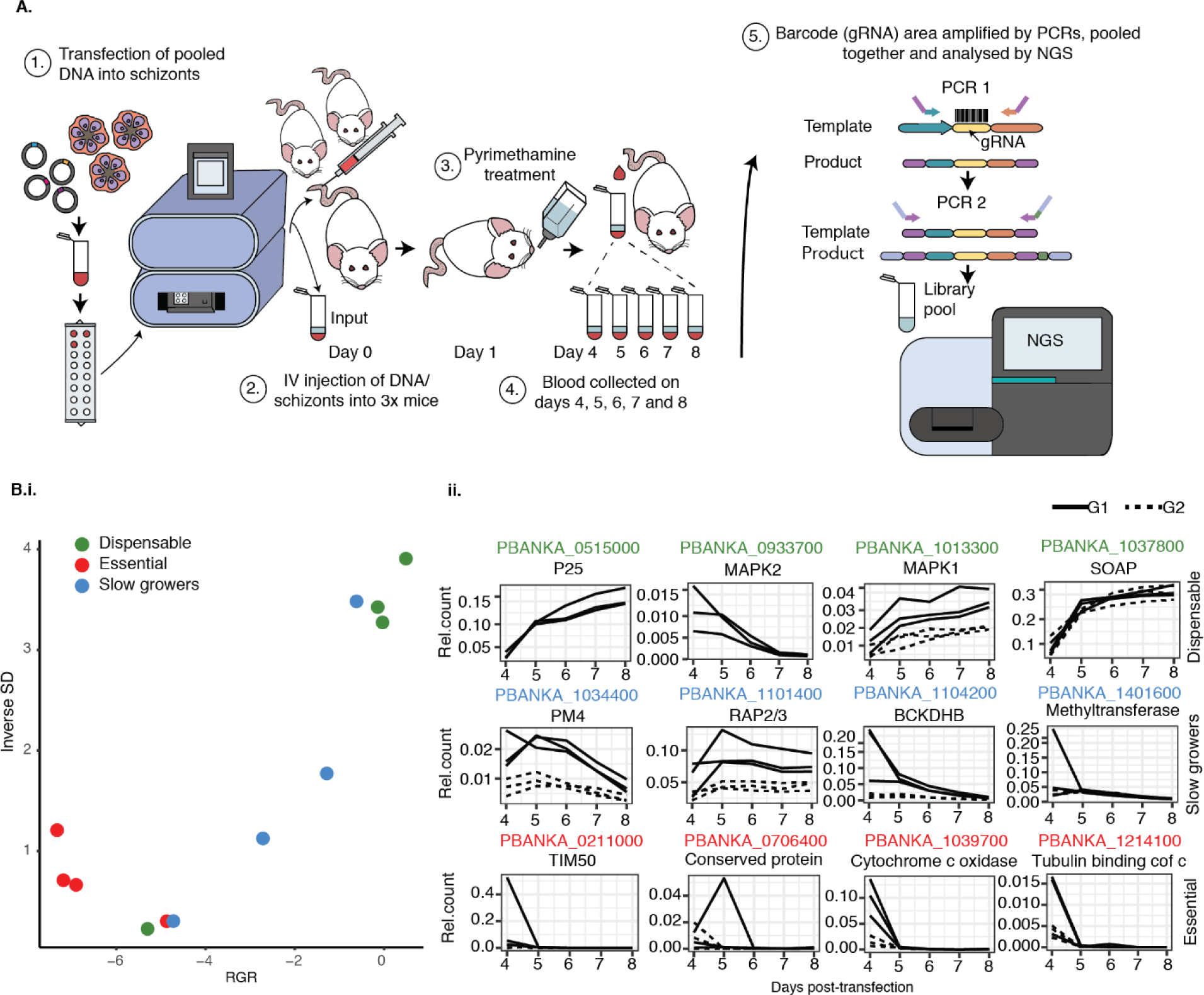
PbHiT pooled transfections recapitulate *Plasmo*GEM phenotypes. (**A**) Experimental workflow for *P. berghei* pooled transfection, sampling, and sequencing. (**B**) Results from pilot CRISPR screen of 22X knockout vectors targeting 12x genes with established blood-stage growth phenotypes. Relative mutant abundances are derived from gRNA sequencing and used to calculate mutant relative growth rates (RGR). (**i**) Scatter plot of mutant RGR, where genes are coloured according to previously published phenotypes^9^: dispensable (green), slow growers (blue) or essential (red). RGR is plotted against the inverse of standard deviation (1/SD) that is used as a robustness metric. (**ii**) Relative mutant abundance where each line represents counts from a single gRNA in an individual mouse. Experiment was completed in three biological replicates.

For analysis of gRNA sequencing data, the relative abundance of individual mutant gRNA serving as barcodes was calculated for each sample (representing one time point from a single mouse) as the ratio to the sum counts for all previously predicted dispensable genes present in the pool. A composite value for mutant relative growth rate (RGR) across all mice and days was then estimated by fitting the relative abundance to a logistic growth kinetic model, which takes into account growth rate, starting abundance, and growth rate saturation at later time points due to depletion of host reticulocytes. A standard deviation (SD) was calculated by assuming an error that is normally distributed, and 1/SD was subsequently used as a metric of robustness. Results revealed a high concordance between the PbHiT and *Plasmo*GEM data (**Fig. 4B**). Out of eight genes that were reported dispensable or slow growers in the *Plasmo*GEM screen, seven genes displayed the predicted phenotype. Furthermore, there was a clear structure in the data with separation between the dispensable genes from those with intermediate slow growth phenotypes. One predicted dispensable gene (PBANKA_0933700, *map2k*) instead had a phenotype typically associated with slow growing mutants, but only one gRNA was available for this gene, and it had low gRNA counts. All four essential mutants were rapidly lost from the pool (**Fig. 4B**). For dispensable genes (e.g. PBANKA_1037800 secreted ookinete adhesive protein, *soap*) the gRNA relative abundance is expected to increase or remain stable over time, while for genes with slow growth phenotypes (e.g. PBANKA_1034400 plasmepsin IV, *pm4*) the gRNA is typically gradually decreasing as it is outcompeted by dispensable mutants in the pool. In contrast, the gRNA relative abundance for essential genes (e.g. PBANKA_1214100, tubulin binding cofactor c) is expected to be quickly depleted (**Fig. 4B ii**). All CRISPR screen data presented in this manuscript is available at http://malaria-crispr2024.serve.scilifelab.se.

PCR indicated that unintegrated episomes, which also contain NGS-quantifiable gRNA barcodes, were absent from all samples taken post-transfection (**Fig. S4E**). However, gRNA barcode counts obtained on day four post-transfection are likely a combined signal from unintegrated and integrated plasmids. This is especially prominent when mutant abundance is very low, and an episomal gRNA signal is probably detectable by NGS despite not being visible by PCR (**Fig. S4E**). This can result in an artificially high gRNA abundance for essential genes on day four (**Fig. 4B ii**). Together, this indicates that PbHiT pooled transfections recapitulate published knockout phenotypes and can be used for CRISPR screens with a low rate of false positives and false negatives.

### PbHiT offers a scalable CRISPR system in *P. berghei*

To scale-up vector production and enable CRISPR screens, we established a pooled ligation protocol for generation of pPbHiT vectors. Briefly, glycerol stocks of pUC-GW-Kan vectors containing the synthetic fragment with gRNA and homology arm sequences were individually grown to saturation overnight in 96-well deep well blocks, and cultures were pooled prior to plasmid purification. The synthetic gene targeting fragments were released by restriction digest and then ligated into the pPbHiT vector. Pooled ligation reactions were transformed into bacteria, selected for on agar plates, and the resulting colonies scraped and pooled together for plasmid purification.

To determine the maximal insert pool size for the ligations, we tested pools of 8X, 12X and 22X inserts in three biological replicates. We used NGS to determine the number of gene-specific vectors recovered from each ligation pool. The optimal pool size, defined as the maximal number of inserts per ligation reaction versus the maximum recovery of individual gene-specific vectors, was determined to be 12X inserts with a recovery rate of 97.2% compared to 81.8% for a pool of 22X, with no measurable benefit of reducing the pool size to 8X inserts (91.7%). There does however appear to be a trade-off, using a smaller pool size of eight inserts can result in a more even proportion of each recovered vector **(Fig. S5 A, B).**

Pooling 12X inserts per ligation facilitates the generation of 96X vectors using only 8X individual ligation reactions. We used this approach to generate pools of pPbHiT knockout vectors for 24, 48, or 96 gene targets, each covered by two gRNAs, where vector pool composition was verified by NGS prior to transfection (**Fig. S5C**, **Fig. S6 and Table S5**). The constructs were automatically designed using a script that selects the highest scoring gRNAs and matching HR sequences for PbHiT mediated knockout of all *P. berghei* protein-coding genes. For gene knockout, the entire coding sequence is available for gRNA targeting and, as a result, close to 100% of protein coding *P. berghei* genes can be targeted by at least three guide RNA sequences. In contrast, for C-terminal tagging we are restricted to targeting the 3’UTR of the gene and editing efficiency decreases with distance of gRNA from the desired editing site. By allowing a gRNA search window spanning from the stop codon and 250 bp into the 3’UTR, 97% of *P. berghei* genes are targetable by at least one gRNA sequence (**Fig. S5D**).

### PbHiT facilitates pooled transfection CRISPR screens in *P. berghei*

The 96X gene targets selected for the PbHiT CRISPR screen included 26 genes with known blood-stage growth phenotypes^9^, with the remainder of the targets lacking knockout phenotypes (here classified as “unstudied”), (**Table S5**). All transfection pools were spiked with pPbHiT vectors generating three dispensable (*p25*, *soap*, mitogen-activated protein kinase (*map1k*)) and three slow-growing (*pm4*, methyltransferase and 2-oxoisovalerate dehydrogenase subunit beta (*bckdh-b*)) control mutants verified in the 22X vector pool experiment (**Fig. 4B, Tables S4 and S5**). Pooled transfections, sample collections, NGS library preparation, and analysis was performed as above (**Fig. 4A**). To test scalability, we transfected pools of 48X, 96X and 192X gRNAs where each gene was targeted by two gRNAs. The 48X pool contained 24 genes with known growth phenotypes^9^, and we show that even in a more complex mutant pool the PbHiT CRISPR screen data closely recapitulates *Plasmo*GEM phenotypes for the majority of genes (**Fig. 5A i-iii**). In addition, a good agreement between gRNAs is seen for most targets (**Fig. 5A ii**), although variation among targets and gRNAs exists which is reflected in relative mutant abundances (**Fig. S7**). Upon increasing the pool size to 96X gRNAs targeting 48 genes, the ability to capture known mutant growth phenotypes was retained, with good agreement between gRNAs (**Fig. 5B i-iii**). In addition, good reproducibility was observed for the 24 genes with known growth phenotypes that overlapped between the pools of 48X and 96X gRNAs (**Fig. 5C**), and both pool sizes performed very strongly (Area Under the Curve (AUC) = 0.99 and 0.97, respectively) when it comes to the ability to accurately call known phenotypes (**Fig. 5D**). To assess our ability to confidently assign new growth phenotypes, we combined the 48X and 96X vector pools and fitted a logistic regression model. This enabled the classification of 16 out of 22 previously unstudied genes as either essential (three) or dispensable (thirteen) to *P. berghei* asexual blood stage growth (**Fig. 5E**, **Table S6**). Most of the unstudied genes that were not assigned phenotypes using PbHiT had RGR values similar to those that were categorised as slow growing mutants in the *Plasmo*GEM screen^9^. Genes that were refractory to disruption with both PbHiT and *Plasmo*GEM vectors show good agreement with results from the *piggyBac* transposon mutagenesis screen in *P. falciparum*^30^. Finally, we provide evidence of dispensability for four unstudied genes (PBANKA_0103700, PBANKA_0812900, PBANKA_0409000, PBANKA_0821200) that were reported non-mutable in the *piggyBac* screen but here we were able to knockout using PbHiT (**Table S6**).

**Figure 5.**
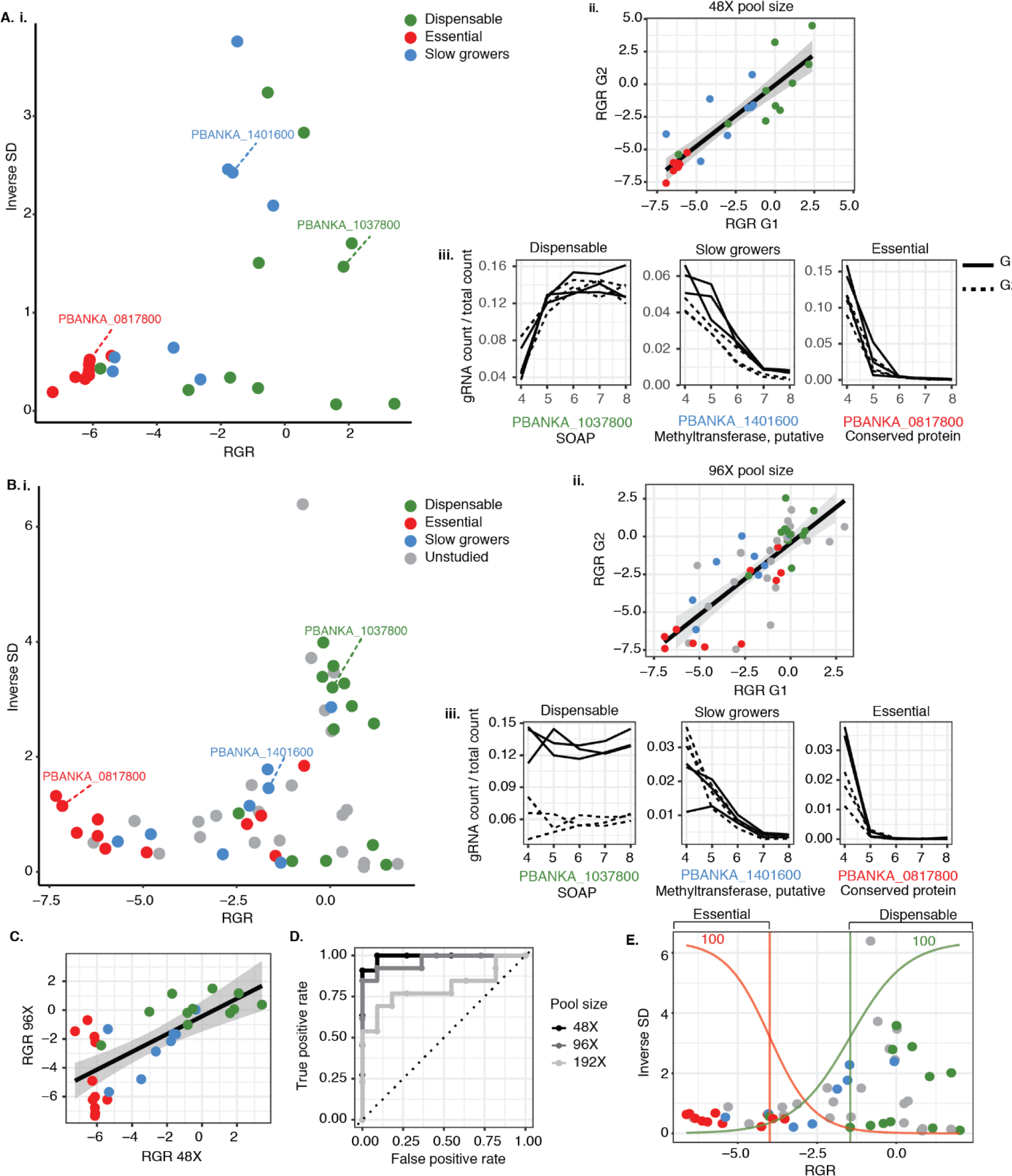
PbHiT CRISPR screen using pools of 48X and 96X vectors. (**A**) Analysis of 48X gRNA CRISPR screen targeting 24 genes with known blood-stage phenotypes. (**B**) Analysis of 96X gRNA CRISPR screen targeting 48 genes, with 24 target genes overlapping with the 48X gRNA pool. Panels (**i-iii**) for both **A** and **B** are displaying: (**i**) Scatter plot of mutant relative growth rates (RGR) plotted against the inverse of standard deviation (1 / SD). (**ii**) Correlation of RGR between two gRNAs targeting the same gene. (**iii**) Selected line graphs of mutant relative abundances, with each line representing counts from a single gRNA in an individual mouse. Relative mutant abundances for all genes in the 48X pool are available in **Fig. S7**. (**C**) Correlation between mutant RGR for genes overlapping between the 48X and 96X pool. (**D**) Receiver operator curve (ROC) that represents the ability to predict mutant phenotypes for 24 genes with known blood stage growth phenotypes within the 48X (AUC = 0.99), 96X (AUC = 0.97) and 192X (AUC = 0.8) gRNA pools. (**E**) By combining the 48X and 96X pools and fitting a logistic regression model we can assign phenotypes for unstudied genes. In all scatter and correlation plots the genes are coloured based on published blood stage growth phenotypes^9^: dispensable (green), slow (blue) or essential (red). Genes lacking *Plasmo*GEM knockout screen phenotype are classified as unstudied (grey). Results are from three biological replicates.

We further increased mutant pool size to that of 192X gRNAs targeting 96 genes, upon which the ability to accurately call known phenotypes was reduced (**Fig. S8A i**), but still with an AUC of 0.80 (**Fig. 5D)**. This was also reflected in a reduced correlation between gRNAs targeting the same gene (**Fig. S8A ii-iii**) and reduced reproducibility between experiments for genes overlapping between the pools of 48X and 192X gRNAs (**Fig. S8B**). Nevertheless, while RGR estimates appear less reliable for the pool of 192X gRNAs, it is likely that this larger pool size can still be useful to make broad calls of mutant essentiality and dispensability. We hereby show that transfection-ready pPbHiT vectors can be generated by a scalable single ligation step and up to 96X vectors can be pooled for parallel transfection of *P. berghei* schizonts thus enabling CRISPR screens.

## Discussion

We here examine the parameters required for efficient CRISPR-mediated gene editing of *P. berghei* and present the PbHiT CRISPR system, which allows robust gene modification using short (100 bp) homology arm vectors. The PbHiT vectors can be used to rapidly tag or knockout individual genes and can be made using a single cloning step protocol that can be scaled up by pooled ligations. We provide construct desings and sequences for homology region and gRNAs to knockout the entire *P. berghei* protein-coding genome. Finally, we demonstrate that PbHiT can be used in high-throughput CRISPR screens, where it accurately reproduces published blood stage growth knockout phenotypes.

The high efficacy of the PbHiT system is likely driven by a combination of: (1) linearisation of the pPbHiT vector to expose flanking homology arms to facilitate efficient ends-in double homologous recombination, which drives (2) stable integration of the *hdhfr/yfcu* drug selectable marker and finally, (3) the constitutive exogenous expression of Cas9 from within the *P. berghei* genome. Cas9 already being present in the cell has been suggested to increase efficiency since Cas9-mediated editing of the target gene can occur during the first replication cycle post-transfection^17^. In concordance with our results, *Plasmodium* is known to tolerate continuous exogenous expression of Cas9, and these parasites display normal blood stage fitness and can be transmitted by mosquitoes^17,19,31^. Interestingly, for the pPbU6-hdfr/yfcu plasmid, we saw no marked improvement in transfection efficacy using the PbCas9 line, this however allows us to reduce plasmid size for integration when using the PbHiT approach. Similarly, changing the *P. yoelii* to the *P. berghei* U6 promoter driving gRNA expression had little effect.

To date, *Plasmo*GEM knockout screen phenotypes are available for just over half of all *P. berghei* genes. The outstanding genes lack phenotypes since knockout vectors could not be generated. Furthermore, the *Plasmo*GEM C-terminal tagging vectors did not scale, only covering 13% of all *P. berghei* genes^32^. The PbHiT system offers a scalable method to rapidly produce vectors. PbHiT can be used to complete the knockout screen of the entire *P. berghei* genome and provide phenotypes of unstudied genes. PbHiT can also be used to close vector gaps for targeted screens surveying all genes with a predicted function, expression pattern, or subcellular localisation no longer restricting studies to genes for which *Plasmo*GEM vectors exist. We here used only two gRNAs per target gene to evaluate the feasibility of high-throughput CRISPR screens using PbHiT in *P. berghei*. Future screens aimed at assigning gene knockout or knockdown phenotypes *de novo* will likely require a minimum of three gRNAs per gene to further improve on accuracy and reproducibility^25^. Introduction of unique molecular identifier (UMI) sequences for gRNA barcoding could also be considered since it improves the statistics by tracking individual barcoded clones, aiding CRISPR screens with a reduced number of gRNAs^33^.

Forty-five percent of *P. berghei* genes are essential for blood stage growth^9^ and are therefore putative targets for novel antimalarials. However, the molecular function of essential genes cannot be readily examined since viable knockout mutants cannot be obtained. Robust conditional knockout or knockdown systems that can be scaled to a genome level are not yet available for *P. berghei* ^34^. We show that PbHiT can be used to effectively introduce C-terminal epitope tags to study protein localisation. However, the modular construction of the pPbHiT vector facilitates the addition of regulatory elements with conditional gene knockdown sequences to generate conditional alleles. This approach was used in a target screen of 147 kinases in *T. gondii*, using the original HiT vector system to introduce an auxin-inducible degron (AID) tag that facilitates inducible proteasomal degradation of the target protein^25^.

Efficient CRISPR-mediated editing using short homology arms is critical for cost-effective synthesis of gene-targeting sequences and its efficient cloning in a pooled format. We here systematically show the effect of homology arm length on editing efficiency, where robust editing was achieved using 100 bp homology arms. We expect the PbHiT system to be directly applicable to the closely related rodent malaria parasite *P. yoelii* and the zoonotic malaria parasite *P. knowlesi,* where short homology arms (80-100 and 50 bp, respectively) can be used in conjunction with CRISPR^16,35^. The human malaria parasite *P. falciparum* does not benefit from the same degree of genetic tractability as *P. berghei* and reverse genetic screens relying on pooled transfections have to date not been possible. CRISPR-Cas9 enhances gene editing efficacy in *P. falciparum* and its genome can be modified using relatively short homology arms of >200^14,36^. Combining linearisation of a drug-selectable double homologous integration vector, which contains the sequences for both gRNA and homology arms, with constitutive Cas9 expression will likely also benefit gene editing in *P. falciparum*^19^. The adaptation of PbHiT for mid-throughput protein localisation and conditional gene knockdown in *P. falciparum* should be explored and could be feasible using an arrayed instead of a pooled screening format as also demonstrated using the HiT system in *T. gondii*^25^ and the recently developed SHIFTiKO system in *P. falciparum*^37^. Should it be transferable to *P. falciparum*, the versatility of the PbHiT system would be a useful complement to the growing CRISPR-based toolkit available to query function in *P. falciparum*.

To conclude, we believe that PbHiT can become the vector system of choice for efficient CRISPR-based single gene editing in *P. berghei* and will take systematic functional gene annotation of its genome to the next level. The PbHiT system also has the potential to be adapted to other *Plasmodium* species as well as other species that lack c-NHEJ pathway, such as the agriculturally and medically important parasite species *Babesia* or *Cryptosporidium*.

## Methods

### Animal work

All animal work was done at Umeå University under Ethics Permit A34-2018 and A24-2023 approved by the Swedish Board of Agriculture (Jordbruksverket). Female BALB/c mice (purchased from Charles River Europe) from six weeks old were used for all routine parasite infections. Mice were housed in groups of four in individually ventilated cages with autoclaved wood chips and paper towels as nesting material, at 21 ± 1 °C under a 12:12 h light-dark cycle and relative humidity of 55 ± 5%. Female Wistar rats (Charles River Europe) >150 g were used for pooled vector transfections as they give rise to schizonts with more merozoites and higher transfection efficiency compared to schizonts raised in mice. Rat-derived schizonts were also used when a large number of transfections were performed simultaneously. Rats were housed in pairs in conditions equivalent to the ones described for mice. Animals were maintained under specific pathogen-free conditions and twice a year subjected to Exhaust Air Dust (EAD) monitoring and analysis. Animals were fed *ad libitum* with a commercial dry rodent diet. Fresh water was freely available at all times. The health of animals was monitored daily by routine visual health checks. To determine parasitemia of infected rodents, thin blood smears were prepared from a tail bleed, fixed by methanol and Giemsa-stained. Infected blood was harvested using heart puncture on animals under terminal anaesthesia (90 mg/kg Ketamine; 20 mg/kg Xylazine in PBS) and collecting blood into a syringe containing 100 µL of heparin (Sigma-Aldrich). Animals were euthanized through cervical dislocation.

### Molecular cloning

All primer and gRNA sequences are available in **Tables S1, S3, S4 and S5**. All vectors generated were verified by Sanger sequencing (Azenta Life Sciences) and full sequences for the vector backbones were deposited to Addgene (#216421 to #216423). Plasmid DNA was prepared for sequencing by GeneJET Plasmid Miniprep Kit (Thermo Fisher, USA).

#### Generation of the pGIMO-Cas9 plasmid

The pL1694 *230p*-targeting Gene In Marker Out (GIMO) plasmid was obtained from Chris J. Janse at Leiden University Medical Center^38^. In pL1694, mCherry is constitutively expressed under the control of the *P. berghei hsp70* 5’UTR (promoter) and 3’UTR (terminator). The mCherry gene was excised from pL1694 by *Bam*HI and *Not*I and replaced by a human codon-optimised spCas9 from pUF1-Cas9^12^ by Gibson cloning (NEBuilder HiFi DNA Assembly Master Mix, New England Biolabs, NEB) to generate pL1694-GIMO-Cas9.

#### Construction of new P. berghei CRISPR-Cas9 vector backbones

To generate a CRISPR-Cas9 system specifically adapted for *P. berghei* we modified the *P. yoelii* pYCm and pYCs plasmids (kind gift from Jing Yuan^26,31^) by changing the *P. yoelii* U6 promoter (PyU6) that drives gRNA expression, for the *P. berghei* U6 promoter (PbU6; PBANKA_1354380). To this end, the PbU6 sequence was amplified from *P. berghei* ANKA cl15cy1 genomic DNA and cloned into the pYCs and pYCm plasmids following digestion with *Kas*I and *Stu*I (NEB), using the NEBuilder HiFi DNA reaction master mix. The resultant vectors were named pPbU6-hdhfr/yfcu-Cas9 (addgene ID 216423, derived from pYCm and containing the coding sequence for spCas9 nuclease) and pPbU6-hdhfr/yfcu (Addgene #216422, derived from pYCs). Both vectors contain the dual selection marker *hdhfr/yfcu* for positive selection with pyrimethamine and negative selection using 5-fluorocytosine (5-FC).

To adapt this system for ease of cloning and pooled transfections, we generated a vector using the pPbU6-hdhfr/yfcu backbone in which the original gRNA scaffold sequence was replaced with a synthetic modular fragment carrying the following features: *Bsm*BI/*Pst*I/3x-cMyc/*Sal*I/*Not*I/stop codon (TAG)/*hsp70* 3’UTR/*Aat*II (ordered from GeneWiz-Azenta). The pPbU6-hdhfr/yfcu plasmid and the synthetic fragment were digested with *Bsm*BI and *Aat*II (NEB) and ligated into the vector with T4 ligase (NEB). The multiple cloning sites allow adding new or replacing all features. The resulting plasmid was named pPbU6-hdhfr/yfcu-HiT (Addgene #216421), referred to as pPbHiT.

#### Generation of gene-specific P. berghei CRISPR-Cas9 targeting vectors

The gRNAs were designed using the EuPaGDT tool^39^. The selection of the best guides was made considering (i) the total score given by the tool, (ii) the proximity to the editing site, and (iii) the off-target score (based on the *P. berghei* ANKA genome, version PlasmoDB-28). Coding sequences of the *P. berghei* target gene along with 5’ and 3’UTRs were retrieved from PlasmoDB^40^.

Vectors to target specific genes using pPbU6-hdhfr/yfcu-Cas9 or pPbU6-hdhfr/yfcu were generated by first cloning the gRNAs into *Bsm*BI-digested vectors. To this end, two single-stranded oligonucleotides (Integrated DNA Technologies, IDT) were designed containing the guide sequence fused to a 4-nucleotide sequence (TATT for the forward guide and AAAC for the reverse guide) corresponding to the overhangs generated when digesting the vectors with *Bsm*BI. Single-stranded oligonucleotides were mixed in a 1:1 ratio, phosphorylated using the T4 polynucleotide kinase enzyme (NEB), and annealed by incubating at 95 °C for 5 min followed by a temperature ramp of −5 °C every minute, until reaching 25 °C. A 1:200 dilution of the double-stranded gRNA was ligated into *Bsm*BI-digested plasmids using T4 ligase (NEB). One µL of the ligated vector was then transformed into chemically competent XL Gold *E. coli* (Agilent Technologies). Integration was determined by colony PCR using the gRNA forward oligonucleotide and the generic primer gRNAseq_R. The HDR templates were synthesised by GeneWiz-Azenta. To generate tagging vectors, the homology arms were designed flanking the cut-site of the gRNA, which was placed in the 3’ end of the coding sequence. Furthermore, the area containing the gRNA target sequence was recodonised to avoid successive cutting of the edited locus, and the desired epitope tag was added to the repair template. To test the effect of different homology arm lengths, vectors for 3xHA tagging of *rap2/3* were prepared with the following homology arm lengths: ∼50 bp (5’HR: 65 bp, 3’HR: 50bp), ∼100 (5’HR: 130 bp, 3’HR: 126 bp), ∼250 bp (5’HR: 268 bp, 3’HR: 281 bp), ∼500 (5’HR: 582 bp, 3’HR: 545 bp). The HDR template was provided either in the same plasmid that carried the gRNA (one-plasmid approach), or as a PCR product (PCR-template approach). For the one-plasmid approach, the gene-specific homology repair template was ligated into plasmids carrying the corresponding gRNA using *Hind*III. For the PCR-template approach, the HDR template was amplified by PCR and the amplicon was incubated with *Dpn*I or gel extracted. For *rap2/3* only one guide was used per transfection but for *sdg*, *piesp1* and *mahrp1a* two guides were mixed together with PCR-template prior to transfection and recodonised area in the HDR template covered the region of both gRNAs.

The pPbHiT vectors to either knock out or tag target genes were designed with 50 or 100 bp homology arms, and the gRNAs were located in the region between the two homology arms, which results in the removal of the target site after recombination has occurred. For pPbHiT tagging vectors, the 5’ homology arm (HR1) is located at the end of the coding sequence and comprises the region immediately upstream of the stop codon without including it, whereas the 3’ homology arm (HR2) starts after the Cas9 cut site, 6 bp downstream of the gRNA protospacer adjacent motif (PAM) site. Guides within 50 bp from the end of the coding sequence are prioritised for tagging vectors to maximise editing efficiency. For pPbHiT knockout vectors, the HR1 is located in the 5’ UTR of the target gene immediately before the ATG codon, and the HR2 is designed in the 3’ UTR of the target gene just after the stop codon. All gene-specific elements and the generic gRNA scaffold were synthesised as a single synthetic fragment (Genewiz-Azenta) according to the following structure: *Bbs*I/gRNA/Scaffold/HR2/*Avr*II/HR1/*Pst*I for cloning into pPbHiT. If any of the cut sites were present in the sequence of the HR, the nucleotides were modified by introducing a silent mutation to remove the enzyme recognition site. The synthetic constructs were then ligated into the pPbHiT vector using the *Bbs*I and *Pst*I cloning sites. This places gRNA expression under the PbU6 promoter of the pPbHiT vector, and for tagging vectors the HR1 (corresponds to the 3’ end of the coding sequence) in-frame with the 3x-cMyc tag followed by the *hsp70* 3’UTR. Ligations were transformed into XL Gold *E. coli* as above and colonies were screened to check the presence of the insert by colony PCR using PbU6prom_F and hsp70UTR_R primers, except for the pooled vectors which were screened by NGS. The final pPbHiT vectors were linearised using the *Avr*II restriction enzyme prior to transfection.

#### Pooled pPbHIT vector ligation protocol for CRISPR screens

A protocol for pooled ligations was established to scale-up production of pPbHiT vectors. To this end, different ligation pool sizes were tested (8X, 12X and 22X inserts) in three biological replicates. Minipreps from individual pUC-GW-Kan knockout vectors were pooled together in pools of 8X, 12X or 22X (total 3 µg per pool) and digested with *Pst*I and *Bbs*I overnight at 37 °C and the pooled synthetic fragment was gel extracted and purified. The pool of fragments were then ligated into the pPbHiT vector (linerised with *Bsm*BI and *Pst*I), transformed and selected on Luria broth agar plates containing 100 µg/mL ampicillin. Colonies were counted, scraped together and incubated in Luria broth with ampicillin overnight at 37 °C shaking at 150 rpm. Plasmid vector pools were purified by Plasmid Midiprep (Qiagen) and prepared for Illumina sequencing using nested PCRs (described in detail below). The optimal ligation pool size was determined by colony counts and Illumina sequencing by looking at individual gRNA representation and diversity in each pool.

For the pilot pool of 22X knockout vectors the pPbHiT vectors were prepared individually and confirmed by Sanger sequencing. Then, 680 ng of each vector was combined (15 µg in total) and linearised overnight with *Avr*II and vector gel purified and precipitated. The final DNA pellet was resuspended in 30 µL water and then used for transfection and injected into three mice. For parasite transfection of pools with 24X, 96X or 192X vectors, glycerol stocks of individual Genewiz pUC-GW-Kan constructs were grown in 1 mL Luria broth with kanamycin (50 µg/mL) in a 96X deepwell plate overnight at 37 °C, shaking and next day pooled together in pools of 12 (with each row on the plate constituting one pool) and plasmid DNA was purified using Midiprep. Three µg of each DNA pool was then digested overnight using *Bbs*I and *Pst*I and fragment pools gel extracted, purified, and ligated into pPbHiT as described above. For transfection, roughly 450 ng of each vector (5,400 ng per pool of 12) was combined to make up pools of 48X, 96X, and 192X vectors. Each pool was also spiked with control vectors (450 ng per vector) (**Tables S4 and S5**). Pools were linearised overnight using *Avr*II, gel extracted, and precipitated for transfection. The amount of DNA for each transfection pool was, therefore, ∼13.6 µg (48X + controls), ∼48.150 µg (96X + controls) and ∼91.350 µg (192X + controls) prior to gel extraction. All DNA pools were resuspended in 30 µL of water prior to transfection as before and injected into three mice.

#### Generation of automated pPbHiT vector design tool

To generate resource of gRNA and HR sequences for the knockout and tagging of all predicted *P. berghei* genes, a script was made where users can enter the PBANKA ID and the script generates synthetic fragments for all available gRNAs for knockout vectors. The script assembles all the components of the pHiT synthetic fragment: *Bbs*I/gRNA/Scaffold/HR2/*Avr*II/HR1/*Pst*I and provides a csv file of the oligo sequence ready to be ordered. Guide RNA sequences for all *P. berghei* genes were obtained from the EuPaGDT^39^, which scores and ranks gRNAs based on the guide efficiency, on and off target effects and microhomology arms flanking the gRNA sequence. For the homology arms of the knockout strategy, 100 bases upstream of the start codon and 100 bases after the stop codon for each gene were identified using bedtools on the PlasmoDB *P. berghei* genome release 57 (57_PbergheiANKA). The code for knockout designs can be accessed on GitHub (https://github.com/srchernandez/PbHiT-CRISPR-Bushell-Lab) and Excel file for the top three ranking guides can be provided by request.

#### Polymerase chain reactions

For standard PCR amplifications (vector confirmation and genotyping), either GoTaq G2 Green Master Mix (2X, Promega, USA) or DreamTaq Green PCR Master Mix (2X, Thermo Scientific) were used. For GoTaq, generic cycling conditions included an initial 2 min denaturation at 95 °C, followed by 30 cycles, which consisted of denaturation at 95 °C for 15 s, annealing for 30 s at 2 °C below the calculated melting temperature of the primers, and elongation at 62 °C for 1 minute for every 1 kb, followed by a final extension step of 62 °C for 5 min. When DreamTaq was used, cycling conditions included an initial 2 min denaturation at 95 °C, followed by 30 cycles, which consisted of denaturation at 95 °C for 30 s, annealing for 30 s at 5 °C below the calculated melting temperature of the primers and elongation at 62 °C for 1 minute for every 1 kb, followed by a final extension step of 62 °C for 5 min. All PCR amplicons were analysed using agarose gel electrophoresis stained with Sybr Safe (Invitrogen) or ethidium bromide. The list of primers and their nucleotide sequence can be found in **Table S1** and full-length agarose gels in **Fig S3**, **S9** and **S10**.

#### Parasite transfections, sample collection and drug selection

Transgenic *P. berghei* parasite lines were generated either in *P. berghei* ANKA cl15cy1 wild type strain or in the PbCas9 mother line produced through this work. For parasite transfections, schizonts were produced using infected blood from mice (single gene transfections) or female Wistar rats (pooled transfections or single gene transfections where a large number of schizonts are required). Parasites were cultured in complete media (RPMI 1640 (Gibco), 25% FBS (Gibco), penicillin-streptomycin (100 U/mL and 100 µg/mL, respectively; Gibco) and 24 mM NaHCO3) for 22 h in flasks gassed with a 3% CO2, 1% O2, 96% N2 gas mixture, at 37 °C and shaking at 80 rpm. Schizonts were isolated on a 15.2% Histodenz / PBS (Sigma Aldrich) cushion and washed in complete media. Schizonts were transfected by electroporation using the Lonza 4D Nucleofector System according to the pulse program FI-115 with the P3 Primary Cell 4D-Nucleofector X Kit S (Lonza). After transfection, the parasites were resuspended in 100 µL RPMI and immediately injected intravenously into a mouse via the lateral caudal vein. The selection of resistant transgenic parasites was achieved by administering the appropriate drug in drinking water starting one day after transfection (considered as day 1 of transfection).

Parasitemia was routinely monitored by Giemsa-stained smears from the tail. For single gene transfections, infected blood was collected by cardiac puncture to prepare frozen stocks and for genomic DNA extraction. Parasite genomic DNA was extracted from 50 µL of infected blood mixed with 150 µL PBS using the DNeasy Blood and Tissue kit (Qiagen). For pooled transfections, an input sample was collected by washing out the electroporation cuvette with 200 µL PBS with samples boiled at 95 °C for 5 min and spun down at 16,200 x g and frozen at −20 °C. Twenty µL blood samples were collected by tail bleed into a tip containing small amount of heparin from day four and until day eight post-transfection, and placed into 200 µL PBS containing 10 µL heparin. Samples were kept at 4 °C until the last collection time point. Samples were then spun down at 16,200 x g for 1 min, PBS removed and pellets frozen at - 20 °C until DNA was extracted using the DNeasy Blood and Tissue kit.

When pPbU6-hdhfr/yfcu-Cas9 or pPbU6-hdhfr/yfcu were used, pyrimethamine (0.07 mg/mL, MP Biomedicals) was removed at day five post-transfection. When pPbHiT was used, pyrimethamine selection was maintained throughout the experiment since the *hdhfr/yfcu* marker stably integrates. To generate the *P. berghei* Cas9 (PbCas9) mother line, the transgenic parasites were selected using negative selection by 5-FC (Sigma, USA; 1 mg/mL). Parasites were cloned by limiting dilution.

#### Growth assays and statistical analysis of transfection efficiency

Daily tail bleeds (from day three or four post-transfection) were performed to generate Giemsa-stained smears and calculate parasitemia to determine transfection efficiency. A total of 1,000 red blood cells were counted using a microscope under 100x magnification, and parasitemia was calculated as the percentage of infected red blood cells. A two-way ANOVA with a confidence level of 99% was performed to compare the transfection efficiencies when using the *P. berghei* and *P. yoelii* U6 promoters in the one-plasmid and PCR-template approaches, as well as the length of the homology arms. Additionally, one-way ANOVAs were performed to compare the transfection efficiencies obtained with homology arms of different lengths in the separate approaches. The test was performed using GraphPad Prism version 10.0.0 for Windows, GraphPad Software, Boston, Massachusetts, USA.

#### Illumina libraries and sequencing

The barcode region (gRNA sequences) were amplified from 5 µL of gDNA by PCR using BC_pHiT_illumina_F and BC_pHiT_illumina_R primers, resulting in a 250 bp amplicon. Advantage Taq polymerase (Takara) was used with an initial 5 min denaturation at 95 °C, followed by 35 cycles, which consisted of denaturation at 95 °C for 30 s, annealing for 20 s at 55 °C and 68 °C extension for 8 s, followed by a final extension step of 68 °C for 10 min. The resulting 5 µl PCR product was used as a template for the second PCR using generic primer PE 1.0 and sample-specific index primers, which facilitates sample multiplexing and incorporates Illumina adaptor sequences. The cycling conditions for the second PCR consisted of initial 2 min denaturation at 95 °C, followed by 10 cycles, which consisted of denaturation at 95 °C for 30 s and 68 °C extension for 15 s, followed by a final extension step of 68 °C for 5 min. The nucleotide sequence of all primers can be found in **Table S1**. The resulting sequencing libraries were quality controlled by standard gel electrophoresis, purified by MinElute 96 UF PCR Purification Kit (Qiagen), eluted in 50 µL pure water and quantified using the Qubit dsDNA broad range (BR) assay (Thermo Fisher Scientific). One hundred twenty-five ng of each sample was pooled prior to quality control and quantification on the 2100 Bioanalyzer system using DNA 7500 kit (Agilent) and KAPA Library Quantification Kit Illumina/Universal (Roche), and finally diluted to 1.5 pM before loading at a low cluster density (4 x 10^5^ clusters / mm^2^) together with 50% of PhiX spike-in (Illumina). For libraries sequenced on NextSeq extra purification and removal of adapter dimers was performed using SPRIselect beads (Beckman Coulter) in a 1: 0.8 library: beads ratio. These libraries were diluted to 0.5 pM and loaded at low cluster density (7 x 10^4^ clusters / mm^2^). The low density and high proportion of PhiX prevent index hopping when sequencing low complexity libraries like these, where only the 20 bp gRNA of each amplicon is unique. The amplified gRNA libraries were sequenced using Illumina MiSeq Reagent Kit v2 (2×150 bp) on the Illumina MiSeq platform at SciLifeLab National Genomics Infrastructure (ligation pools and 22X knockout vector pool) or NextSeq 500/550 v2.5 Mid output (2×75 bp) kit on the Illumina NextSeq platform at Umeå University Hospital (48X, 96X or 192X vector pools).

#### Growth rate statistical analysis for CRISPR screens

A custom R script extracted gRNA abundances by finding the U6 sequence CAATATTATT and parsing the following target sequence. Only gRNA sequences exactly matching the expected list were retained. For each mouse and time point, the fraction of gRNA abundances were calculated. Relative abundances were obtained by dividing to the sum of control (non-essential) genes. A function representing logistic growth was fitted:

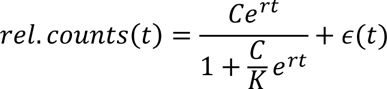

To ensure robust fits, we chose the carrying capacity as *K* = max (rel. counts). The relative growth rate *r*, and the initial abundance *C*, remained to be fitted. The error ε is assumed to be normally distributed, allowing the fit to be performed by a nonlinear least squares solver. The R package minpack.lm v.1.2-4 was used for this purpose. We also tested models with ε being log-normally distributed, but these yielded poor fits when the abundance was low, and thus the number of counts (reads) being low and uncertain. As the starting point for the Levenberg-Marquardt algorithm, we set *r* = 0 and C = E[count]. ε is assumed to be normally distributed, and we found that performing the fit in linear rather than log space greatly improved the fit for essential genes, which frequently had low counts and were difficult to sample at late time points. In case nls() did not converge, the result was excluded. This mainly happens when the mutant coverage is low and variance thus high, and effectively this functions as an outlier removal algorithm. Growth rate of genes were taken to simply be the average over the corresponding gRNAs. Variances were propagated based on Var[X ± Y] = Var[X] + Var[Y], if Χ and Y are independent variables. RShiny, ggplot2 and plotly were used to visualise the results. ROC curves were generated using the plotROC R package.

#### Immunoblot assays

To obtain samples for immunoblot assays, BALB/c mice were infected intraperitoneally with 200 µL of frozen parasite stock. When parasitemia reached around 5%, mice were bled as mentioned previously and put into culture. Mixed late-stage parasites were harvested using a 67% Percoll (RPMI and PBS) density gradient media (Cytiva). The Percoll purified pellet was washed 2x times in PBS containing protease inhibitors (cOmplete, Roche) and frozen at −80 °C until used. The pellet was either thawed on ice and resuspended in a 20x pellet volume of 1X Laemmli buffer (Bio-Rad) or lysed in 20x pellet volume of 1% Triton X-100 and freeze/thawed 2x and spun down at 16,000 x g for 10 min at 4 °C and supernatant resuspended in Laemmli buffer prior to Western blot. All samples were reduced in 100 mM DTT (Thermo Fisher) for 10 min at 80 °C. Samples were loaded on 4-20% TGX gels (Bio-Rad) and run in 1X TGX running buffer at 150 V for 50 min. Proteins were transferred onto a PVDF membrane (Bio-Rad) using a semi-dry transfer system (Bio-Rad) at 20 v for 7 min. The membrane was blocked in 1% casein in PBS for 1 h and then incubated with primary antibodies (in blocking buffer) overnight at 4 °C. The membrane was washed three times in PBS and subsequently incubated with HRP-conjugated secondary antibodies for 1 h and washed as before. The membrane was then exposed to a chemiluminescence substrate (Millipore) and protein bands were visualised using the Chemidoc imaging system (Bio-Rad). Primary antibodies used: rabbit anti-FLAG (Sigma-Aldrich #SAB4301135, 1:1,000), rabbit anti-cMyc (Cell Signal #2278, 1:1,000), rabbit anti-HA (Cell Signal #C29F4, 1:1,000) and rat anti-EXP1 (custom made from Proteogenix, 1:1,000). Secondary antibodies used: anti-rabbit IgG (H+L) HRP conjugate (Promega W4011, 1:2,500); anti-mouse IgG (H+L) HRP conjugate (Promega W4021, 1:2,500) and anti-rat IgG (H+L) HRP conjugate (Invitrogen #31470 1:10,000).

#### Immunofluorescence assays

The localisation of proteins was examined by IFA using a protocol adapted from Tonkin *et al.* 2004^41^. Briefly, infected red blood cells were placed onto poly-D-lysine coated glass slides and fixed with 4% paraformaldehyde and 0.0075% glutaraldehyde in PBS for 20 min at room temperature (RT). The fixed cells were permeabilised and quenched with 0.1 % Triton X-100 and 0.1 M glycine in PBS for 15 min at RT and further blocked with 3% BSA in PBS for 1 h at RT. Cells were then incubated with primary antibodies overnight at 4 °C. Antibodies were washed off and cells were subsequently incubated with secondary antibodies for 1 h at RT in the dark^42^. Cover slips were then mounted with an anti-fade medium containing DAPI (VECTASHIELD Plus; Vector laboratories) and sealed with nail polish. Images were acquired using an inverted confocal microscope (SP8; Leica) with a 63x oil-immersion objective lens. Primary antibodies used and concentration: rabbit anti-cMyc (Cell Signal #2278, 1:250), rabbit anti-HA (Cell Signal cat#C29F4, 1:250), rabbit anti-FLAG (Sigma-Aldrich #SAB4301135, 1:250), rat anti-EXP1 (1:500). Secondary antibodies used and concentration: Alexa fluor 594-conjugated goat anti-rabbit IgG (Invitrogen A11012, 1:2,000), Alexa fluor 647-conjugated goat anti-rabbit IgG (Invitrogen A32733, 1:1,000), Alexa fluor 594-conjugated goat anti-rat IgG (Invitrogen A48264, 1:1,000). All images were analysed using ImageJ/Fiji^43^.

### Figures

Schematics were made using Affinity designer (Fig. 2 diagram) or Adobe Illustrator (all other diagrams). All figures were arranged together using Adobe Illustrator.

## Data availability

The analysed CRISPR screen data is available and visualised using R Shiny (http://malaria-crispr2024.serve.scilifelab.se). The scripts to extract counts from FASTQ, analyse and visualise the results are available on GitHub (https://github.com/henriksson-lab/malaria_crispr2024). The code for knockout designs can be accessed on GitHub (https://github.com/srchernandez/PbHiT-CRISPR-Bushell-Lab) and an Excel file containing the pPbHiT synthetic fragment design with gRNA and HR sequences for the three top ranked gRNAs for the entire *P. berghei* protein-coding genome for the knockout strategy is available upon request.

## Reagent availability

Tool kit plasmids are available from Addgene and all gene specific vectors are available upon request.

## Acknowledgements

We acknowledge the Biochemical Imaging Center at Umeå University and the National Microscopy Infrastructure, NMI (VR-RFI 2019-00217) for providing assistance in microscopy. We thank Jing Yuan, Xiamen University for the kind gift of the pYCs and pYCm plasmids and Colin Herd for construction of the pGIMO-Cas9 vector. Thanks to Sebastian Lourido, Whitehead Institute and Massachusetts Institute of Technology, for input during conception of the PbHiT system.

## Author Contributions

T.K.J., M.S.P. and T.I. planned experiments and analysed data. T.K.J., M.S.P., T.I., M.I., S.H., A.H.C, P.S. and M.S. conducted experiments. S.H. and D.D. generated the automated CRISPR design tool. J.H. designed and performed the CRISPR screen analysis. T.K.J., M.S.P. and E.S.C.B wrote the original draft. T.K.J. and S.H. generated the figures. E.S.C.B. conceptualised the study. E.S.C.B. and J.H. supervised. All authors participated in the review and editing of the final manuscript.

## References

1. World Health Organization. World Malaria Report. https://www.wipo.int/amc/en/mediation/ (2023).

2. Suarez, C. E., Bishop, R. P., Alzan, H. F., Poole, W. A. & Cooke, B. M. Advances in the application of genetic manipulation methods to apicomplexan parasites. Int J Parasitol 47, 701–710 (2017)

3. Janse, C. J., Ramesar, J. & Waters, A. P. High-efficiency transfection and drug selection of genetically transformed blood stages of the rodent malaria parasite *Plasmodium berghei*. Nat Protoc 1, 346–356 (2006)

4. Maier, A. G. et al. Exported proteins required for virulence and rigidity of *Plasmodium falciparum*-infected human erythrocytes. Cell 134, 48–61 (2008)

5. Tewari, R. et al. The systematic functional analysis of *Plasmodium* protein kinases identifies essential regulators of mosquito transmission. Cell Host Microbe 8, 377–387 (2010)

6. Guttery, D. S. et al. Genome-wide functional analysis of *Plasmodium* protein phosphatases reveals key regulators of parasite development and differentiation. Cell Host Microbe 16, 128–140 (2014)

7. Kimmel, J. et al. Gene-by-gene screen of the unknown proteins encoded on *Plasmodium falciparum* chromosome 3. Cell Syst 14, 9–23 (2023)

8. Gomes, A. R. et al. A genome-scale vector resource enables high-throughput reverse genetic screening in a malaria parasite. Cell Host Microbe 17, 404–413 (2015)

9. Bushell, E. et al. Functional profiling of a *Plasmodium* genome reveals an abundance of essential genes. Cell 170, 260–272 (2017)

10. Russell, A. J. C. et al. Regulators of male and female sexual development are critical for the transmission of a malaria parasite. Cell Host Microbe 31, 305–319 (2023)

11. Stanway, R. R. et al. Genome-scale identification of essential metabolic processes for targeting the *Plasmodium* liver stage. Cell 179, 1112–1128 (2019)

12. Ghorbal, M. et al. Genome editing in the human malaria parasite *Plasmodium falciparum* using the CRISPR-Cas9 system. Nat Biotechnol 32, 819–821 (2014)

13. Wagner, J. C., Platt, R. J., Goldfless, S. J., Zhang, F. & Niles, J. C. Efficient CRISPR-Cas9-mediated genome editing in *Plasmodium falciparum*. Nat Methods 11, 915–918 (2014)

14. Lee, M. C. S., Lindner, S. E., Lopez-Rubio, J. J. & Llinás, M. Cutting back malaria: CRISPR/Cas9 genome editing of *Plasmodium*. Brief Funct Genomics 18, 281–289 (2019)

15. Kirti, A., Sharma, M., Rani, K. & Bansal, A. CRISPRing protozoan parasites to better understand the biology of diseases. Prog Mol Biol Transl Sci 180, 21–68 (2021)

16. Mohring, F. et al. Rapid and iterative genome editing in the malaria parasite *Plasmodium knowlesi* provides new tools for *P. vivax* research. Elife 8, e45829 (2019).

17. Shinzawa, N. et al. Improvement of CRISPR/Cas9 system by transfecting Cas9-expressing *Plasmodium berghei* with linear donor template. Commun Biol 3, 426 (2020).

18. Deligianni, E. & Kiamos, I. S. Gene editing in *Plasmodium berghei* made easy: Development of a CRISPR/Cas9 protocol using linear donor template and ribozymes for sgRNA generation. Mol Biochem Parasitol 246, 111415 (2021).

19. Nishi, T., Shinzawa, N., Yuda, M. & Iwanaga, S. Highly efficient CRISPR/Cas9 system in *Plasmodium falciparum* using Cas9-expressing parasites and a linear donor template. Sci Rep 11, 18501 (2021).

20. Sidik, S. M. et al. A Genome-wide CRISPR screen in *Toxoplasma* identifies essential apicomplexan genes. Cell 166, 1423–1435 (2016)

21. Harding, C. R. et al. Genetic screens reveal a central role for heme metabolism in artemisinin susceptibility. Nat Commun 11, 4813 (2020).

22. Wang, Y. et al. Genome-wide screens identify *Toxoplasma gondii* determinants of parasite fitness in IFNγ-activated murine macrophages. Nat Commun 11, 5258 (2020).

23. Young, J. et al. A CRISPR platform for targeted in vivo screens identifies *Toxoplasma gondii* virulence factors in mice. Nat Commun 10, 3963 (2019).

24. Ishizaki, T., Hernandez, S., Paoletta, M. S., Sanderson, T. & Bushell, E. S. C. CRISPR/Cas9 and genetic screens in malaria parasites: small genomes, big impact. Biochem Soc Trans 50, 1069–1079 (2022)

25. Smith, T. A., Lopez-Perez, G. S., Herneisen, A. L., Shortt, E. & Lourido, S. Screening the *Toxoplasma* kinome with high-throughput tagging identifies a regulator of invasion and egress. Nat Microbiol 7, 868–881 (2022)

26. Zhang, C. et al. CRISPR/Cas9 mediated sequential editing of genes critical for ookinete motility in *Plasmodium yoelii*. Mol Biochem Parasitol 212, 1–8 (2017)

27. Lin, J. W. et al. A novel ‘Gene Insertion/Marker Out’ (GIMO) method for transgene expression and gene complementation in rodent malaria parasites. PLoS One 6, e29289 (2011).

28. The Plasmodium Genome Database Collaborative. PlasmoDB: An integrative database of the Plasmodium falciparum genome. Tools for accessing and analyzing finished and unfinished sequence data. Nucleic Acids Res 29, 66–69 (2001).

29. Tufet-Bayona, M. et al. Localisation and timing of expression of putative *Plasmodium berghei* rhoptry proteins in merozoites and sporozoites. Mol Biochem Parasitol 166, 22–31 (2009)

30. Zhang, M. et al. Uncovering the essential genes of the human malaria parasite *Plasmodium falciparum* by saturation mutagenesis. Science 360, 506 (2018).

31. Qian, P. et al. A Cas9 transgenic *Plasmodium yoelii* parasite for efficient gene editing. Mol Biochem Parasitol 222, 21–28 (2018)

32. The Plasmodium genetic modification project. https://plasmogem.umu.se/pbgem/ (2024).

33. Giuliano, C. J. et al. Functional profiling of the *Toxoplasma* genome during acute mouse infection. Preprint at https://www.biorxiv.org/content/10.1101/2023.03.05.531216v1 (2023) doi:10.1101/2023.03.05.531216.

34. Kudyba, H. M. et al. Some conditions apply: systems for studying *Plasmodium falciparum* protein function. PLoS Pathog 14, e1009442 (2021).

35. Walker, M. P. & Lindner, S. E. Ribozyme-mediated, multiplex CRISPR gene editing and CRISPR interference (CRISPRi) in rodent-infectious *Plasmodium yoelii*. Journal of Biological Chemistry 294, 9555–9566 (2019)

36. Ribeiro, J. M. et al. Guide RNA selection for CRISPR-Cas9 transfections in *Plasmodium falciparum*. Int J Parasitol 48, 825–832 (2018)

37. Ramaprasad, A. & Blackman, M. J. A scaleable inducible knockout system for studying essential gene function in the malaria parasite. Preprint at https://www.biorxiv.org/content/10.1101/2024.01.14.575607v1 (2024) doi:10.1101/2024.01.14.575607.

38. Burda, P. C. et al. A *Plasmodium* phospholipase is involved in disruption of the liver stage parasitophorous vacuole membrane. PLoS Pathog 11, 1–25 (2015)

39. Peng, D. & Tarleton, R. EuPaGDT: a web tool tailored to design CRISPR guide RNAs for eukaryotic pathogens. Microb Genom 1, 1–7 (2015)

40. Amos, B. et al. VEuPathDB: The eukaryotic pathogen, vector and host bioinformatics resource center. Nucleic Acids Res 50, D898–D911 (2022)

41. Tonkin, C. J. et al. Localization of organellar proteins in *Plasmodium falciparum* using a novel set of transfection vectors and a new immunofluorescence fixation method. Mol Biochem Parasitol 137, 13–21 (2004)

42. Jonsdottir, T. K. et al. Characterisation of complexes formed by parasite proteins exported into the host cell compartment of *Plasmodium falciparum* infected red blood cells. Cell Microbiol 23, e13332 (2021).

43. Schindelin, J., et al. Fiji: An open-source platform for biological-image analysis. Nat Methods 9, 676–682 (2012).

