## Supplementary Figures for "A scalable CRISPR-Cas9 gene editing system facilitates CRISPR screens in the malaria parasite *Plasmodium berghei*"

Fig. S1

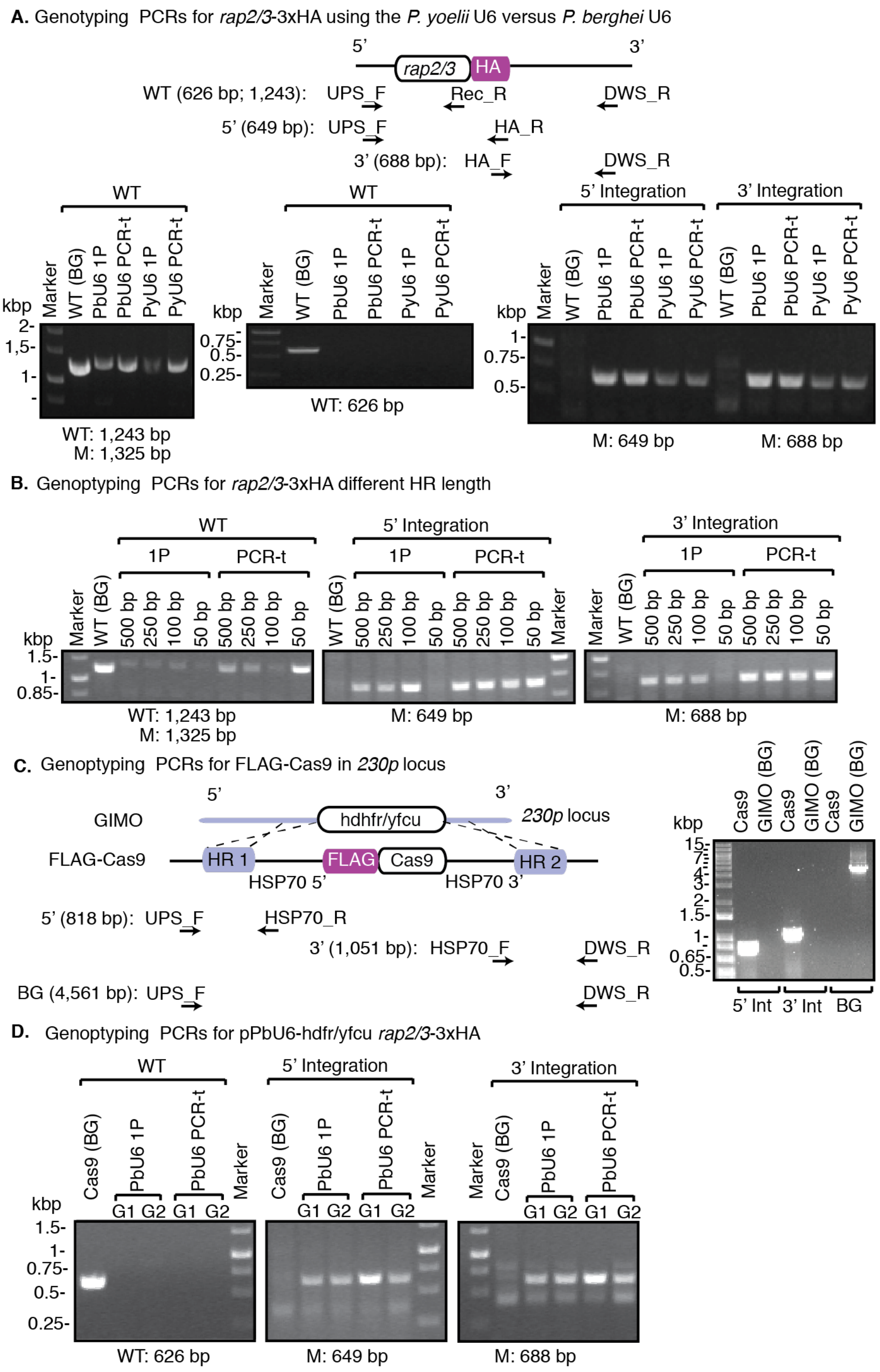


**Figure S1: Genotyping PCRs for Figure 1.** (**A**) Genotyping PCR for *rap2/3*-3xHA using either *P. yoelii* U6 or *P. berghei* U6 promoter for one-plasmid (1P) and PCR-template (PCR-t) approaches. The schematic diagram shows the location of relevant primers to confirm correct integration and presence of WT DNA. The WT serves as background (BG) line control. M = mutant (*rap2/3*-3xHA here). (**B**) Genotyping PCRs for *rap2/3*-3xHA with different homology region (HR) length. Primers are shown in panel A. (**C**) Genotyping PCRs for the integration of FLAG-Cas9 into the *230p* locus. The schematic shows where primers are located. (**D**) Genotyping PCR for *rap2/3*-3xHA using the pPbU6-hdhfr/yfcu plasmid transfected into the FLAG-Cas9 motherline. Schematic diagram in panel A displays where primers are located. G1 = gRNA 1 and G2 = gRNA 2. All primer sequences are found in **Table S1**.

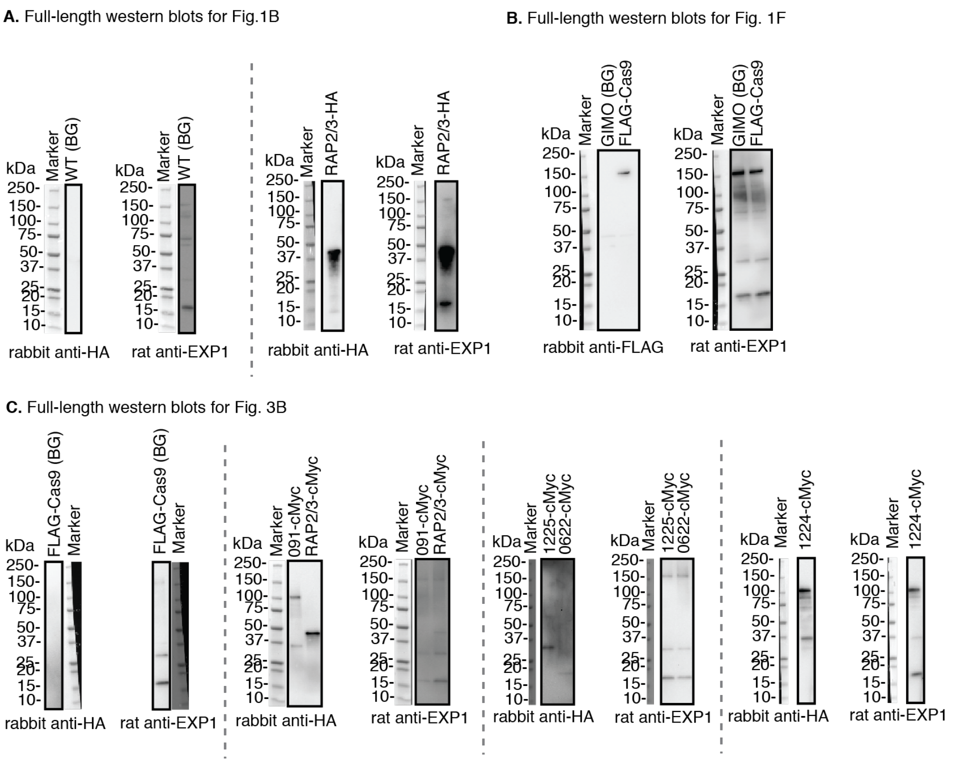


**Figure S2: Full-length western blots for Figures 1 and 3.**

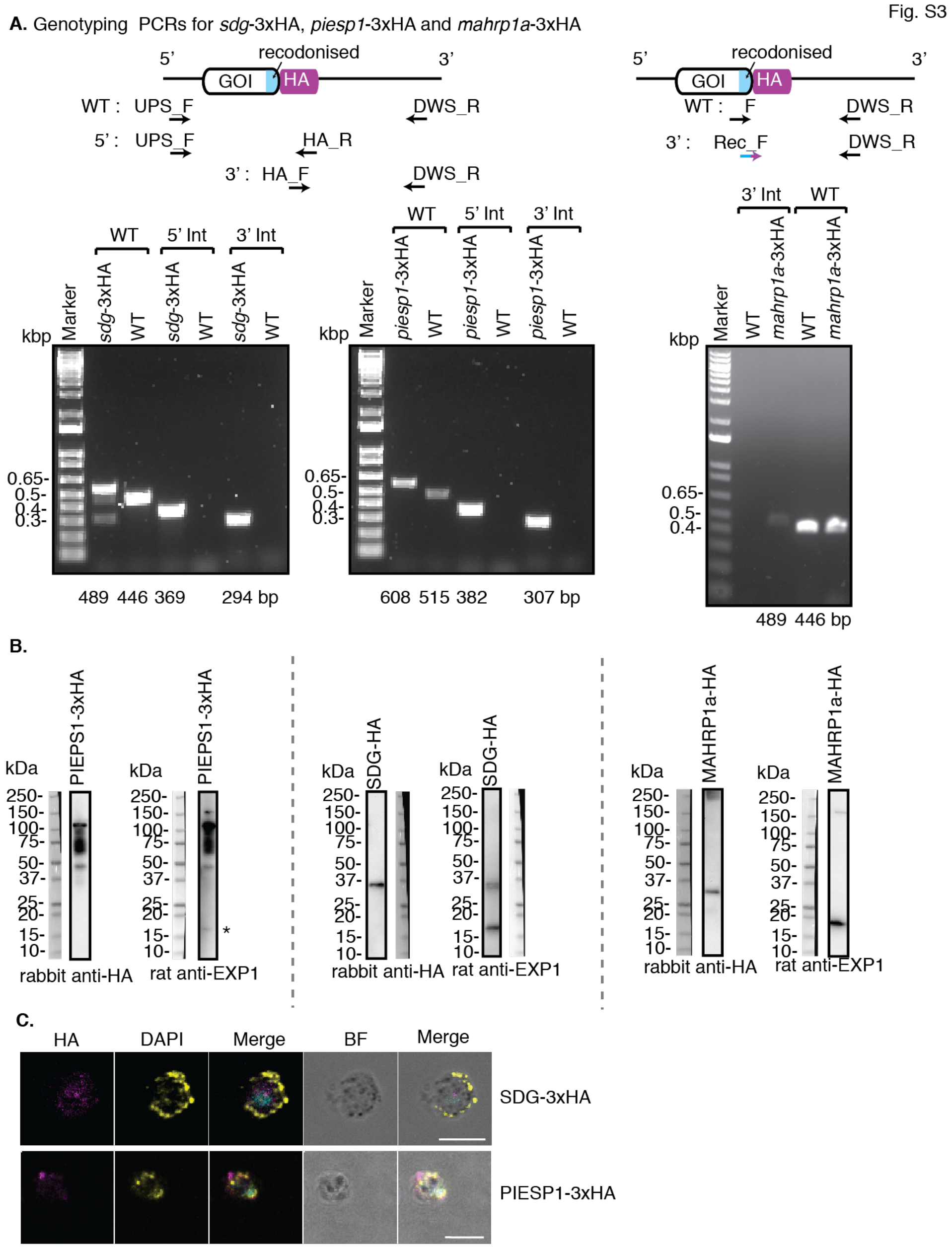


**Figure S3: Additional genes tagged using PCR-template approach.** (**A**) Genotyping PCRs for *sdg*-3xHA, *piesp1*-3xHA and *mahrp1a*-3xHA lines that were generated using PCR-template approach. Schematic diagram shows primer localisation, one for *sdg* and *piesp1* (left) and another one for *mahrp1a* (right). For *mahrp1a*, the 3' integration had a forward primer with the recodonised sequence and part of the HA tag, which is not present in WT parasites. Full-length agarose gels can be found in **Fig. S10**. All primer sequences are found in **Table S1**. (**B**) Expression of tagged proteins was observed by Western blot, where anti-HA detects protein of interest and anti-EXP1 serves as a loading control. WT parasites were used as negative control for the tag. (**C**) Immunofluorescence assay was used to localise both SDG-3xHA and PIESP1-3xHA where anti-HA detects proteins and anti-EXP1 is used as a marker for the parasitophorous vacuole membrane. Scale bars = 5 $\mu$m.

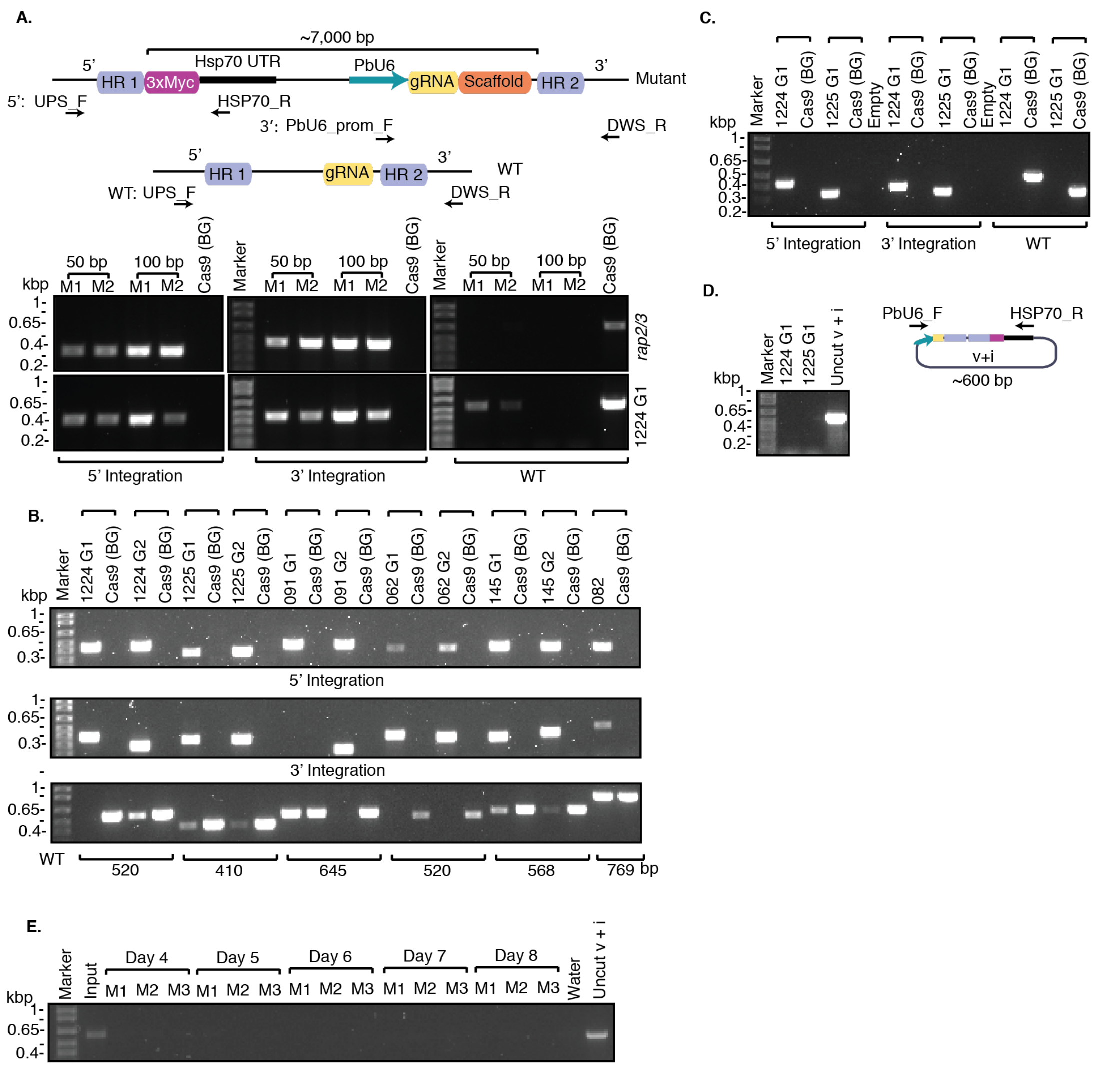


**Figure S4: PCRs for Figure 3 and 4.** (**A**) Genotyping PCRs for pPbHiT transfections for 1224200-3cMyc and rap2/3-3xcMyc comparing 50 bp and 100 bp HR length. Two independent transfections are shown, mouse 1 (M1) and mouse 2 (M2). Cas9 was used as background (BG) control for WT locus. Schematic above shows primer localisation for integration and WT PCRs. M = mutant. (**B**) Genotyping PCRs for the pHiT tagged lines using 100 bp HR length. Primers are shown in schematic in Panel A. (**C**) Genotyping PCRs for the second transfection of pPbHiT 1224200-cMyc and 1225600-cMyc where DNA was not gel extracted post *Avr*II digest. Schematic in panel **A** shows primer localisation. (**D**) No episomal plasmid was detected in genomic DNA of transfections where DNA was not gel extracted post *Avr*II digest, where pPbHiT + insert (i) serves as positive control for the presence of an episomal plasmid. (**E**) No episomal plasmid was visible in 22X knockout vector pool on collection days (day four to day eight). All primer sequences are found in **Table S1**.

Fig. S5

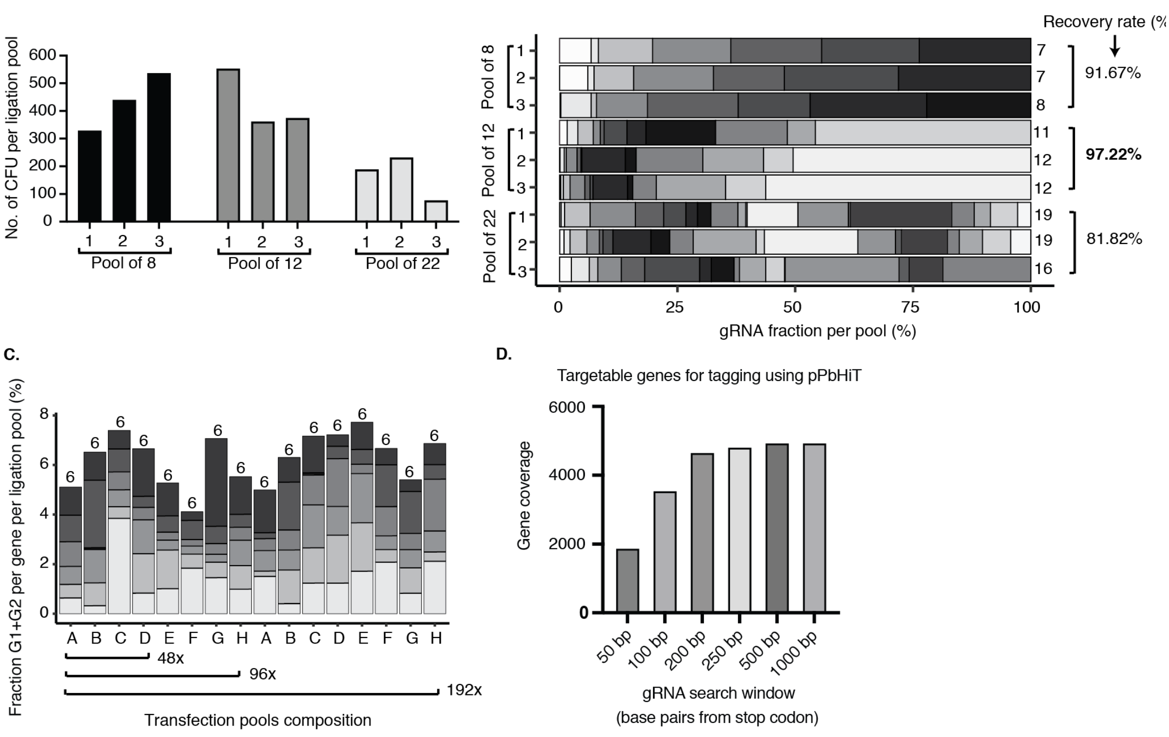


**Figure S5: Pooled pPbHit vector ligations.** (**A**) Colony numbers and (**B**) NGS results for testing ligation pools of 8, 12 or 22 inserts (synthetic fragments) cloned into pPbHiT in three biological replicates. Recovery rate is given as an average percentage (across replicates) of unique gRNA detected by NGS compared to possible maximum and the absolute number of gRNA for each ligation pool. The percentage of each gRNA within each ligation pool is plotted along the X-axis. (**C**) Vector pool composition used for larger PbHiT CRISPR screen (pools of 48X, 96X and 192X). The proportion of each gene is displayed by combining the two gRNAs targeting the same gene and is plotted along the Y-axis. This means one row (12X) on a 96-well plate should have six stacks in the histogram. (**D**) Coverage of *P. berghei* genes targetable for C-terminal tagging when allowing for different maximal distances between stop codon and gRNA.

Fig. S6

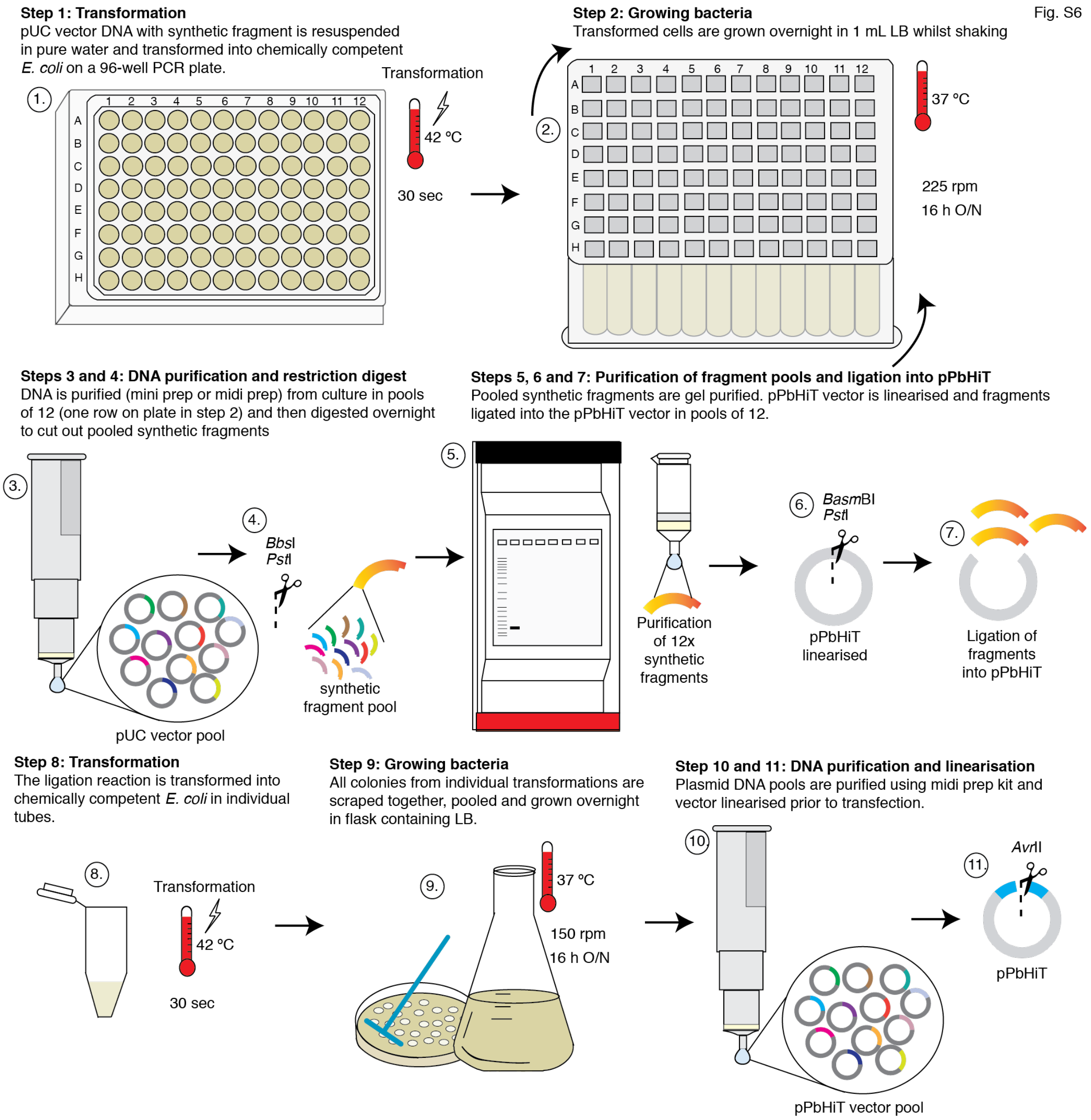


**Figure S6: Pooled pPbHiT vector ligations.** Experimental workflow for generation of pPbHiT vector pools for CRISPR screens.

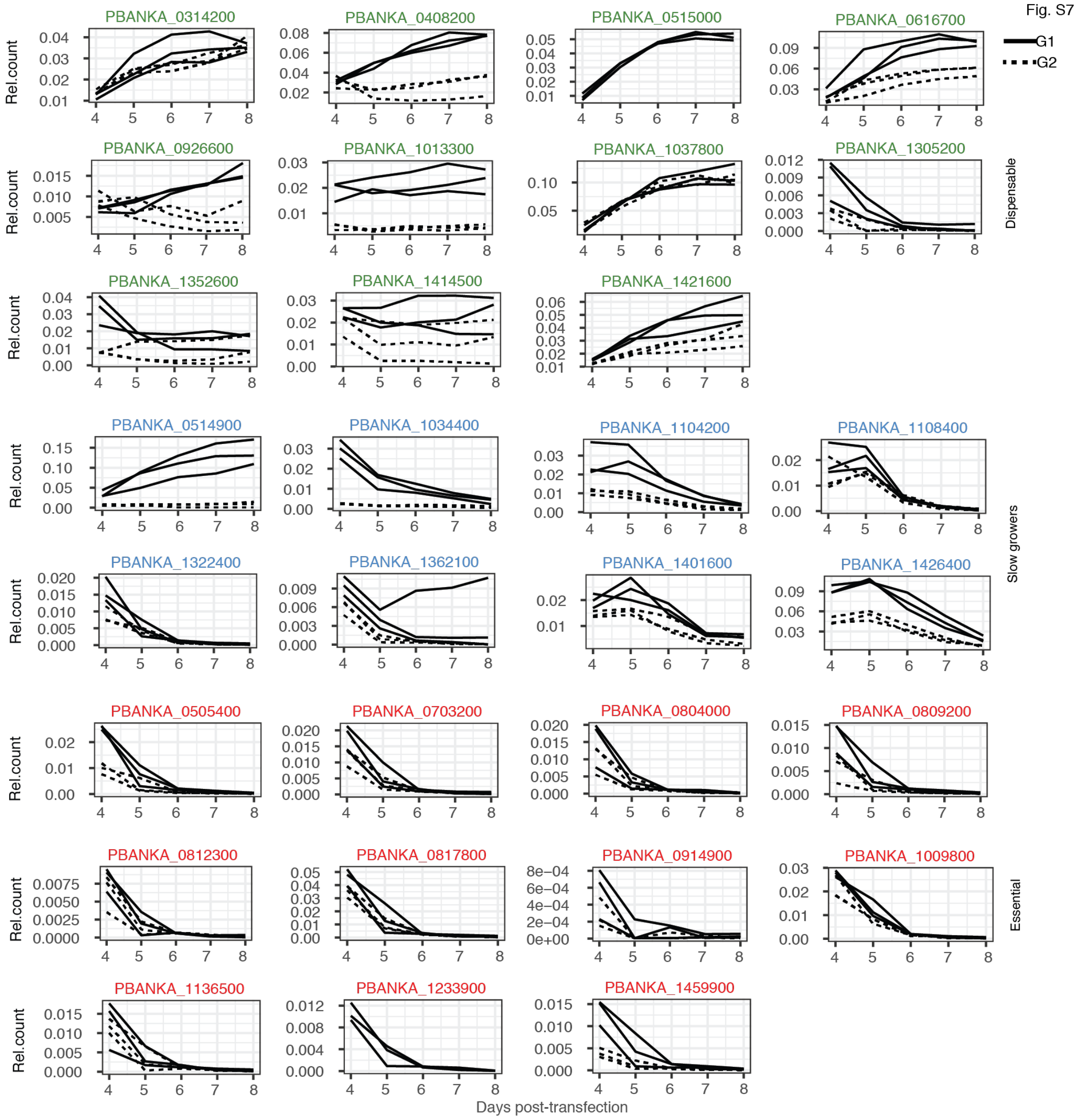


**Figure S7: Relative gRNA abundances for all genes in the 48X CRISPR pool.** The relative abundance of mutant gRNA (barcodes) was calculated for each sample as the proportion of the sum counts for all predicted dispensable genes present in the pool. Each line represents the change in relative abundance within one single mouse where the two gRNAs targeting the same gene are plotted separately. PBANKA IDs are coloured based on previously published blood stage growth phenotypes[9]: dispensable (green), slow growers (blue) or essential (red). G1 = guide one, and G2 = guide two.

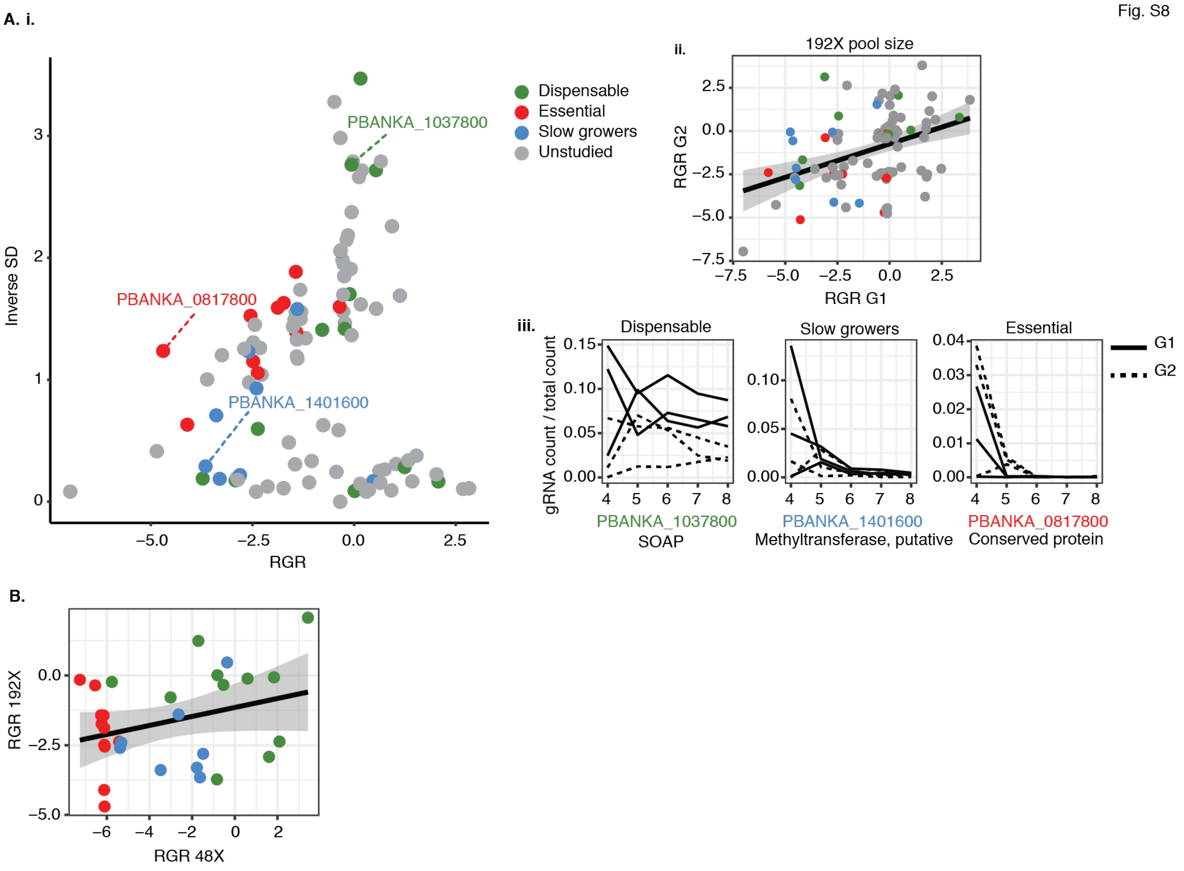


**Figure S8: PbHiT CRISPR screen using a pool of 192X vectors.** **(A)** Analysis of 192X gRNA CRISPR screen targeting 96 genes, including 24 target genes with known blood stage growth phenotypes and overlapping with the 48X gRNA pool. (**i**) Scatter plot of mutant relative growth rates (RGR) plotted against the inverse of standard deviation (1/SD). (**ii**) Correlation of RGR between two gRNA targeting the same gene. (**iii**) Selected line graphs of mutant relative abundances with each line representing counts from a single gRNA in an individual mouse. (**B**) Correlation between mutant RGR for genes overlapping between the 48X and 192X pool. Genes are coloured based on published blood stage growth phenotypes[9]: dispensable (green), slow growers (blue) or essential (red). Genes lacking *Plasmo*GEM knockout screen phenotype are classified as unstudied (grey).

Fig. S9

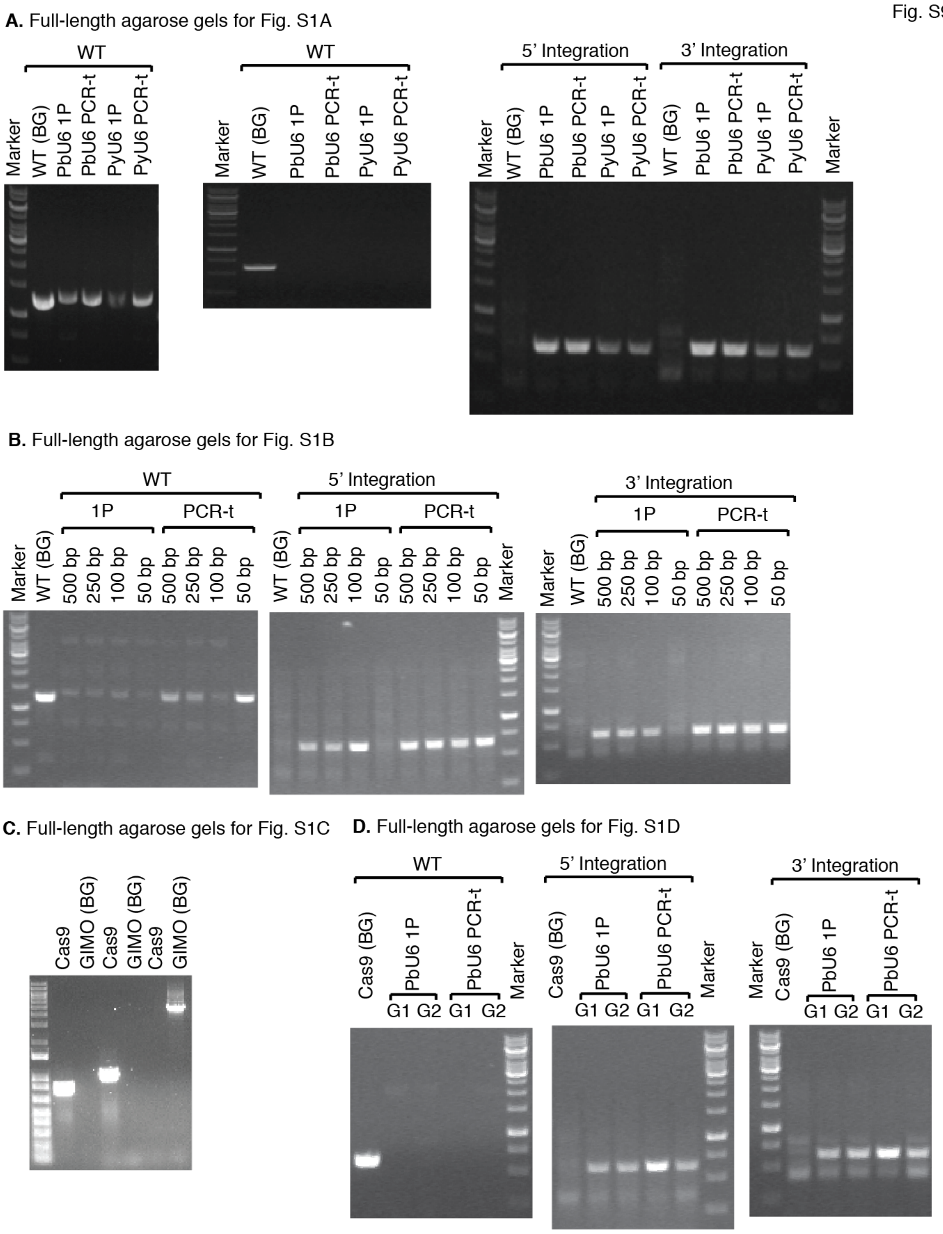


**Figure S9: Full-length agarose gels for Figure S1.**

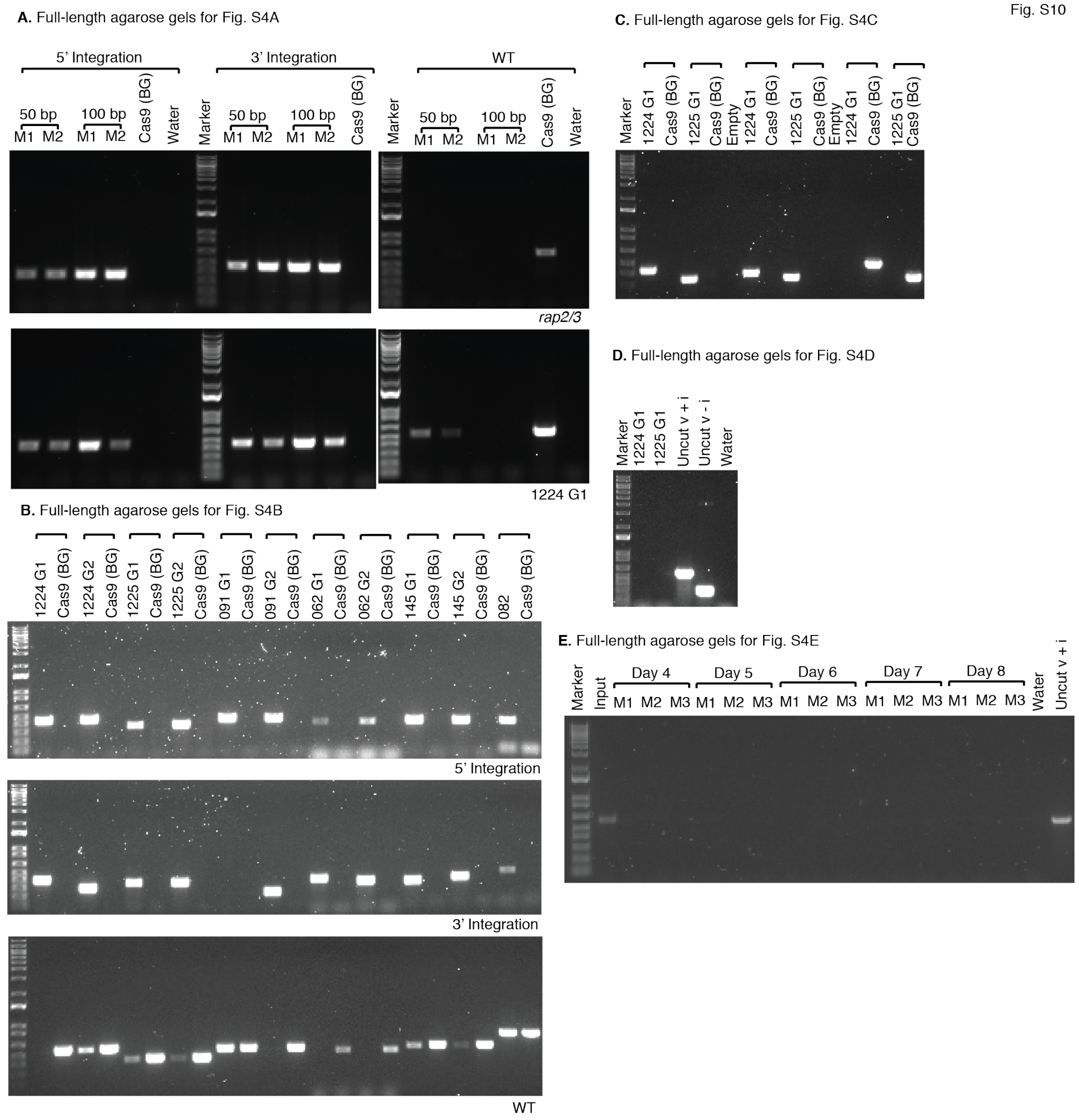


**Figure S10: Full-length agarose gels for Figure S4.**
