## Supplementary Tables for "A scalable CRISPR-Cas9 gene editing system facilitates CRISPR screens in the malaria parasite *Plasmodium berghei*"

**Table S1: Primer sequences and fragment sequences used in this study**

| Purpose | Primer/fragment name | Sequence (5'-->3') | Description |
|---|---|---|---|
| Cloning of spCas9 GIMO vector | Cas9_GIMO_F_Gib | CAATTACAATTAAAAAAAGGATGGACTATAAGGACCACGAC | Cloning of spCas9 into pL1694, primer with Gibson overhang |
| | Cas9_GIMO_R_Gib | CAGAACAATAATAGCGGCCTTACTTTTTCTTTTTTGCCTGGCC | Cloning of spCas9 into pL1694, primer with Gibson overhang |
| Cloning of new CRISPR/Cas9 vector backbones | PbU6_F_Gib | CGCAGCCTGAATGGCGAATGATCGAGAACTATTGTTTCCTTTTTTGTTTTTTTATTTTTAA<br>TTAAAATG | Cloning of PbU6 into PyCM or PyCS, primer with Gibson overhang |
| | PbU6_R_Gib | GCTCTAAAACTGAGACGAGGCCTCTACGCGTCTCCAATAATATTGTATAACTCGAAGTA<br>TGC | Cloning of PbU6 into PyCM or PyCS, primer with Gibson overhang |
| | PbU6_BbsI_R | ACTTGCTATTTCTAGCTCTAAAACAGGTCTTCTCGAAGACCCCAATAATATTGTATAACT<br>CGAAGTATG | Cloning of PbU6 into PyCM or PyCS, primer with Gibson overhang |
| Building of pPbHiT vector | pPbHiT building synthetic fragment | ATATTATTGGAGACGCTGCAGgaacaaaaacttattagcgaagaagatcttGAACAAAAATTAATAAG<br>TGAAGAAGATTTAGAGCAGAAGTTGATTTCAGAGGAGGACTTGgtcgacGCGGCCGCgTA<br>GATAAATAATTTTTTTTTTTTATATTATTGTTCTGTACTTCTTTTGTGAATAGTATTTTTACAT<br>ATGTACTGTTAATTTTAAAATGTGGAATGCCACGAACATATTTTTTCGAACAAAATATATA<br>TTTTTTGTGTCGAAATACATAAATTTGTGTACCTCCTTTTTGTTGTATATATGAAAAAGGA<br>ATAAAGTGAATGGAGTCGAATTACTCTAGACCCTACAACTTGTATATCACTACTTTATAA<br>TACTATAATATTATTATTAACATTAAACTATAGAAGAAATATCAACAATACTTATGTATTTG<br>TTCAAACTTTATCAGTATAGTTTGGTAATCGCATATCATATATTTGTTATGTGTAATATTT<br>TATGAATATGCGTCTTTTTTTTTAACTATAAATAATTCAAAATAAAATAACATAATTAACAT<br>TTGCAAGCATAATTAAGATGATTACTTTCACATTTCATCCAAAGTGGAAAATGACCACCA<br>CATTTGATCGAAGCTTTTACCACTTTTGATGCATTAGCTCTGAAATGTATACAAGCTTTC<br>ATTTGTTAATGTTACAAAGGGAAGTTTTTATTTGGGGGCACATTAAAAGTGAAAAAATTA<br>AAAGTTAAAAAAGTTAAAAAATGATCGGTGTAGTTGGTCATATAGACATGTTGTTATGCA<br>AGTCTTTCAAAACGTTCCTCATAAAATAAAAGGAGGGATCTAGAGAGCTTTCAATTCTCC<br>AAACGGTATTATTGGTATTTTCTGAAAACTGCGTTTCCTTGTTTTTATTTATATTTGACGT<br>C | Synthetic sequence used to build pPbHiT upon modification of pPbU6-hdhfr/yfcu |
| Diagnostic primers CRISPR/Cas9 vectors | gRNAseq_R | CAAATAGGGGTTCCGCGCAC | Diagnostic PCR and sequencing of gRNA insertion into pPbU6-hdhfr/yfcu and pPbU6-hdhfr/yfcu-Cas9 |

| | | | |
|---|---|---|---|
| | hsp70UTR_R | TATGTTCGTGGCATTCCACAT | Diagnostic PCR and sequencing of gRNA insertion into pPbHiT |
| | PbU6prom_F | ACATATGCGCATACTTCGAGTTATAC | Diagnostic PCR and sequencing of gRNA insertion into pPbHiT |
| PBANKA_1101400 (rap2/3) construct (not pHiT) | Recodonized rap2/3 fragment | CCAGGAGACATTTACCTTGACACCAAAATGGTGGGTGCA | |
| 500 bp construct | PB_Rap23_gRNA1_F | TATTGTATTTGGATACTAAGATGGT | Guide RNA oligo |
| | PB_Rap23_gRNA1_R | AAACACCATCTTAGTATCCAAATAC | Guide RNA oligo |
| | PB_Rap23_gRNA2_F | TATTGTATCCAAATATATATCACC | Guide RNA oligo |
| | PB_Rap23_gRNA2_R | AAACGGTGATATATATTTGGATAC | Guide RNA oligo |
| | PB_Rap23_HA_5HR_F | CTATGACCATGATTACGCCAGAGGTTAATGATAAACGCATAGATGC | Amplification primer |
| | PB_Rap23_HA_3HR_R | GGACCATGGCATGGGTACCAGAATATCCTAAAAGCATGACTATACC | Amplification primer |
| 250 bp construct | Rap23_5HR_250bp_F | CTATGACCATGATTACGCCACATACGATTTTTGGACAATTTTAGTATAGCAATAC | Amplification primer |
| | Rap23_3HR_250bp_R | GGACCATGGCATGGGTACCACTAAAAATATGGATATAAAATAAAGGGAATAAAC | Amplification primer |
| 100 bp construct | Rap23_5HR_100bp_F | CTATGACCATGATTACGCCACATATCATCCGAATTAATCTTCTTAAGAGAAAG | Amplification primer |
| | Rap23_3HR_100bp_R | GGACCATGGCATGGGTACCACACGATTTTATCTAAATATTTTTCTAATAG | Amplification primer |
| 50 bp construct | Rap23_5HR_50bp_F | CTATGACCATGATTACGCCAGTTACACCAATAAATATATGAATTTACCAGGAGAC | Amplification primer |
| | Rap23_5HR_50bp_R | GGACCATGGCATGGGTACCACTATTACATAAAAAATAAAAACTTAATAGCTC | Amplification primer |
| PBANKA_1101400 (rap2/3) genotyping | Rap2/3_UPS_FW | GCAAAGTTTATAGATGATCCTATG | Genotyping primer |
| | Rap2/3_DWS_RV | GTATATTTATATGTGTATTCCCC | Genotyping primer |
| | Rec_R | CTTAGTATCCAAATATATATCACC | Genotyping primer |
| | HA_F | CGCTTATCCCTACGATGTGCCC | Genotyping primer |
| | HA_R | CATCATATGGATATGCACCCACC | Genotyping primer |
| Note: Rap2/3 HR actual sizes | **5' HR (lenght referred in text)** | **Actual lenght** | |

| | | | |
|---|---|---|---|
| | 50 bp | 65 bp | |
| | 100 bp | 130 bp | |
| | 250 bp | 268 bp | |
| | 500 bp | 582 bp | |
| | **3' HR (lenght referred in text)** | **Actual lenght** | |
| | 50 bp | 50 bp | |
| | 100 bp | 126 bp | |
| | 250 bp | 281 bp | |
| | 500 bp | 545 bp | |
| PBANKA_0522400 (sdg) PCR-template approach | SDG_sgRNA1_F | TATTGACAATACTGAATAGCCATTT | Guide RNA oligo |
| | SDG_sgRNA1_R | AAACAAATGGCTATTCAGTATTGTC | Guide RNA oligo |
| | SDG_sgRNA2_F | TATTGACTTAAGATAGTCAGACAAA | Guide RNA oligo |
| | SDG_sgRNA2_R | AAACTTTGTCTGACTATCTTAAGTC | Guide RNA oligo |
| | SDG_HR1_F | CTGAATGTATCATCAGAAAAATGAAG | Amplification primer |
| | SDG_HR2_R | GTAAAGTCTAAAGGATAAATATGAATATCC | Amplification primer |
| PBANKA_0522400 (sdg) genotyping | SDG_UPS_FW | CAAGTTCAGTGTCATACCCCTATGT | Genotyping primer |
| | SDG_DWS_RV | CTGATTACAAAGGTGCATTTATATG | Genotyping primers |
| | HA_F | TACCCATATGACGTTCCAGACTACGCGT | Genotyping primer |
| | HA_R | AGCGTAGTCAGGTACGTCGTAAGGGTAA | Genotyping primers |
| PBANKA_0408500 (piesp1) PCR-template approach | PIESP1_sgRNA1_F | TATTGTTTCGGCTCTCTCATTATAT | Guide RNA oligo |
| | PIESP1_sgRNA1_R | AAACATATAATGAGAGAGCCGAAAC | Guide RNA oligo |
| | PIESP1_sgRNA2_F | TATTGAAATATTCATCTGGAGTGT | Guide RNA oligo |

| | | | |
|---|---|---|---|
| | PIESP1_sgRNA2_R | AAACACACTCCAGATGAATATTTC | Guide RNA oligo |
| | PIESP1_HR1_F | TCTGAAGAAGATGATGAAAATGAT | Amplification primer |
| | PIESP1_HR2_R | GTATATCAAATGAGCATGTACATGA | Amplification primer |
| PBANKA_0408500 (piesp1) genotyping | PIESP1_UPS_FW | CGTAAATAATTACGTTGTAGAAAATACTG | Genotyping primer |
| | PIESP1_DWS_RV | CGAATAGAGCTTATTTTTGTTTAC | Genotyping primers |
| | HA_F | TACCCATATGACGTTCCAGACTACGCGT | Genotyping primer |
| | HA_R | AGCGTAGTCAGGTACGTCGTAAGGGTAA | Genotyping primers |
| PBANKA_1145800 (mahrp1a) PCR-template approach | MAHRP1a_sgRNA1_FW | TATTGAGACCCCCAGGATGTACTTG | Guide RNA oligo |
| | MAHRP1a_sgRNA1_RV | AAACCAAGTACATCCTGGGGGTCT | Guide RNA oligo |
| | MAHRP1a_sgRNA2_FW | TATTGAAAGCTCATGGTACTACTAG | Guide RNA oligo |
| | MAHRP1a_sgRNA2_RV | AAACCTAGTAGTACCATGAGCTTT | Guide RNA oligo |
| | MAHRP1a_HR1_FW | GTCTGTGCATGTGTTAAGTGC | Amplification primer |
| | MAHRP1a_HR2_RV | CATGTACATCCATTAGAGCTTGAT | Amplification primer |
| PBANKA_1145800 (mahrp1a) genotyping | MAHRP1a_FW | TGCGGAGGACATAAAGCTC | Genotyping primers |
| | MAHRP1a_Rec_FW | GACAAATGAATTCTATCCATACGACGT | Genotyping primer |
| | MAHRP1a_DWS_RV | AAAGTTAAAATTCACCATTTAGCC | Genotyping primers |
| FLAG-Cas9 genotyping | HSP70_UTR_FW | GGAGGGATCTAGAGAGCTTTCAA | Genotyping primers |
| | HSP70_UTR_RV | ACAAATTCAATGAACTTCTAATATGACTC | Genotyping primers |
| | p230p_UPS_FW | GAATTGTATATGGTAAAGAACCTACTAACAC | Genotyping primers |
| | p230p_DWS_RV | GAATAGTGACTTTCAGTGAAATCGCAAA | Genotyping primers |
| Illumina libraries for gRNA sequencing | BC_pHiT_illumina_F | ACACTCTTTCCCTACACGACGCTCTTCCGATCTCGAATGCACTATTCATTTTATGGGG | Amplifies gRNA from all pPbHiT vectors and parasites, with overhang |

| | | | |
|---|---|---|---|
| | | | for priming in second PCR. Nested PCR 1 for seqeuncing libraries. |
| | BC_pHiT_illumina_R | TCGGCATTCCTGCTGAACCGCTCTTCCGATCTGACTCGGTGCCACTTTTTCA | Amplifies gRNA from all pPbHiT vectors and parasites, with overhang for priming in second PCR. Nested PCR 1 for seqeuncing libraries.. |
| | PE 1.0 | AATGATACGGCGACCACCGAGATCTACACTCTTTCCCTACACGACGCTCTTCCGATC*T | Standard Illumina library primer with Illumina adaptors to attach to flow cell. Nested PCR 2 for seqeuncing libraries. |
| | Illumina index primer 1-96 | Sequences available from Bushell et al 2017 (PMID: 28708996) | Custom indexed Illumina library primer to allow sample multiplexing, with Illumina adaptors Nested PCR 2 for seqeuncing libraries. |
| | Illumina custom index seq primer | AAGAGCGGTTCAGCAGGAATGCCGAGACCGATCTC | Custom Illumina index sequencing primer |
| pHiT 50 bp tagging constructs primers | PB_1224200_g1_50bp_FW | GAAGACggTATTTTAGATATGCATACAAA | |
| | PB_1224200_g1_50bp_RV | CTGCAGTGACTTGTTTTGATTATGC | |
| | PB_1101400_g1_50bp_FW | GAAGACggTATTAGAAAATAAGTAAATTTGATG | |
| | PB_1101400_g1_50bp_RV | CTGCAGTGCGCCGACCATCTTAGTA | |
| pHiT protection base pairs primers | All_50bp_RE_prot_bp_F | AGTCCATAGAAGACggTATT | |
| | PB_1224_50b_prot_bp_R | TAATGTTTCTGCAGTGACTTG | |
| | PB_11014_50b_prot_bp_R | TAATGTTTCTGCAGTGCGCC | |

| | | | |
|---|---|---|---|
| PbHiT 50 and 100 bp genotyping | PB_1224200_UPS_FW | GCCTTGAAACAACTAGGAGATATGAT | Genotyping primers |
| | PB_1224200_DWS_RV | GGATTGATTGCATAGCAACCATATA | Genotyping primers |
| | PB_1225600_UPS_FW | GGCTCTATATGAAAAATGCAAAATGAAGG | Genotyping primers |
| | PB_1225600_DWS_RV | GTTTGGAGTTGCTTATACTTGTAACATG | Genotyping primers |
| | PB_0914500_UPS_FW | ATGGAAATGTGAAAATTATACCTTCAAAGG | Genotyping primers |
| | PB_0914500_DWS_RV | TATAATATGCACACAAACAACGTGC | Genotyping primers |
| | PB_0622900_UPS_FW | CAGAATGGATACCCCTGATCCATT | Genotyping primers |
| | PB_0622900_DWS_RV | TTCGTAACGTATATTGCGTTTACATTG | Genotyping primers |
| | PB_1451000_UPS_FW | GACAACATTTGCTGTGCATTTAGTG | Genotyping primers |
| | PB_1451000_DWS_RV | TCCATTAAGTTGTGCTAATAAACAATTTTC | Genotyping primers |
| | PB_0829400_UPS_FW | CTCCTCTGAAAATGTAAAAAATGCCA | Genotyping primers |
| | PB_0829400_DWS_RV | GTTTCTCCTTGTTTTTTTGTTGATATATGC | Genotyping primers |
| | PB_1101400_UPS_Fw | GAATTCTGATTACATATCATCCG | Genotyping primers |
| | PB_1101400_DWS_Rv | GGTATAGTCATGCTTTTAGGATATTC | Genotyping primers |

**Table S2: Parasitemia counts for Figures 1 and 3**

| Figure 1Ai | | | | | | | | |
|---|---|---|---|---|---|---|---|---|
| | P. yoelii One plasmid | | P. yoelii PCR-fragment | | P. yoelii no donor | | | |
| | Mouse 1 | Mouse 2 | Mouse 1 | Mouse 2 | Mouse 1 | Mouse 2 | | |
| Day 4 | 0 | 0.08417508418 | 0 | 0 | 0 | 0 | | |
| Day 5 | 0.09505703422 | 0.1895734597 | 0.09216589862 | 0 | 0 | 0 | | |
| Day 6 | 0.4562043796 | 0.2949852507 | 0 | 0.09057971014 | 0 | 0 | | |
| Day 7 | 2.97219559 | 2.364864865 | 0.1748251748 | 0.09033423668 | 0 | 0 | | |
| Day 8 | 3.646833013 | 5.975103734 | 0.4389815628 | 0.2915451895 | 0 | 0 | | |
| Day 9 | N/A | N/A | 3.093721565 | 2.844036697 | 0 | 0 | | |
| Day 10 | N/A | N/A | 7.527881041 | 5.838041431 | 0 | 0 | | |
| Day 11 | N/A | N/A | 6.107566089 | 5.450941526 | 0 | 0 | | |
| | P. berghei One plasmid | | P. berghei PCR-fragment | | P. berghei no donor | | | |
| | Mouse 1 | Mouse 2 | Mouse 1 | Mouse 2 | Mouse 1 | Mouse 2 | | |
| Day 4 | 0.09615384615 | 0.1953125 | 0.0946969697 | 0 | 0 | 0 | | |
| Day 5 | 0.4901960784 | 0.198019802 | 0.1968503937 | 0.09514747859 | 0 | 0 | | |
| Day 6 | 1.532912534 | 1.168014376 | 0.09250693802 | 0.09950248756 | 0 | 0 | | |
| Day 7 | 3.818827709 | 3.062200957 | 0.09174311927 | 0.1782531194 | 0 | 0 | | |
| Day 8 | 7.446808511 | 8.252427184 | 0.4 | 0.2906976744 | 0 | 0 | | |
| Day 9 | 7.632093933 | 4.812319538 | 1.263362488 | 0.8204193254 | 0 | 0 | | |
| Day 10 | N/A | N/A | 4.929577465 | 2.235179786 | 0 | 0 | | |
| Day 11 | N/A | N/A | 11.34020619 | 6.150793651 | 0 | 0 | | |
| **Figure 1Aii** | | | | | | | | |
| | One plasmid, 500 bp | | One plasmid, 250 bp | | One plasmid, 100 bp | | One plasmid, 50 bp | |
| | Mouse 1 | Mouse 2 | Mouse 1 | Mouse 2 | Mouse 1 | Mouse 2 | Mouse 1 | Mouse 2 |
| Day 4 | 0 | 0 | 0 | 0 | 0 | 0 | 0 | 0 |

| | | | | | | | | |
|---|---|---|---|---|---|---|---|---|
| Day 5 | 0.1742160279 | 0.2617801047 | 0.1813236627 | 0 | 0 | 0.0998003992 | 0 | 0 |
| Day 6 | 0.6711409396 | 1.391650099 | 0.1943634597 | 0.1872659176 | 0.3518029903 | 0.2961500494 | 0 | 0 |
| Day 7 | 4.12979351 | 4.8 | 0.826446281 | 0.8952551477 | 0.5703422053 | 1.157184185 | 0 | 0 |
| Day 8 | 6.343283582 | 6.048780488 | 5.135658915 | 6.978879706 | 3.796203796 | 3.237410072 | 0.580270793 | 0.3549245785 |
| Day 9 | N/A | N/A | N/A | N/A | 5.221674877 | 4.835589942 | 2.263779528 | 1.833976834 |
| Day 10 | N/A | N/A | N/A | N/A | N/A | N/A | 5.722326454 | 5.801376598 |
| | PCR-fragment, 500 bp | | PCR-fragment, 250 bp | | PCR-fragment, 100 bp | | PCR-fragment, 50 bp | |
| | Mouse 1 | Mouse 2 | Mouse 1 | Mouse 2 | Mouse 1 | Mouse 2 | Mouse 1 | Mouse 2 |
| Day 4 | 0 | 0 | 0 | 0 | 0 | 0 | 0 | 0 |
| Day 5 | 0 | 0 | 0 | 0 | 0 | 0 | 0 | 0 |
| Day 6 | 0.09910802775 | 0 | 0 | 0 | 0 | 0 | 0 | 0 |
| Day 7 | 0.1843317972 | 0.09009009009 | 0.1945525292 | 0.09115770283 | 0 | 0 | 0 | 0 |
| Day 8 | 0.9460737938 | 1.764705882 | 2.418604651 | 0.5623242737 | 0.6641366224 | 0.1976284585 | 0.7627118644 | 1.479289941 |
| Day 9 | 2.389705882 | 3.182256509 | 2.93040293 | 1.868327402 | 1.320754717 | 1.98019802 | 1.744186047 | 2.189781022 |
| Day 10 | 4.658077304 | 5.454545455 | 3.734827264 | 4.326923077 | 3.61328125 | 3.746397695 | 2.756653992 | 3.424657534 |
| **Figure 1C** | | | | | | | | |
| | PbGIMO | | | FLAG-Cas9 | | | | |
| | Mouse 1 | Mouse 2 | Mouse 3 | Mouse 1 | Mouse 2 | Mouse 3 | | |
| Day 1 | 0.08503401361 | 0.09310986965 | 0.2985074627 | 0.1683501684 | 0.08710801394 | 0.1718213058 | | |
| Day 2 | 0.3894839338 | 0.285986654 | 0.4212299916 | 0.5576208178 | 0.2879078695 | 0.1774622893 | | |
| Day 3 | 1.56114484 | 1.617507136 | 2.00729927 | 2.140945584 | 2.042801556 | 1.561065197 | | |
| Day 4 | 5.284147557 | 4.545454545 | 6.338028169 | 5.678537055 | 6.932573599 | 5.544747082 | | |

| | | | | | | | | | | | |
|---|---|---|---|---|---|---|---|---|---|---|---|
| Day 5 | 7.605633803 | 6.951871658 | 9.067357513 | 7.222222222 | 7.047619048 | 8.600182983 | | | | | |
| **Figure 1F** | | | | | | | | | | | |
| | pPbU6-hdfr/yfcu Rap2/3-3xHA one plasmid | | pPbU6-hdfr/yfcu Rap2/3-3xHA PCR-template | | No donor | | | | | | |
| Day 4 | 0 | 0 | 0 | 0 | 0 | 0 | | | | | |
| Day 5 | 0.5115089514 | 0.2830188679 | 0.09803921569 | 0.1943634597 | 0 | 0 | | | | | |
| Day 6 | 1.967799642 | 3.489531406 | 0.3787878788 | 0.5988023952 | 0 | 0 | | | | | |
| Day 7 | 4.207699194 | 5.927342256 | 0.5870841487 | 0.6876227898 | 0 | 0 | | | | | |
| Day 8 | N/A | N/A | 2.53411306 | 1.360544218 | 0 | 0 | | | | | |
| Day 9 | N/A | N/A | 1.996007984 | 3.411306043 | 0 | 0 | | | | | |
| Day 10 | N/A | N/A | 7.848837209 | 6.413994169 | 0 | 0 | | | | | |
| **Figure 3Ai** | | | | | | | | | | | |
| | rap2/3 50 bp | | rap2/3 100 bp | | PBANKA_1224200 G1 50 bp | | PBANKA_1224200 G1 100 bp | | | | |
| | M1 | M2 | M1 | M2 | M1 | M2 | M1 | M2 | | | |
| Day 4 | 0 | 0 | 0,05 | 0 | 0 | 0 | 0 | 0 | | | |
| Day 5 | 0,05 | 0 | 0,8 | 0,2 | 0 | 0 | 0,1 | 0 | | | |
| Day 6 | 0,1 | 0 | 2,4 | 2,3 | 0 | 0,05 | 1,6 | 0,3 | | | |
| Day 7 | 0,3 | 0,08 | 4,3 | 3,4 | 0 | 0,08 | 3,5 | 1,2 | | | |
| Day 8 | 1,05 | 1,08 | 2,25 | 5,42 | 0,4 | 0,2 | 3,95 | 2,63 | | | |
| Day 9 | 2,19 | 3,34 | 1,78 | 4,81 | 2 | 1,19 | 2,7 | 2,38 | | | |
| Day 10 | 2,72 | 4,31 | 2,22 | 3,25 | 1,62 | 1,86 | 1,71 | 1,52 | | | |
| **Figure 3Aii** | | | | | | | | | | | |
| | PBANKA_1224200 | | PBANKA_1225600 | | PBANKA_0914500 | | PBANKA_0622900 | | PBANKA_1451000 | | PBANKA_0829400 |
| | G1 | G2 | G1 | G2 | G1 | G2 | G1 | G2 | G1 | G2 | G1 |
| Day 4 | G1 | G2 | G1 | G2 | G1 | G2 | G1 | G2 | G1 | G2 | G1 |
| Day 5 | 0,4 | 0,3 | 0,3 | 0,3 | 0,2 | 0,1 | 0,2 | 0 | 0,3 | 0,1 | 0 |
| Day 6 | 1,5 | 0,2 | 0,1 | 0,3 | 0,1 | 0,5 | 0,4 | 0,3 | 0,4 | 0 | 0 |
| Day 7 | 5 | 1,3 | 0,4 | 2 | 0 | 2,5 | 2,5 | 2,5 | 4,4 | 0,3 | 0,2 |
| Day 8 | | 3,2 | 0,4 | | 0,4 | | | 4,4 | | 2 | 0,6 |
| Day 9 | | | 2,3 | | 0,8 | | | | | 4 | 1,3 |
| Day 10 | | | | | 1,6 | | | | | | 2,3 |

**Table S3: Genes tagged using PbHiT and relevant sequences**

| Target gene ID | Gene name | Guide RNA | Guide sequence (5’-->3’) | HR length (bp) | HR1 | HR2 |
|---|---|---|---|---|---|---|
| PBANKA_1224200 | DnaJ protein, putative | G1 | AGATATGCATACAAAACA | 100 | AGAAGAAGAAGCAGCAGCAGAAGCAGCAGCAGAAGCTGAATCAACCAAAATTGAGAGGGATGAAAAATACAGTGGTCTAAGGCATAATCAAAACAAGTCA | TGTACTTTCGCTTCGAATTAAATTCTATGACTTTTTTCTCTTTTAATTAAAAAGTGCAGCATCTGTAATATATATAACTCCCTTTGACATTCAAATTCCA |
| | | | | 50 | TTGAGAGGGATGAAAAATACAGTGGTCTAAGGCATAATCAAAACAAGTCA | TGTACTTTCGCTTCGAATTAAATTCTATGACTTTTTTCTCTTTTAATTAA |
| | | G2 | TCTTGGAATTTGAATGTCAA | 100 | AGAAGAAGAAGCAGCAGCAGAAGCAGCAGCAGAAGCTGAATCAACCAAAATTGAGAGGGATGAAAAATACAGTGGTCTAAGGCATAATCAAAACAAGTCA | CAAATCAATTTAATTCAAATGCTTGATGATTTATAAAAATTTTATTTAAACATAAAATAAAATAGTAACTTCCTATAATTTTAAGTGCCATACACTGTGT |
| PBANKA_1101400 | rhoptry-associated protein 2/3 | G1 | AGAAAATAAGTAAATTTGAT | 100 | AAGTTTTTTTGATGCACTTGATAGTACTTTAAATTGTTACACCAATAAATATATGAATTTACCTGGTGATATATATTTGGATACTAAGATGGTCGGCGCA | TGTTTATTCCCTTTATTTTATATCCATATTTTTAGTTGAAAACTTTTTTATATATTTAAAGGTTATACTGACAAATGGTAATATATTTTCCCCTTGCATC |
| | | | | 50 | ATATGAATTTACCTGGTGATATATATTTGGATACTAAGATGGTCGGCGCA | TGTTTATTCCCTTTATTTTATATCCATATTTTTAGTTGAAAACTTTTTTA |
| PBANKA_1225600 | alpha/beta hydrolase, putative | G1 | CAAATAAAACCACAAAAAAG | 100 | CTATTGGGTTGCCAATGGAAAACATAACGATGTTGAATTAATTGACAATAAAAAATTCAACGAAAACATCAAGTTTTTCCTAAATTTTTTAAATAATTCA | AACTTTTTTAAATGTGAAAGTTATCAAAAAAATATATATAATTTGCATGTTATTTTATTATTTTTTGTATTCAAAATATATGTTTATGTAATACCTTATT |
| | | G2 | AATGAATTTCCCCTTTTTTG | 100 | CTATTGGGTTGCCAATGGAAAACATAACGATGTTGAATTAATTGACAATAAAAAATTCAACGAAAACATCAAGTTTTTCCTAAATTTTTTAAATAATTCA | TTTTATTTGAACTTTTTTAAATGTGAAAGTTATCAAAAAAATATATATAATTTGCATGTTATTTTATTATTTTTTGTATTCAAAATATATGTTTATGTAA |
| PBANKA_0914500 | conserved Plasmodium protein, unknown function | G1 | ATAACAATAATAATAATACA | 100 | TTATATTGACATATTTTACGATAAAATGAGCGAAATTATGAAAAACAACATGAATTTGGAATTATTTGGAAAAACATTCTTTTTACATTTGGAAAAAAAA | TATTTTTTTAGTTGCAGTATCTATATGATTCTCTATATATTATATCAAATAATTCCTGTGAAGAAATTTATTTCGTAAGTTTCACAAAATTTTCGAGATT |
| | | G2 | TACGAAATAAATTTCTTCAC | 100 | TTATATTGACATATTTTACGATAAAATGAGCGAAATTATGAAAAACAACATGAATTTGGAATTATTTGGAAAAACATTCTTTTTACATTTGGAAAAAAAA | AGTTTCACAAAATTTTCGAGATTTTTAAGTACCCAGACGCCAGGCTGCTTTAATTTGCATTTAATTTATTATTATTATTTTTTTTAATAATTAACTTTAG |
| PBANKA_0622900 | conserved Plasmodium protein, unknown function | G1 | TCCATAATACATATAGCAAA | 100 | AGTAACTGTTTATAATAAACATGGAGAACCATTATATTTTTATATAAATAAAAAGAAAACAAAAAAAGACTCCAAACTAAGAAAGAAAAAAAAATAAATCT | ATAAAGTTGTTAGTTTATAATACCTGTCAAAAACATTATTTCGTTGATTTTTATATATGCGTGCATTAAGATAGAATTGTTGTAAATGCTACGCTCATGT |

| | | | | | | |
|---|---|---|---|---|---|---|
| | | G2 | AACGAAATAATGTTTTTGAC | 100 | AGTAACTGTTTATAATAAACATGGAGAACCATTATATTTTTATATAAATAAAAAGAAAACAAAAAAAGACTCCAAACTAAGAAAGAAAAAAAAATAAATCT | GATTTTTATATATGCGTGCATTAAGATAGAATTGTTGTAAATGCTACGCTCATGTTTAATAATTTTGTTCCAAGAGATTATATATCATATACATTTGATG |
| PBANKA_1451000 | conserved Plasmodium protein, unknown function | G1 | AAAAAGTATGACCCTATAAG | 100 | TTATACAAGCGATGACAAATTAACATTACTTGTTAATAATCAAACGAAAAATTATTCCATAAATCAGAAAAAAAAAAAAAAAAAAAAAAAAAATCTGGAAAA | AACCGTATTTAGTCATTAATGTATTACTATATATTATTAAATACCGCTAATTTTTTTTTTTTTATGTACATACATGTATAATTGCATTCCGAGCTATTTG |
| | | G2 | AAAAAATAAGGTATTTCAAT | 100 | TTATACAAGCGATGACAAATTAACATTACTTGTTAATAATCAAACGAAAAATTATTCCATAAATCAGAAAAAAAAAAAAAAAAAAAAAAAAAATCTGGAAAA | ATTGTCAATATATATATTCCCCTTATAGGGTCATACTTTTTTAACCGTATTTAGTCATTAATGTATTACTATATATATTATTAAATACCGCTAATTTTTTTTTTTT |
| PBANKA_0829400 | conserved Plasmodium protein, unknown function | G1 | ACAAAAAAAAACAAATTATA | 100 | AGTATCGATTTTATTAGTATGTTCAGCTGTTTATTTTACTTTCAAAACATTAGCAAATAAAATAATAAAATTCTCACATATAAGTGCTATAAAATTCATT | ACATGCGATTTTGATATAATATTTTAAAATTTGTGTGTACTATTATTAAAGCTATCTTATATTATTTTTTTTCATTTTTTTTATATTAGTATTTAATTTTT |

**Table S4: Knockout pools of 22X pPbHiT vectors**

| Target gene ID | Gene name | PlasmoGEM phenotype | Spike-in control in larger CRISPR pools | G1 sequence (5’-->3’) | G2 sequence (5’-->3’) | HR1 | HR2 |
|---|---|---|---|---|---|---|---|
| PBANKA_0515000 | ookinete surface protein P25 | Dispensable | Yes | ATATTACAAGAGCATATTGG | ACCGCAAACTAATGAACATG | ATATTTCCATTTTATACAATACATAAAAGCCCATAAAAAAAATATATACACTTTATTAATAAAAATTTTATTTTTGTTATTCGTTTAAATTTATTTAAAA | ATAAACAAATATACCTGGATAATTTTCACTAATTCAACCTTAACTTTTAAGTTTAAACGCTTTATAGTTATATTTTTGTGGGCAATAAAAAAATATATATA |
| PBANKA_0933700 | mitogen-activated protein kinase 2, MAPK2 | Dispensable | No | ATTGATTCAAAAAAAAAACG | CAGATTATGTAGCAACACGT | TTATCATAAAATTGTGCATTAACAGTTAGAAGAGGATTGCCATTTTTTGTTTTTAATTTTTATGCTATTATTTTTTTCTTAATTTTTTGGACAAAAAAAA | TTTCAAATTATAATTACTCGAAAATAAATATGTATGTATCATAAACTGAATTTTGGGGTAACATATAAAAATAAATAAATGTAAAAAATAAAACCATATT |
| PBANKA_1013300 | mitogen-activated protein kinase 1, MAPK1 | Dispensable | Yes | ATTGATTCAAAAAAAAAACG | CAGATTATGTAGCAACACGT | TTATCATAAAATTGTGCATTAACAGTTAGAAGAGGATTGCCATTTTTTGTTTTTAATTTTTATGCTATTATTTTTTTCTTAATTTTTTGGACAAAAAAAA | TTTCAAATTATAATTACTCGAAAATAAATATGTATGTATCATAAACTGAATTTTGGGGTAACATATAAAAATAAATAAATGTAAAAAATAAAACCATATT |
| PBANKA_1037800 | secreted ookinete adhesive protein, SOAP | Dispensable | Yes | TCGTGAAGATGCCTTAACAT | TAGTCCCTTTGCATGTGCGG | TAAATATTGCTTACATATTCCTCTCTATAAATATATATATTCTTTTTAGCAATATATTTCTCTTATATATATACCCTTTTTTGTATTTATATTAATCAAA | TTATGTGTAAATACTGTATTATTATAAGGAAATATTTATTTATCAAAAAATATCAAAAGAAAAAATTATTTTTTTGTGTGACTTAGTCCGCATTATATAT |
| PBANKA_1034400 | plasmepsin IV, PM IV | Slow | Yes | TGACATAAAACTATTCTGAG | AAAGAGTCAAATTACTCCAA | AATTTATTTTTTTTATATAATCCGTCATTATTTTTACATTATTATGCTTTACAACATATATATATATCCAAATTTTTTTTTCTCCTTTAATTAGTTCAAA | ATAAAAATAAAAAAAATTATATATGATATATTACACGTACCATAACATGCCTGCTTTTATATATTTTATGTAAGTATCATCATATATATTTTAACAAAAG |
| PBANKA_1101400 | rhoptry-associated protein 2/3, RAP2/3 | Slow | No | TGCTTGAACCAAATCGTCGG | ATCATCTATAAACTTTGCAG | AGTTATTATTACTTTATTATTTTATATATATATTTTTATTTACAACGCATATATTAATTGCGCCAGTTATATTTTTATATAAAACGAAAGTGAAGGCAAA | GTTTTTTTATCAATAAATGAGCTATTAAGTTTTTATTTTTTATGTAATAGTTATATATATATATTATGTTTTTTTATTGATGATATGTTATTTAATCTAT |
| PBANKA_1104200 | 2-oxoisovalerate dehydrogenase subunit beta, | Slow | Yes | TGGGGTGCAGTAGGACATGG | AAAAATAAAATAATTCAAGG | AATATAATGTATCCATATCAATCATAGTGCCGCAGTTTCATTTAATTAGTGTTACTATTATTACAATTTTTATTTTTTATAAATCCGATTAACATATAAA | AATTACAAAACTAATTTTTTATTATCTCGTTTTTAATCATTATACATAACAAGGAAATATAATTTTATCTACCCTTATTTGTGTAAATCACAGCTTTTTT |

| | | | | | | | |
|---|---|---|---|---|---|---|---|
| | mitochondrial, putative, BCKDHB | | | | | | |
| PBANKA_1401600 | methyltransferase, putative | Slow | Yes | ATTGTCGCTTCATATGCTGG | GTACTAGTCATAATTTCATG | CACTTCTTCATGATCACATTGCTTTCATAAGAAGCTCATACTTAATCAGAACCATATCAGGTGTAGATAATTATTTTATTTATTTAAATACGTGTTAACA | TAAATCTTCAAATCAAACACATGTTTACCTATAACATATGCACACATATACTTAAAACATGAAATATTATTAATGTATAACATAGTTTTTATTACATTTC |
| PBANKA_0211000 | mitochondrial import inner membrane translocase subunit TIM50, putative, | Essential | No | CAAGAAGTAATATCAAAGTG | ACAACGCCAAAAAACAACAA | AGTATTTAAAAATACTACTAAATTTGATTTTTTTGCTATTTTTCTTAATGAAATTATAAAAACAACATATTGAAAAAGTGGTGAATGAAATAAAGATAGA | AGGGAAAAAATTTGGGAAAAAATTTGGGAAAAAATGAGAAAAAATTTGGGAAAAAAATATGAAAAAATGAAAATATTTTAAGCAAAATGCATATATGTATA |
| PBANKA_0706400 | conserved Plasmodium protein, unknown function | Essential | No | TTGGCGAAAGAAGAGGAAAG | AAAAAGAGAAGATTTCAATG | CATATGAAATAATTTTTTAATTATGATTATAGTCAAATTATAGTTAATTAGAAGTAATAATTATATATAAGGAACTTTGAGAATACGAAGAAAAAAAAAC | TTTAATGATATTATTTTATTTCATTTCATTTTTTCCGTTGTCTTTTCTTGTTTATGGCATCTCATGGTGTACTACATACCGTTTGGTTTGTCCACGTTGA |
| PBANKA_1039700 | cytochrome c oxidase subunit ApiCOX19, putative | Essential | No | ATAGATGATTTAAATAGTGA | TGACCATAAGAACCAAACAA | CTAATATTGAAGAGTAGAAAGGGGAGAAGATTACAAAGCCAATAATTTTCAAAAATAATCACAAGTTCAATTATATTACCATATATGTGCGAACACAAGA | ACAAAAAAATTGTAGTAGCATATGCCTACAAATTCCTTTTAAAGCAAATGCCCTTATATGCGCATTCCATTCAATTGTTTATGTTTATACTCCACTTTGTA |
| PBANKA_1214100 | tubulin binding cofactor c, putative | Essential | No | AAACAGAACTTAATATGTCA | ATATCATATTATATTGCATG | ATCATTTTTTTTATCAATATTAAAATTGGTGTATCCTTTTACAAATATACAATTATTAATATACTTATTATTTATTTGTTTTCAAAAAAAAAACAAAACA | GAAAAATAAACATATATTTTAAATCACCCTGGTGTAATATGCATATGTGTATTATATAATCTTGTAAATATCAATTTTTTTTTTTTTAATTTTTCCGCAA |

**Table S5: Knockout pools of 48X, 96X and 192X pPbHiT vectors and spiking controls**

| | | | Target gene ID | Gene name | PlasmoGEM phenotype | Guide 1 sequence (5'-->3') | Guide 2 sequence (5'-->3') | | |
|---|---|---|---|---|---|---|---|---|---|
| From pool of 22X knockout vectors | | | PBANKA_1013300 | mitogen-activated protein kinase 1, MAPK1 | Dispensable | ATTGATTCAAAAAAAAAACG | CAGATTATGTAGCAACACGT | | |
| | | | PBANKA_1037800 | secreted ookinete adhesive protein, SOAP | Dispensable | TCGTGAAGATGCCTTAACAT | TAGTCCCTTTGCATGTGCGG | | |
| | | | PBANKA_0515000 | ookinete surface protein P25 | Dispensable | ATATTACAAGAGCATATTGG | | | |
| | | | PBANKA_1401600 | methyltransferase, putative | Slow | ATTGTCGCTTCATATGCTGG | GTACTAGTCATAATTTCATG | | |
| | | | PBANKA_1034400 | plasmepsin IV, PM IV | Slow | TGACATAAAACTATTCTGAG | AAAGAGTCAAATTACTCCAA | | |
| | | | PBANKA_1104200 | 2-oxoisovalerate dehydrogenase subunit beta, mitochondrial, putative, BCKDHB | Slow | TGGGGTGCAGTAGGACATGG | AAAAATAAAATAATTCAAGG | | |
| **Larger CRISPR pools** | | | | | | | | | |
| **Pool sizes** | | | **Target gene ID** | **Gene name** | **PlasmoGEM phenotype** | **G1 sequence (5'-->3')** | **G2 sequence (5'-->3')** | **HR1** | **HR2** |
| 192X | 96X | 48X | PBANKA_1459900 | signal recognition particle receptor subunit beta, putative (SRPRB) | Essential | TTTCGTCTTACCACGGGTAT | TGAAAGGGCTCCAATACCCG | TGTTGGTGAAAAAAAATATAAACATAGAATAAATTAATAAATATATTAATAAAAAACGAGTATATGAATGAATAAATATAATAATTTAATGAAATAAAAA | TTTATGTCTATTAAAAAAAATCTATATGTTTGGATTTTGTATTATGATACAGAAGGCTAGTTATACCTTTAGTAATTTTGTTAGTACTAACTTTATTATT |

| | | | | | | | | | |
|---|---|---|---|---|---|---|---|---|---|
| | | | PBANKA_0804000 | 60S ribosomal protein L37, putative (RPL37) | Essential | GGTAAAGCTGGAAAAGGTAC | AGAAGAAATACAATTGGTAC | AAAAAAAATAATATTTATTTATGTTATTTAAAAAAAAAAATATATAATTTTACTAATAAATATTAATTAATTTATATAATTTTATTATTTAAAATATAAA | TTGTTATGTAGATATACCCCATCAAAAAATTACCAAAATATACTATATTTGCTTGTGACTTAATTTTTTGGGAATAACGATTTAGGCAAAACGAGAACAC |
| | | | PBANKA_0809200 | ribosomal protein L35, apicoplast, putative | Essential | CAACGTAGTTAAACATAGCC | CAAATAAATCGATTGCAAAA | ATTCCAGAAACATATAAAATAAATATATAAAAAGGAACAGAAAAAAATAAAAATATGTTTAATATATAAAAATGTGTACAGATATATTGGAACATGTGCTG | AATTATTTCGAAAGTTTTGTATCCAAATCATGACACTTTCTATTATCTCCCATATATAACCTTGTAAAATGTGTAAATAAAATATGAACATATCAATATT |
| | | | PBANKA_0703200 | ribosomal protein L21, apicoplast, putative | Essential | AAAATTGAGTAAAAGGCCTG | ATAGGATCAAAATCATCCAG | TTTTAAAAAAGAAATGAAAAACATTTAATAGTATATTTTCGTATAGTTAAAAATGAGAGAAACAAAAATATGTAAAAGGCTAGAATGAAAATGTGGAAAT | TGAAATATATATTATGTGAATAATTTTTTTTTTTTTTTTAGGTCTAGTTTTAGGCCTCCTTCTATGTTTTGTCTTTTTTTTCTGTGCAAAATTATAATTT |
| | | | PBANKA_0505400 | conserved Plasmodium protein, unknown function | Essential | TAGATTGAATGGATCAGAAG | GTTGGTACATCTGTATTGTC | AAAAAATAAAAGAGAGAAAAGCACGACAAGCCAAACAGTAACAATAATAGTAGTAATAGCTAGCTAAAAAAAAAAAAAAAAAACATATCAGAATAATAGA | GAAGTGTAGCTAGCTGATTTAGGAAAATAAATGCAGTTATTACCACATTTTCCTTATTCAAATAATTTTTTTTATTTTTTAATCCCGTAACTAGGTAGTA |
| | | | PBANKA_0812300 | spindle and kinetochore-associated protein 1, putative (SKA1) | Essential | CATTTGGCTTCCCAGATCTA | TAAGTCTTAATTGCTCATCA | TCAATTTGGCACGTTACTATTTTTTATTTTATCTTCAAATTGTATGTATTAATACTTTTTATTTTTAATCTTTTTATGTATTTATTACAATTTGATATTC | ATTTTTTAATATTTTTATTAATAATATTCATAAATGTAATAAAATTAAAACATTTCTGTTCTGAAAGGGAGGTTTTTTGTTTTGGGGTATTCTTTTGCGG |
| | | | PBANKA_1009800 | cytochrome c oxidase assembly protein COX15, putative (COX15) | Essential | GGATGGTGGATGGTTAAAAG | AATTGGTGCACTAATGCCAG | GTGATGAATATATGCTTGTGCACCTCTCTCTCTTTAATACAGGTATATACCCTTTTTTAACACAATTATTTTTAGGAAAAATTGAATAAAATAAGGAAGT | ATAGTTGACACTCCTTTCTTTCAAGTATATATACTCAAACATTAACATCGAAGCGTTAAAAATATATGCGTTTGCCAATTTGTAAACATATATTTTTTTA |
| | | | PBANKA_1136500 | conserved Plasmodium protein, | Essential | AAAATATGATAAAATACTGG | GTACCCATTGTAGATTTGAA | TTTTGTTTGCACCTTAACTTTTTTTATTGACAAACGTGCATGTTTTATAAGAAACCCTTTTTTTTACATCGATATAT | GTTAAAAAGGAAAGTGTGAATACATCATAAAAATTTGCGCGGAGAATTATAATGTATGGGAAGAAAATAATGACG |

| | | | | | | | | | |
|---|---|---|---|---|---|---|---|---|---|
| | | | | unknown function | | | | TGCTACAAAATACCAATAAGAAT | GAATTTTTTTACTACTTGCTAATTA |
| | | | PBANKA_0914900 | onserved protein, unknown function | Essential | TTTACGGTGAAGCGATAATT | ATGAACGCGTCATAATAGTT | TTGAATGCTGTATATAAGTGTAAATAAATACAAGTTTTTCGAGTTAGAGAGTATATATCAAACATGATCTACTATATATATCAGTCTTCAACCATAAAAA | TCACATATAGTTTGCAATACATTTTTGTGTATAATTTTTTCATGTTTACATTTTTGTGTGAAAAAGTAATGCGTGAAAATGTTCATACATATTTATATAG |
| | | | PBANKA_0817800 | conserved Plasmodium protein, unknown function | Essential | CGAAAGCGAAGAATCCCCAA | AAACAATTCCTGATAAAATG | TATAATTTTATATGGATGTTCTTCCATATTCCATAGTTGCTTCTCCATTGTTGGATATTTCGCGCTCTGGGGTATCATTTCCTTGCTATTTTCTTCTATT | TTTAAAAAAAGTTGAAAAAATTGAAAAATTGTAACAAATTATAGCAATTAAAATATGATAAATGTTTAATTTTGTAGTAATAAATCCTCAGTATAATATG |
| | | | PBANKA_1233900 | 50S ribosomal protein L14, mitochondrial, putative | Essential | GTTAGGTGTGCAGATAACAG | ATGCTTTATGTGGCCTTTGG | TACAAAATAGTGGTAAATATATATGCACACGCCTAAATACTCTTATATGTAAGGATGTAGCACATTAATTTTTTTCGAAACAAAAAAATTTTGAGTATACT | TAATCAAGATCTTAATTCACACACACATGAACAAGCATGCGTATATCCAATATATACATATATTACATGTGTTATAACTTTTTATTTTGGAATTCAAAGT |
| | | | PBANKA_0514900 | ookinete surface protein P28 | Slow growers | TTTCTAAGCCAAAATTTCCC | AAACCCCAAGCACCAGGTAC | ATTTATATTCTCATAATTTACGTAAAAAAACAAAACAATTTCATAAAATTATATCATAACAGTTATTTTAACAATTATTTTATATTAAATTTTCACGAAA | TATATTCAATTGTTATCGCATATTGTAGGAATATTTATACATATTTATATATAGAGACACAAAAAAAAAATTGAAGCAAATTTGACTTTTAAATAATTTA |
| | | | PBANKA_1362100 | tyrosine kinase-like protein, putative (TKL3) | Slow growers | TTTATAGGGGTAAAGGTATG | TAATGGATCTGCATTGTCAA | ATTTCGACAAATTTTTTCGAATAAAGAAAGTTTTTTTTATTATTATTTTATTTTTTTATTATTTTTTTTTTTTTTTAAGCATGCGTTTTATTTCTTAACT | CTTTTTAATAAACTTAAGTTATTCATATTTAATCCATATAAGCACACATTGCTTACATATTTTTATACATTTGGATTTCAATACTTATACTATTCAACTT |
| | | | PBANKA_1108400 | single-stranded DNA-binding protein, putative (SSB) | Slow growers | CCTATAGACAAAGATGAAGG | CCTCCTTCATCTTTGTCTAT | ATAAAATAAATATAACGTGATATTGGATAATTTCCATATGCATTATACATATATATGGATGTGCCATAGTTATAATTTGTGATAATCGTACAATAAAATA | TTTTTGCAGAAAAAAAACGTATTACCATGCAAGGTGAATATTTTTTTTGTTAAATTTAGCTAGCTAACTAGCTATTTCATATGAAAAATGATATACACAT |
| | | | PBANKA_1322400 | exonuclease V, | Slow growers | GTTCGAAATTATGTACTTAG | AAAGTGCCAAATTCGAAAAA | TTTTTTTAAGATTATAAATGCACACGCACACATATATATATACATTTTTTTGTTCGAG | ACAAAAGTTACAATTTGAATCATATTAAACTATAAAATCATAAAAATAATTTATCATTT |

| | | | | | | | | | |
|---|---|---|---|---|---|---|---|---|---|
| | | | | mitochondrial, putative | | | | ATATATATACCTTGCCACCTCTACATCAATAGGAATATAAAA | GCACAAATGTATTATCTCTATGTGCGCATTTATATATGCAC |
| | | | PBANKA_1426400 | mitochondrial carrier protein, putative | Slow growers | ACAGTTCCTTAACAAATGCG | GAATTACCACGCATTTGTTA | TAAAATTTTTACATATTATATAAGTTTAAAAAAAACAAAGCATCATAAACGGAAAACATAGAGAAACAATAAGTGCATGTATGCATAGTAAAACATGTCC | TTTACACGTTTCATCTTTTTGGCTAGTATATATATTTTTATATTTTCCAATTAAGTTACAAAATTATAATCTTCATTTTTTACGCCTATATTATGTTATT |
| | | | PBANKA_0314200 | calcium-dependent protein kinase 1 (CDPK1) | Dispensable | TCAGTGAAGAGAGGCTAAGG | ATGGAATGATGTCTTAGGGG | TAAAAAGCTATATGGTATAGCAAATATATTATTTTAGTTAGCCAGAAAAATGTCTTAGAATATTATTTTTCTTTTTCTCTCTTATTTCTATTCCTCTTTT | TTCATGTTAATTTAAATCACTCTGTCTAATTATGCTTGGGTAGTTAGGGATGACTCAATGGTTTATATATTTCTGTATGTGTATATATTTATACATGCAC |
| | | | PBANKA_0616700 | NIMA related kinase 4 (NEK4) | Dispensable | TCCCTTTGTTGAATGAAATG | ATTGTAATGAAGCATTGTAA | ATATTTTTTATCAATCATTTGTATATATATATATATATATATATATGCCTGTGTATGGACTGTAATATTATATTCTTTTTTTTTGTGGAAGAATAACATA | CTTTAGGAATTTCAAATTGTAAGGTTACAGGAAATTATATTAGTTAATAAAAAAAAATTTTGTGAGAATATGTATAATAAAGAATGAGCAATACACAATC |
| | | | PBANKA_0408200 | calcium-dependent protein kinase 3 (CDPK3) | Dispensable | ACACCCCGTTAAAACTTGAG | GAAGTTGACAAGAACAATGA | GCCTCATTAAAACAAGAGTCTACAACACTATAGGTTTACAAAAAACCAATAAATAATCATATGTTCCTGTATATTATAATTCTTTAATCAATAGTTCCAT | TTTTACGTATTAAACTATTTCCAAAATAAAAAATAAAAGTATATATTTAAAAAGGCAAGTGGACAAATTTAAAGAAGTAAATGAACAAATTTTAAAAAAT |
| | | | PBANKA_1305200 | serine/threonine protein kinase, putative | Dispensable | TCTATGAGACTTCTCACTTG | GTATGAGTTGAATTTGATGG | TATAAAATTATAGCATTAATATATAAGGGTCTTTCCTTGTATATTATTAACATACACGTAGAAAGGTATGAATATAATTTTTCCTTAGTTTTAATCCTAA | TTTAAAATCAATACTCCAAAAAGGAAAAGGAGGAACGATACAATAAAGACGCGAAAAAAAGCGAATTAAGGAATAAACATTAGAACAAAGTAGTAACTAT |
| | | | PBANKA_1414500 | glycogen synthase kinase-3 alpha, putative (GSK3alpha) | Dispensable | AGGGCACCGGAATTGCTTTG | AAAGGAGAAACAAAGTTTCT | ATATAGACACAAATAATAAATATAAACTTATTTATTTGTGCTATATGCATATATCGAAGAAAAATATAGTGATTAAGCTCATAAAGTATATAAATACAAA | AAAAAAAAAATAGGATGCATTAAAATATATAAAAAATAAAATATAAAGAGATAGACATATTGTAAGAAAAAAATACATATATATTAATTTGTATAAGAAT |

| | | | | | | | | | |
|---|---|---|---|---|---|---|---|---|---|
| | | | PBANKA_0926600 | protein GCN20, putative | Dispensable | ATATGTGGTGTTAATGGTAG | GCATTGAGTTTGTATAAAGG | TATGATAAATATACCAAAATAATATAATTATTGAAATAATATTCTCACAAAAATCGATATATATAAGGATTTATAATCACACATGGATATATGGGGAAAT | TTATATACAACTTTCATATATAAATTTATAGCCATTAATTCTATTTTTAATATATATGTTTGTGTAAATACATATGAAAACGTGGGCATAATTAAAAAAT |
| | | | PBANKA_1352600 | serine/threonine protein kinase, putative | Dispensable | TATGTAAGCATTTCCAATTG | TCCGGAAGTCCAACGCCTGA | TTATTCCTTCACGAATGTTGGATTGTTTCATTTACTTGGCTAGTTATTATTCAACTATTTTTTTTGTAATATAATATAATTAAAAAAATTTTCAAAAAAA | GAAACTTTCTTGACAATTATATGACTTAACTAGCGAAATTGTATTTTTATTTTATTATTTTTTTTTTGTTGTATTTACCTGCACGTACATATTTTATTT |
| | | | PBANKA_1421600 | calcium/calmodulin-dependent protein kinase, putative | Dispensable | GATCGAACCCTTGATAATAA | AATCTAACTTGAAATAATTC | TTTACTTGGTCATGTATTCTTTATTTTTTTTAATTTATTTTCCCGTTTGGATATAATATAGAATAAAGCATACACAAGCATGCCTTTGGAAAAAATTCCG | ATATTATAAGTATATTTGATACAAGCTTATTTATCACATATAAATTTAATATTTATTTATAATTAAATCAATATGACCATAATTATTTCCATTTAAAAAT |
| | | | PBANKA_0308500 | tyrosine kinase-like protein, putative (TKL1) | Dispensable | ACTTGAGCTATGATGTCTAG | TATTCGAGTGCTATTGCTTG | ATGAAACATGAGGAAAACATTATTGTTCTCCAAAAAATGTGAAAAAAGTTTAAAATTTGTAAACATACATGTCTGTCCTTTCTGAAAGGAGTGAAAACGC | TTTTGTTAAATTATATAATTCATTTGTGCTTGTTCAAACAGTTTTGAACCATTGCTTCTATCAATTACCATGCTTACGCATATGTATGTATATATATGCC |
| | | | PBANKA_0604400 | targeted glyoxalase II, putative (tGLO2) | Dispensable | TGGCTTGCCTATATGCGCAT | TACATTGCCTATGCGCATAT | TTCTATATTGCGAATGTATACAAAAGTGTGAAACAAGCAAGCAATTATGATATTTACACTTCGCATATGCTTGGTACATTCATTTAATTGGTTAAAACAG | TTTATAATATCATACCTTGAAATTGTTATATTAGTAGTATATGTGTGCAAACACATTCAAAATAATCAAACAATATATTAAATAAAATGTGGCATCCTTC |
| | | | PBANKA_0103700 | conserved Plasmodium protein, unknown function | Unstudied | GGCAGGAATAATTCCATCAG | AAAAGAAGAAAAGATGACAA | ATCATATCCCCTATAATACATATAATTTAATGTTTACCATTTAAAAAAATGCCACTATATTGATACTTTTAAAAGACGAATTTCATTGTAGAATAAAATG | GTTCCAAATTTATAATAACCATTTTGTTTTTTCCGTATATATGTGAGTAATACTATTATGCATACTTTTGTAAAGACATAACAATTCAATAAATAATTAT |
| | | | PBANKA_0111400 | conserved Plasmodium protein, unknown function | Unstudied | TTTGTTGTCGCGTTTCTTTC | GCTATTTGACGTCCCGATCT | CTATGAAATAATTTTAGATTTATTTCTGACAATAATGAATAAATATACTTTAAAAAAAATATATAAAGCGAATTAC | TTTCCCTCCTATATATTTTCAAATAGTATAATTTTGTTTTCAGTAATTCAAACTTTTAAGTAAAAAAATATATTTATAT |

| | | | | | | | | | |
|---|---|---|---|---|---|---|---|---|---|
| | | | | | | | | ATAAATTCTTTTCTCTTAGGCATT | ATCTATATAATAATTTAATAC |
| | | | PBANKA_0111500 | conserved Plasmodium protein, unknown function | Unstudied | GCTCATTGTTGATAAGACTA | GTTCCTGTACTTGTTTAAAA | AAACATAATATACCCGGTTTACTACAATAAAAATGTTTATTTATAAATGTTTACAAAATTGATATCATATTACTATTTATGAATAAAAAAAATACAAAAA | ATTATTTTTGTTAAATATTATAATCTAAATGTTGTATCATTTTGTTATTTTATGTAATGTAATATAAAATTATATTATTTTTTTATTTATTTGTTAAAGA |
| | | | PBANKA_0210800 | conserved Plasmodium protein, unknown function | Unstudied | TAATAAAACGCCATGTATTT | TCATATAACTCCTAAATACA | TTAAGGCTACAATATTGTTACAAAAAAATAAAAAAAAGATAAAACAATGTAATATTTACCACTTAGACCACAATTTAAAATAACGATAAATATGCAATAA | AATTGTATCAAACGTGAAAAAATTAATGAACATCTTTATGCATATATTTTTCATCATTCTACATAGTTGGGGCTGTTTATTTTTTTTTCATTTTTTCAAT |
| | | | PBANKA_0315700 | conserved Plasmodium protein, unknown function | Unstudied | ATATACTCGAAGGATATTCT | GGATATTCTTGGCGATTTAA | ATTTTAAAAAAACATAAATGCGAGTATCATTTTCTTATTATGTTTACGCAAACTTAATTTTAATTTTGTACTAATATTTCATTATTCTTATTAATAAACT | CATATTTATTTTACAAAATTAGCATATTTATAGGACTCCCATAACACCAATTAAAGACAAAATATTAATTAAAATTAATCGACATATTTTTTTTAAATTA |
| | | | PBANKA_0408800 | conserved Plasmodium protein, unknown function | Unstudied | CTGGGTAGTTGCACATATTT | ATTTGATGAGTAATAGGTGC | GTCCACATACATAATTTATAAAGGATTATTTCCAAATTATATTTTTTTGTTTTTATTCCACTATTTAAATGTTTTACTATAATATTTCTTCAGTTTTTTT | TTTATATATATGTTATGTAATAATGCAAAATTTTCGAAATACAATTAGATATGATTAATATATATAGGTGTGTATGATGTTTCGAGAAATATCAATGAAT |
| | | | PBANKA_0409000 | conserved Plasmodium protein, unknown function | Unstudied | CAAACTGCCAGATCAACTTC | TAATGCGATCCAAAATTCCA | ATAGACAATGAAGAAAAAAATGATAGAAAAAATATATGAAGAATGAAAAAATATATAAAAAAAGATAAAATATATAAATAATTTTTGCTTTAAAAATATG | TATACTTTAATAAAGAATGTATTTTTTTTTATTAATTCCAATTTTGTCTTTTTAGTTATTGTTTAATTTTTATCTATTTATGTTCCCAATATATATATAG |
| | | | PBANKA_0502900 | conserved Plasmodium protein, unknown function | Unstudied | ATTGTCTAGAAATATGTGTG | CATGTTCTCAAAACTAATGA | ATTAAATGGTATACAATATATATATATATATATATATATATATTATTTGTGTGTTATACAAATTTGAGCCAATTCAATATTAATGTAGTATTTTGAAAAA | ATATTGCGAGCAAATGTTAACAAATAAATAAAATAAATAAATACTAAGCACGTAGTTGAACATTTAGGAAAATAATTCCATATTCACTATTATTAATGTG |
| | | | PBANKA_0509600 | conserved Plasmodium protein, | Unstudied | TATGGAAATATGCTTATTTC | GTTTGCATCTGAATCGAGGC | GAATAAACACATAGGCATATATATATGTATGCATATTTTGTGTTTTTTATAATTAAA | CAAATCTGTGCATATCATTATACGGCTATGATTGTGGGAAAAAAAAATTTAATCGAA |

| | | | | | | | | | |
|---|---|---|---|---|---|---|---|---|---|
| | | | | unknown function | | | | TTTAATTTCGAGCATTTTAGTGTATTTTTTTTTGGAGATTAAA | CAATACACAATTAGTTAGTATATGCAATATATTTATTATTTCA |
| | | | PBANKA_0519100 | conserved Plasmodium protein, unknown function | Unstudied | TCAGTTTCGGGACTTACACC | GGTGGTGCTGGTGCTGTTCC | AAAAGTCAATAAATTAATAACAATATATTAATGGTAATATATTAAAAAAATAGGTTAAAAAATGATAAATATTTCATTTTTTTGACATTTTTTAGAACAT | TTTTATATAATTTATAGTAATACGTCACAAAAATTTACAATAAATTAAACAAGAGGATATAATATATTTATAATAATATTCTGTTTTCTAAATGCGAACA |
| | | | PBANKA_0519200 | conserved Plasmodium protein, unknown function | Unstudied | ATTCTAGGCAAGATGCATCA | AATCTAGACAAGATTCATCA | ACAATTGAGATGCTATATTTAATTACTAAAATAAATGAAATAACCTTAAAATTCTTCATTTATCGCCTGTTGTTATATATATGTATATGCATATCCCCTC | TGTGAATTTATGTTTAAAAAAATAAAATAAAATAAAATAACAATAGTGGTTGTAATACTAGTAAAGGTAAAATAACTATTTGTGTGCAAAAATTATAATT |
| | | | PBANKA_0519400 | conserved Plasmodium protein, unknown function | Unstudied | TCAGTTGAAAACCCTTCAGG | TCTGATCCAGGGACGCCTAA | TTTAATGATCACTTCGTAACGTGACCAATATAAATATATATAGGTACATACTAACAAGCCTTCTTACTTTTAAATTTCTGAGTTTGCACTTTTCCCCTTA | TTTTTTATATATTTAACAATAAATAACAAAAAGTTGTATGCATGTATGTATATATATTATTTCGCAAAAATGGCTATGCATAGTACATTACACATTTATT |
| | | | PBANKA_0523700 | conserved Plasmodium protein, unknown function | Unstudied | TATCATCCACAGACGAAATA | GAAAGATATTCTAGAGAATG | AAATAAAATGATTAAAAAATATCAATTTTTATATAATATTAAAAAAATAACGAATGACAAATGAATATATATAAAGTATATGCAAATAATCAATTTAAAA | AAAATATTTCATAAATCCCCTTTAAAATATCACCTACCACATATATAGAATAAATTATTTGGAAAAAAAATAAAATATTTAATAGACATACATAATACGC |
| | | | PBANKA_0524100 | conserved Plasmodium protein, unknown function | Unstudied | TGATCGGGCCCTATTACTTC | TCGTTATATCACAACTCAAG | CTACCACAAAAATATAAACTAACAGTTAAACAATATATGACAATATGTCATTTTGTATACAAATGAATTATAATAATTAGGTATATTTTATATTTGTATA | TATAATAGTAAGTAAATATTTATTTGACCAAATTCGTCTGTATTTTTTTATATATCCATAATTCTTCATTAAAATTAAAGGGATTTTCAAAAAAAAAATG |
| | | | PBANKA_0603650 | conserved Plasmodium protein, unknown function | Unstudied | AATGGTGTAAATATTAACCG | AGAGGAGCTAACACGCCGAA | AATACACAATAGACTATAAATTTATTAAAAATGTATAATTCACTAATGTAGTAGCATAATATACAATTTTATATATTTCCTATTTTTATTTTATTATATT | TTTATACTCAAAAAAATTAAAATAAAATAAATTTTTGGCAACTTATACACATTTCCCCATCACTTATGTTACAATACTTTATAATTTATTATTTGTTAGT |

| | | | | | | | | | |
|---|---|---|---|---|---|---|---|---|---|
| | | | PBANKA_0619000 | conserved Plasmodium protein, unknown function | Unstudied | GTTTGTTTCGATCCTTCAGG | AATCTAGAAAATGAGAATTC | TTTGAGGATAATAAAAAAGACGAAAATGGTGCTTATATAAAATTGCCAAAGGCATATTGGTACTTGCGATATTATGCCACACATTTTATTTGCATTAAAT | CATTTTATTTTATTTGGAAAATTGAGGATTTTATTTATTTTTTTTTCTTATCAAATATTTTAATCATTAAATGCCATAATTACACCTGGTTGAAAATTT |
| | | | PBANKA_0705800 | conserved Plasmodium protein, unknown function | Unstudied | GTTGAGGAGCAGCTTGGTAT | TTTAGGGGTAGATATAGAAG | TATTATTAAAAATAATAATTTAAAAAATTTGTAGTAAGCCTATTGTTTTCCCAAACTATGTACGATTAAGAAAGCTTGCTAACCAACAATAAATAATAAA | ATGTTGATAATACAACATAATATGGAAAATGGGAAAAATATCAAATATAAAAAAATAAGAAATGTAGGAAAGGAAAATACCGAAAAAAGGCTCAACATAG |
| | | | PBANKA_0709800 | conserved Plasmodium protein, unknown function | Unstudied | AAAGTGCGAGTATTAATTAT | GGACAGAAAGTTGAGTAAAG | TATATAGACAATATATAAAATTATTAAATAGCTGTTCCACTTGTACTTGTTACATATTTATAATTATATTTACAGTTGTACAATCATTGTGAGTGTTAAA | CAATAACAATTGTATGGAAGGAAGTATAAAAGATATCATTATTTTTCATATTTTAAAATATCGATATCGTTAAATTGTTAAAAAATGAAATAATAATAGC |
| | | | PBANKA_0715300 | conserved Plasmodium protein, unknown function | Unstudied | ATTGGATCGATAATTTCTCC | TCCGGGATTGGCATTACATT | TATTATTTATTTTCTGTTTTATTAAACGGCTGCATTCCTTGACATACAGTATTGAGATCAAACACTACCTCTCCTTCCATATTGTACAAACTAATATATA | AAAAAAATATATAGCCTTAATAATAATAAAAATTAAAAAAATACTAAATACATAAGTATGAACAATTCGTATTTTATCTGTTTGATGGTTTGGCCCATTC |
| | | | PBANKA_0812900 | conserved Plasmodium protein, unknown function | Unstudied | ACAGGCATACCACTTATAGT | CCATATGAGCCAACTATAAG | CGTGATATTATAAGGGCGTTTTAAAAAGCAGCATAGTAACATAGCTTTATTATTTTGTTGAATATTATTAAATGTACTATGGGGTGAATTATTCAATTAA | AAAAAAACAATGATATAATACACACATTTCCTCACGTTTTCTTTTTGTTCTTTTCTCGATAAACCAAGAGTTAGCAATAGTAATATATGCTATATTTGTA |
| | | | PBANKA_0817400 | conserved Plasmodium protein, unknown function | Unstudied | TTCCGCGAAAAGTATTATAA | AATATTTAAATAATGACTAG | ATAAATGCTGTAAGGAATCCCAGAAAAAAAAAGGAAAGAAATAAAAAAAAAGCATATACACTTGGATTTTATATAGGGCATATCGGATTAATTGATAAAACT | TTTATTAAGCCAAGCTCGCTTTTTTTCTTTTTTTTATAAATTGAGGAGAGAAAAAAATTTTTTTTTTCTCTTTCCCCTTTTCTTTCGTTTCTTTTCGGCC |
| | | | PBANKA_0821200 | conserved Plasmodium protein, unknown function | Unstudied | CCTATTTCATGCTCCATCTA | CCATAGATGGAGCATGAAAT | AGCAAATATCACAAAATTATTATTTTTTTCTCTTTAATTATTTATTTAAAATAAAAAATGACAGACAAAAAAAGGAA | ATTGATGGTTTATATTTTTTTAATTCTTAGTATATCATTAATAATGAATATAAATTCAAATTTGAATACTTGTCAAAGA |

| | | | | | | | | | |
|---|---|---|---|---|---|---|---|---|---|
| | | | | | | | | ACATAAAACAATTTTATCAAATA | ACAAATAAAAACAACAGTGAT |
| | | | PBANKA_0915400 | conserved Plasmodium protein, unknown function | Unstudied | ATTATGTTATTAATACTTAA | TAACATAATTAAAGCAAATA | AAAGTACTAAGAGGTTACTTCAAAATATTTTAATTTAAAAGGATTATTAAAATATAGTCCAAAGATTATTATTCACAGTTATAATATATTTTTGACCAAA | GGCCATCATTTTTTAATTGGGAAAGAACAAAAAATATAATAATGTAGAATTAACTATATTATTGTTATATACATCTGTATATTTGCAATTATTATTTTCT |
| | | | PBANKA_0930600 | conserved Plasmodium protein, unknown function | Unstudied | GGTGCGGGAATAGTTATTGG | TCTCGAAGGTTAGTTGCTGT | ACAAACACTATATATTTATACATTATATAAACAAAAGTTACAAAACATGATATATATAATAATTTTTTTTAAAATATAGATAATCAAAATAGTTAAAAAT | CACATCATCATTAAATTTGTCAGTTTTAAATATCAAAAAGTAAAACAAACAAAGCAAAGAAAACTGTTTAATTTTGAAAAACCAATGCAGCTTTCAGGGT |
| | | | PBANKA_1015100 | conserved Plasmodium protein, unknown function | Unstudied | TATTTGCTTGGATATTATCG | TATGAATTTCGAAAGTTTGG | AATATCCACATTGTTTCAAAATACAACATATTTATAGGCATATATTATTAATAAGCCCAAAGGGAGATATAATAACAATTCTATAAATTTTTTAACGACG | GTTATTTTTTTTTATTTTATTTTATTTTTTTTCTTTGTATTTTTTTTGTTTTTATTAAATAATTTAACTTTATTTTTATGTACAAAACGTATATTATATT |
| | | | PBANKA_1016900 | conserved Plasmodium protein, unknown function | Unstudied | CAAAACCAAGTACACATTTG | TTGGCTAAAAGCTTGTGAAA | TTTCCTGTTTTTCTCCTTTTTAACTTAAGGTTGAAACGCTTTTAATAATAGCTTAAAAAAATAATAAATTCCTGAATTTTATATGCATCAAAGTTCATTT | TGCACTTTACTTCCAAGGTTTAATAACAATAAAAATGTAAAAAAAATATATTGCTATATATATGGTTATATTTGTGTATTGTTGTAGTGGAAATAGTGAG |
| | | | PBANKA_1017200 | conserved Plasmodium protein, unknown function | Unstudied | AATGTGCTACAATATTTAGC | TATGTAGCATCGATATCGTA | ATATATTGCATGCTGGGATATATTGTTTAAAAAAAATTAAAAAGAACTTTAAATATCCACAAAGGAGTAAAATTGCAAATATTGGGCTTTATAACTGAAC | TATGTAGAAAAAAAAACTTCAAAATTTGTGATATTTAAAACAATGGGTAATTATAAACGAGGAAAAAGAAGACTAAAAAATGCTTTAATTATTATTTGTT |
| | | | PBANKA_1035000 | conserved Plasmodium protein, unknown function | Unstudied | TTGGTGTTTAAGGATTTCGT | CGAATGCAACTTGGTGTTTA | AAAGCCTTGGATAAGGTTTTTCAAATGTGTGTATATTTCATAACAATGTTTTCCGTTGGTTATTCATGGGGTTTTCAATTTTACAACTTATCCTGACTGT | TATTCACACATCAATGGGGTTTATTTTTATTTTTTATTTTTTTAAAAAAAGACTGATCAAATTAAATGGTTATTATAAAACTATGTATATTTTATTTTTA |
| | | | PBANKA_1037300 | conserved Plasmodium protein, | Unstudied | AATATATGTTCGTTAATTCC | CTCTCAAGAAGCTATTCATA | TTATATAGTTTCATTAAATTTTGTTTATTGTTATTTAACATATATTATACAAAAAGAAA | ACATGAAAGAAAACACCCATTTTTATGCTATTAATATACAATATAGACTAAATCATGA |

| | | | | | | | | | |
|---|---|---|---|---|---|---|---|---|---|
| | | | | unknown function | | | | TATTTTTAAAAAATATACTCTAAAAATTTACAATCACTAAA | GTATACATAAATATATATACACCGATTTTATACTGACAAAGT |
| | | | PBANKA_1120000 | conserved Plasmodium protein, unknown function | Unstudied | CAAACACGGAGAAATAGATG | GCAAACTGTTAAAACCGAAG | ACAAAATGAACTAATAAACTTATAAAAATAAATAATACATAAGAAGTGTAAAAACAAGCGTCATTGACAGAAAAAAAAAAAAAATTGGGGCTAGTAGGTT | TCCCCATATGTCTTTTTCATTGTTTAAGTTATTAATATTTTCACATTCTATTCCCTCGATTTGTTTGCTTATTAATTTATGCAAATGTATACATCATTAC |
| | | | PBANKA_1134700 | conserved Plasmodium protein, unknown function | Unstudied | CTGCCCATGGATTAGCTCAA | TATCACTGCTACAACACATG | TTTTTTGTAACATATATAATATATTTGTATGTGCTTATATTATGCAAGTTTTTTTCCATTTATTGTAATAATATTTTTTTTCATCCTCCACTTAATATTA | CTATAAAAAATATTAAATAACATTACTTTTCATAAATACTGAAATTAAACTGCACATTTGGGAATAATTGTATCATATAAAATCAATAGCATTACTTCAG |
| | | | PBANKA_1135300 | conserved Plasmodium protein, unknown function | Unstudied | TTTCGACGATAGTATCTTCT | TCTTGGTCGCTACCTGAGTC | TTTAATAATCAATATACAAACCACTAAATTATTCACACACATCTATATGCTTACTTATTATAAAATATTTAGAGAGATATAGATACGGATAAAATACATA | ATAAAATAAAACAAATTAATGTGAAAATAGTTGTAAGAACAAAACTGCTTATTTAGTTATATATTCATATATTGTATATTTGTTTTAAATGACAAATAAT |
| | | | PBANKA_1201300 | conserved Plasmodium protein, unknown function | Unstudied | AAAATCGAGGGTGTTAAAAG | GGGTGTTAAAAGTGGCGATC | ATAAATAAATAAATAAATAAATAAATAAATAAATAAATATAATAAATAGTAACTAAATAGAATATATATAATTAGATTGTGGAAATTGTTTTTGATGTTC | TTTTTTTAACAGATATCAATAAGTTAATTTTATTTTATATTTTTATATGCTAAAATAAAGAATTATTTTTATTTTTCATTATATATTTATATTATTTTTT |
| | | | PBANKA_1206000 | conserved Plasmodium protein, unknown function | Unstudied | TATATAGAAGGATTGACTGG | TCGGGCCAATATTATATTAG | TTAGCAATTTCGCAGTAAATACATGTACACACACACATTCCTATAATTCAACGTTTTATAAAAAAACAAATAGAAGTATATACAAAGGGAAAAACAAAATA | TTAATTTGATAAACATCATTCTATTTATTTCCCAATCATTTATAAGTTATGCTTTATTTTATTAAATTATTATTATATTTGTCTTTATACTTTTAGTGTT |
| | | | PBANKA_1211700 | conserved Plasmodium protein, unknown function | Unstudied | TAATTAGTCTTCGTATCTTG | AATGGAACATACGAAACGAA | GTTAAATATATTGTGTGTATACATATGCGATTGTGTGAACTAGTGGGGTTGTAGTAAAATATAAATATATTAAAGGAATAAAAAAAGTTAAAAAGAAAAA | AATTGCTAATAATAAATAAACGGGCATCATGATGTTTATGTACTTATAAATCGTTTTTCACTTCGTCATTTTTTTTTGGAATTTTTAAACCTATATAAAT |

| | | | | | | | | |
|---|---|---|---|---|---|---|---|---|
| | | | PBANKA_1237100 | conserved Plasmodium protein, unknown function | Unstudied | ATAGTGGCAGCTAATCCTAG | TGTTTGTAGTGGTTTATTCG | GATGAATATCCTAAGTTCATTCGATTTTAAATAAACTCTAAATATGCAATTACATATATGCAAAATATTTTGTCCATTCTTCTTCTACTATTATATATAC | TTTTATTTTTAAAAATTATCCCAATTATATTGTCTTCCATAAATAACACATGGTATAGTTGCATATATGTATTTTTTTGCTACACAAATTTTACACGGAA |
| | | | PBANKA_1321900 | conserved Plasmodium protein, unknown function | Unstudied | AAATGATACTATACCTACAG | GAGCAGGACATAAAACCCAA | TGAAAAACACATTAAGCATAAAATATAAAAGATATGTAAACTCCAATGAATAATATACGTCTTTTAATGAATTTTGTTTTAATTTTTTTTTATAATTTATT | TTCATTAAGGATAAAGATTGTGGCTTTGTAATTTATATATTTTTTTTAAACACAATATATGTATGTGTTTGTTAATTAAATTATAAAAAATTATTTGTTT |
| | | | PBANKA_1327080 | conserved Plasmodium protein, unknown function | Unstudied | CAATGCATGCTAAATGTGTT | TATGAAATTAAATGAAATTG | CAATATTTATACTATAATAATTTCCTAATTTTGACATTACTATATTGCAGATATATTCACATAATAATTGTAATTGTATAAAATATCGAATGTATTCTGT | TTTTATATGAGTTATAAAGATTTATGAATAATTATAAACTATTTGGATAGGTGTGCATATATAATATAATAATATTCCCTATAAATTCACGCTTTTTTAT |
| | | | PBANKA_1329900 | conserved Plasmodium protein, unknown function | Unstudied | ATGAAATGGTGCCTTCACAA | TATTGAAGGAACAATAATTC | TAAAAAAAAAAAAAAACATATAATTAGAAAACCTATGTGTAAATAAAAAATTATAAATTTAAATTATTTTACATTTCTTTCATTTTGTTTTTTTATAAAAT | TATGTGTGTATTTAATATTAATTAATGATTTAAGTATTTATTTTTTATGTTACAAAAATATTAAGCTAGTATAATGAATACAATGACAATCATAAAATAA |
| | | | PBANKA_1345100 | conserved Plasmodium protein, unknown function | Unstudied | GAAAGCCAAAGAAATGATGA | TTTGTAACCTATAAAAGGTG | AAATGTTGTATAAATATATGTACATATCTTTGCACACATGTATGTCGACCATATAGTTAGTTTTACACTCTAAATATATTTCTTTATTTTTAAATTTTAT | TATTTTATGAACAGTTCAGAAAAAAAAATACAATTATCCCATTTTTATGCTTCGCATTAATTGTTTAAAAGGCAAGCAAAAAAGTCATATATACATATTT |
| | | | PBANKA_1354800 | conserved Plasmodium protein, unknown function | Unstudied | ATCTTATTAAACATCACACG | AACAGAATTAGGCTATAATA | ATTTGAATGAAAGAATAAAAAGTTTTTAAATATATTAAAATTTTTATTATTCCATTTTTTTTTTACTTTGTTTTTAGATTATCATAAATATGCCATAAAT | TATTATTCATCCCAACAAACGCATATTATATACATCCATATATTTTTGCAAAAAATATATATAGTTTATTGATATTGTCTTATTACTCATTCAGATATTA |
| | | | PBANKA_1360200 | conserved Plasmodium protein, unknown function | Unstudied | AGTGAAGAAGCAAAGTATAC | GACGATGTGCTAAGTAGCAA | ATACCCTTTAACATAATTATCAATAATGTGTCGAGTGTTTTACTAAAATATTACATAACATAAATATAAAAACAAA | TTTTCAAAAATACTGGAATTATAAATATACCAAAAAAATAGTTAAAAACAAAAATATATTTATAATATGTGTATGCTTATTTTTATGAATAAAGCCTTTA |

| | | | | | | | | | |
|---|---|---|---|---|---|---|---|---|---|
| | | | | | | | | ACAAAAAAAATATGCATTGAGATAT | |
| | | | PBANKA_1408700 | conserved Plasmodium protein, unknown function | Unstudied | CCTTTCGCTTGGTATGCATC | AGTTTAGTATGCCTTTCGCT | ATTTATTTTTTAATCATTATAATTTTATAAAAAATATATTATAATAGGGCATATAAGTATTTACAGCTTTTTTACTATAGTTACTGGAATAAAAAAACGC | TTCGACGTGAATTTCTATTCATTTTATATATCTGTTAGGTCATTTTCATAAATTTCGATTTTACGTTATTTTCAGCGTGACTGCTGTTGTTATATATAAT |
| | | | PBANKA_1436400 | conserved Plasmodium protein, unknown function | Unstudied | TAAACCCTCAAGGTCACCTG | ACAATACAAGTAAACCCTCA | AAAAACGAAAAAAATAATAATCAAATAAAATAAATGCAATATATAGCGTAAAAAAAATAATAAATAAGGATATGTTTCGTTGCAGTTCAATATTACTGCC | TTTTTATTAATTTGGGAATGGAAGAATATATATTATTAAAAAATTGTATACAATGTATTTTTACGCGTTTTTTTTGTTTATAATTATATTGTGACTTGTT |
| | | | PBANKA_1437100 | conserved Plasmodium protein, unknown function | Unstudied | CATCTGATTCATATGGGTTC | TGCAGGTGTATCAACATATG | AAAAAAATGAGGTGATATTAGAATTATATGCATATATTATAAAGTTATATATATAATTTTTGCAATATATAGATATGTGTATAGATTGAGAAAATAGCAA | AATTTATCTGAAAGCGATGGCTGAAGTTTGTACCTAAAAAACATTTCGACAAAAAGTGGAAAAATAAAATGACTAGTGATGAAAAAAATTGAGCAAATTA |
| | | | PBANKA_1437900 | conserved Plasmodium protein, unknown function | Unstudied | CCAAAATCGTCAAACTCCGT | ATCGACACAACCAATGTTCA | ATTTTTGTACAAAGTACAAGCTAATTTTTTTTTAATATTATGCATATATATAATTATAAGGGGGAAAGGCAAAGTTTAAATAAAGCTCAATCAAGTTAAA | TTATTTTCCTATTCGGAATTTTTTTTATTCATTTTTTTGCGGTATTTATATTCAAGACTTGCGCTATAATTCATTTAAAAATTTGCACACACAAAAAAAA |
| | | | PBANKA_1452500 | conserved Plasmodium protein, unknown function | Unstudied | GCTACTTCTGATTCTTCTGC | AGAAGTAGAAGCTGAAGTAC | TAAAATAAGATGCATCACATATAGAGAATACATATACAACGATCACTTTAAAAAAAATTAAAGAACATGCATTGTTAATATAAAATTATTCTTTTACTTT | TTTTTTTTTTTTTTTTTTTTTAAATTACAAATATGATATTTCTACTAATAAACATATATATTTATTTGATTTTTTTCTAAAATATTATATTTAAAAAAAA |
| | | | PBANKA_1464000 | conserved Plasmodium protein, unknown function | Unstudied | CTTGGAATAATTTGTGCTTG | ACTAGTCATAATAATTTCAA | CTTATTTTTAACTTTCAAAATATTTTATAAAAAATTAACAAATGAAACATACCCCCAAAAAAATATATAAAAAACAGATAAAAAAATTACAAATAATATAA | TTATTATTTTAATACTAATTCATTCAAAGTTTAAGTATAAAATGCAATGATACACAATTACCCATCTAATATAAATTATGCATAAATTCAATTTTAATAT |
| | | | PBANKA_1234100 | protoheme IX farnesyltransfe | Unstudied | TTTGGATATTTCATGCTAGG | GCTGCACATCCCATTAGCAT | ATAAACCGTGATAATTATAAGCAAAACACATACTACGCTAAAATATAAAGTTTTAA | TCTTTTCTCATTTTTATATAATTCTTATCTCGTTAATATATCCTCCTTTTTTTGTATATA |

| | | | | | | | | | |
|---|---|---|---|---|---|---|---|---|---|
| | | | | rase, putative (COX10) | | | | TTAATATGGTTTTATGAGTTCATTTTTCTGTTTGTATTCGCATA | TAATATATCCATATAAACACATGTTCATATAAATTCGTAA |
| | | | PBANKA_0605000 | thiosulfate sulfurtransferase, putative (TUM1) | Unstudied | TGTTTGATTCGCTCTTTATC | TATATTCGGAAAAACTTCCA | ATAAAAAATAACAATAACAAATAAAGTATTTTGAAAAAATATAGTTAAAGATGCAAGTACATTGTCTTGCTCATACGATAAGTTAAAGGAATATATATTT | ACTGCATTATTAGTATATATATATTGTTTTAGTAATTATAAATTAGTTTTGTTTTCGCAGAAAATATATCACATATATGCATATATTCATATATTGAAAG |
| | | | PBANKA_0514600 | tRNA N6-adenosine threonylcarbamoyltransferase, putative (KAE1) | Unstudied | GAAGGAAGCGCTAATAAGTT | GGTGCTATGATTGCTTATAC | ATTGAGCTAGAAGTATACGTTTCAAAATAAAAGCGGAATAATAGATATATAAGGTCCTATACACAAGCCTAAACACATACACATATGCTAGTTTCCAAAA | TGTCTTTAAAAAAAAGATACGGCATTATTTTAGTTACTATTCCCATAAGGATAACCATTTAATGAAATCACAATTTGTGATATATAATTTTTTGCTATTT |
| | | | PBANKA_1140700 | ferrochelatase (FC) | Unstudied | CAGTGCATTCTGCTGATTGG | TATATATAAAGGCTCATTCG | TTGTTTTGTCATTTGCTTGTGTTTTCTAAAAATCGTATATATTACACATATTTGTTGAATTGTATCGAATTTATAGTTTGATATACTTTTATATTTAATA | AATTATTATAAAATTCTTTAACAAGAATAAATCAAAACATCAAAAAAAATTAAATATTCTCAAATTAATATTAACCATAACGAATATTTTTATATTTTTT |
| | | | PBANKA_0306400 | 2C-methyl-D-erythritol 2,C4-cyclodiphosphate synthase, putative (IspF) | Unstudied | GGTGCAGGTTCATATGATTT | AATGGAATGCGAATAGGGCA | AATTCACATTATTTGGATGAAATAAACTGCTTAGGTTTTGTTTCATTTATTATTTTATATATTTTTATTTGGTCGCATTAAATTTTTAAACACAAAAACG | TCTTTAAAAAGATAACTAATTATTTGGAAAAATCTATAGAAAATATATATATATATATATATATATCACGTTTCACGAAAATATGTAAAATAAAAGAATA |
| | | | PBANKA_1111400 | inositol-3-phosphate synthase, putative (INO1) | Unstudied | AGATGCTGATTATTGTAAAG | TAACTGATATTATAGTATTA | AACACACAAATTAGGCTACGCGATATAAATCGTTGCAAATATAAAACAACTATTCGCATAATAATTCTGTATCTGTTTATTTTCGTCAATATATTTATAA | ATGTAATGTGTTTTATATGAATATAACCTTGTATACATATAGAATACTAGTAGAAACATCCATTGATTCTTAAGTTAATAGTTTACATTATTTTATTTCT |
| | | | PBANKA_0828100 | fumarate hydratase, putative (FH) | Unstudied | GCTGGGCCTGCATAATATAT | GGAATAGGTGCACAATTTGG | AAAGATAACGAATAAAAAATCTTGGCCCAATTTTATTTGTTTGAAAATGTTTAAAACTTATGAGTTTTTTTTTCCAACAATGTACATATTTTTGGAAAAA | TGTTGAGTTCAAAAATTTACAAATTTATTAAATAAATTTCACACAAAATCCCTGTGATGTTAATGCTAGAAAAATATTTTTTATTCACAAAGCACAAACT |

| | | | | | | | | | |
|---|---|---|---|---|---|---|---|---|---|
| | | | PBANKA_1344400 | AMP deaminase (AMPD) | Unstudied | CATCAGAATCGGGATCCCAA | AAACATCAAAGGATCATCAG | ATAAATTCGTGTTATTTATGTACTGCTCTAGTAGTTGAAATAACTTGCACCCCATTTTATAAAAATTCAAATTGTCACTGAATTATAAAAATGTGAACAT | TATTTTTATATATTTATAAGTATATAAATTTTTTTTAAAGACATATAAACATATATATTTATTATCAATTTGACGTTTATCGATTTGTTGGCTCATATCG |
| | | | PBANKA_0719300 | bifunctional dihydrofolate reductase-thymidylate synthase, putative (DHFR-TS) | Unstudied | TAAGGGAATTGGAAATGCGG | TATGGGAAGCTAATGGAACA | GCATGTGCATGCACAAAAAAAAATATGCACACAACATACACATTTTTACAGTTATAAATACAATCAATTGGTATAAATATATAAATAAGAAAAACGAACA | TTTGTAACATTTAGGTGTGTATTTATATATATATAAGCAATATACAAATGAAAATACTATTATAAAACGAAAACTCAAAAAATGTTGAAACAAATAGTCA |
| | | | PBANKA_1127700 | nicotinate phosphoribosyltransferase, putative (NAPRT) | Unstudied | ATTGGTATACGAATCGATTC | GACCTAATGATGGCTGTGAT | CCCAACATTGACATTTCTTAATTTTCCGATATAATAAAAAAAACTAAATTCCGTTATAATATATATATATATATATATATATATATATACAATAAGAACAAA | ACAATATTCGCTTATAAAAATGGCGAAACTTCAAAGATTCTTTCCATTTTACTAGTATAATATCACTATATGCATGGACTTTGTTGAATGAATAATTTTT |
| | | | PBANKA_1435100 | ribulose-phosphate 3-epimerase, putative | Unstudied | GGAGGGCCAAATGACAAATT | GACAGTAGAACCTGGATTTG | AATGCACAAATTTATAAACATTTTTTTGGATAATATATATATGTATGTGTACTACATGCATCCATGCTTATTTTATGCGTCTTAGAAAAACAATAGATAA | CTTTAATATTAATTTGGAGTTATATTTTTTCCCCTTATACTAAATAGCTAGATTTTTATTCAAAAAAAAAACATAAAATATTTTCCTTTGATTTGATGAA |
| | | | PBANKA_1416700 | glycerol-3-phosphate 1-O-acyltransferase (G3PAT) | Unstudied | ACAGTGAACGGGCGTGACAG | ATATGGGTAGCACCAAGTGG | TTTAAAAATAGGATGATTATTTTTTGTCATAGTCCATTTCCCTTCATAAAATTGAAAGAAATTAATAAAAGTAATAAATTCAATTACATTGATAAAATAG | CATTTTTAGGAATCACAATATTAACAATGTTATAGGTCAAACTTTAAGGCACGATTCAAGATGATTACTTGTTAACATATATATATATCGTATAAATACG |
| | | | PBANKA_0304000 | pentafunctional AROM polypeptide, putative (AROM) | Unstudied | TTGCTGAACTGATACTAAGG | TTACGGGAAATATAGACAAA | TGATTTATTATTAGTATAACAATTTGTTTTGTTTTTAAAATGATTTATTTAAAATCGATTGAAATAATTTTTTAAAACACATAAAAATAAAAAAATATAA | AGTTAAAATAAAGCATCAAAGGTATATAACAATATTTAACTAAATTTAATTATTTATTTTTATAATAAAAAAATTATTTTATATATGGATACACATTACT |
| | | | PBANKA_0306200 | patatin-like phospholipase 1, putative (PATPL1) | Unstudied | ATTCTATCTCTTGACAGTGG | GTACGTACTCATATGGCTCT | TAAAGAAAAGCGTCTTTCTTTCCTTTCTTTATTATTATTTTTTTTTTTTTTACAAACTTTTTAATTTTTCACAATTTAA | GTACATATATGCGTATGTGGCTATTAGTTTGTTTGCTTTTCTTTTTATTTGAAAGTAATATATAAAATGTGTACATATA |

| | | | | | | | | | |
|---|---|---|---|---|---|---|---|---|---|
| | | | | | | | | TGATATAAAAAAAAAAAAAAA | AAGTATTTATTTTGGGTGTGA |
| | | | PBANKA_1360100 | nucleoside transporter 1 (NT1) | Unstudied | CTAGTTGGTAACAACACTCT | AATCATCGCAAAAATCGGCA | ATATATTTCCCAAATATTATTTGAGAAAAAATATAAACATTTTGCGAATTTTAATATATTTATTTAATTATTTTTTTCTATTTTTCCATAATAAGTCAAG | AAAAAACAATAAAGCATACAGTATAGCATACAATTGATGTATATTTATTTTTACATATGTGATCATATTATGTTTAAATAAATCAATGTGTGCCTCTAAT |
| | | | PBANKA_0824700 | thioredoxin reductase, putative (TRXR) | Unstudied | TGAGTGACTTCACCTGCATT | CAAGGTGTAACATTTATGTG | AAACAAGCATACATTAATTTTTCTTTTTCAAACATGCCCTCATTTTTATGCACAAAATAGCTTTTTATTCGGATTAAAAAAAAAAAAAAAAAAAAAAATT | ACAAAATTTGATATAAATAGGACCTATATATTTTTTCATAATTTATATATTATTATTCTATATTTTATTACATCTAAAATATCTTTAAAAAAGTGAAACA |
| | | | PBANKA_1127600 | glucosamine 6-phosphate N-acetyltransferase, putative (GNA1) | Unstudied | TATGAATTCAGTTAGTAAGG | CTATCACAATCGTTTATTTC | ACATTTGCTTTTAATGAAATGTGATTTATATTTCTATATAGGTATATAATATTTTTTTCATACGGAATTTATATGTAAAAAGAAACTTAATTTTTAAAAA | TAAGAAAATACGAAAAAAAATACATAAAATAATAAAAAAGGCACAAATAAATATTAATACAATATTTCGACATTTGGAAGCCCACAAAAAATATTTCTCT |
| | | | PBANKA_1027000 | serine palmitoyltransferase, putative (SPT) | Unstudied | CATGGACCACGAATGTTAGG | AATTGGTATACAAACATCCG | TTTATATTATGCTTATATATATATATATGTTTTTTATTCAAGCATGTATTAATATATTTTTTTCTACTTCTAATACAATCCTAAAATATATAAAATATAA | TATTACTATTTCTTGTATTATATCGTGATTTACCTACTTCTTGATATCCTTTATCTAATTTTTTATCTAGAATTAAAAATCACGCTTTCTATTTTTATTC |
| | | | PBANKA_1028900 | diphthamide biosynthesis protein 1, putative (DPH1) | Unstudied | TTAGGGGATGTTACATATGG | AATTGGCTTTAATTCTGTTG | ATATAGATTTATGGGAGAAAAAACATTCCCATAAAATAAACAAAATTGGAAAACGGCATTGAATAAATTTTGTTGCTCATAGTTTACCAGAAATTAAAAA | TGATTTATATGCGATAGAAATGGATAGAATGGCCCCCCAAAATATTATAGTTTATGTTTATTATATTTTTTTATTTTCATAAATACATATGCATATGTAA |
| | | | PBANKA_1207300 | diphthine methyl ester synthase, putative (DPH5) | Unstudied | TATCATTGGACTAGGGTTAG | ACTAGGGTTAGGGGATGAAA | AATTTAAGACAAAAAAGGGATTATTCAAATATAAGAAGAAACAGTTTGAGAAAAAAGTTTATATGCATATAAACTGAGAATAGTCACATGAATGTGGAAA | GTTTTCAAATATGAAAATTTTATAAGTGTCTGAGTATATTCATAAAAATGTATGAGATTATACGCGCAAATTAATATATAATCACTTATTTTTTCATTTA |
| | | | PBANKA_0506700 | citrate synthase, | Unstudied | TGTGTATATCGACTGATTTG | ACACCTAATAATGTGATTGG | AATATTTGCAATTTAAAGGGAAATTATATATTAGCAGTGTAGATATATTAATCCGCG | TTATATCAGAACTAATGGGAGAAAAAAAAACAATTTTTGATGTATATATGCATAGTT |

| | | | | | | | | | |
|---|---|---|---|---|---|---|---|---|---|
| | | | | mitochondrial, putative (CS) | | | | CGCAAAAAAAAAACGAAAGTGTAACACATAATTTTTTTGCAAAA | GGAAAAACGAGAAGGAATTTTTGTGAACATTTTTCTTCAATGC |
| | | | PBANKA_1112400 | orotate phosphoribosyltransferase, putative (OPRT) | Unstudied | ATAGATGATGTATTTACTTG | TTATTTGGAGCATCGTATAA | AATATTTAAGAAAAACATATTTATATTGACAAAATTTTTTATTGTTTATTTCACAATTTGAGATTTTTTTTTTTTGGCGATTGAAAAAACTTTGAAAAAA | TTTGGAAAAATTAAAATTAAAAAAAAAAATAGAGATAAAATAATGGTGATTTTTTTTTCTTTTTTAATCTGAGTTCTGTATTTACTTTCATAAGTTTTT |

**Table S6: PbHiT classification of phenotype compared to *Plasmo*GEM and *piggyBac* screens**

| | | | Disruptability | | |
|---|---|---|---|---|---|
| Target gene ID | Gene name | PlasmoGEM phenotype | | piggyBac phenotype | PbHiT phenotype |
| PBANKA_1459900 | signal recognition particle receptor subunit beta, putative (SRPRB) | Essential | Refractory | Possible | Not assigned |
| PBANKA_0804000 | 60S ribosomal protein L37, putative (RPL37) | Essential | Refractory | Refractory | Refractory |
| PBANKA_0809200 | ribosomal protein L35, apicoplast, putative | Essential | Refractory | Refractory | Refractory |
| PBANKA_0703200 | ribosomal protein L21, apicoplast, putative | Essential | Refractory | Refractory | Refractory |
| PBANKA_0505400 | conserved Plasmodium protein, unknown function | Essential | Refractory | Refractory | Refractory |
| PBANKA_0812300 | spindle and kinetochore-associated protein 1, putative (SKA1) | Essential | Refractory | Refractory | Not assigned |
| PBANKA_1009800 | cytochrome c oxidase assembly protein COX15, putative (COX15) | Essential | Refractory | Refractory | Refractory |
| PBANKA_1136500 | conserved Plasmodium protein, unknown function | Essential | Refractory | Refractory | Refractory |
| PBANKA_0914900 | onserved protein, unknown function | Essential | Refractory | Refractory | Refractory |
| PBANKA_0817800 | conserved Plasmodium protein, unknown function | Essential | Refractory | Possible | Refractory |
| PBANKA_1233900 | 50S ribosomal protein L14, mitochondrial, putative | Essential | Refractory | Refractory | Refractory |
| PBANKA_0514900 | ookinete surface protein P28 | Slow growers | Possible | Refractory | Not assigned |
| PBANKA_1362100 | tyrosine kinase-like protein, putative (TKL3) | Slow growers | Possible | Possible | Not assigned |
| PBANKA_1108400 | single-stranded DNA-binding protein, putative (SSB) | Slow growers | Possible | Refractory | Not assigned |
| PBANKA_1322400 | exonuclease V, mitochondrial, putative | Slow growers | Possible | Refractory | Not assigned |
| PBANKA_1426400 | mitochondrial carrier protein, putative | Slow growers | Possible | Possible | Not assigned |

| | | | | | |
|---|---|---|---|---|---|
| PBANKA_0314200 | calcium-dependent protein kinase 1 (CDPK1) | Dispensable | Possible | Refractory | Not assigned |
| PBANKA_0616700 | NIMA related kinase 4 (NEK4) | Dispensable | Possible | Refractory | Possible |
| PBANKA_0408200 | calcium-dependent protein kinase 3 (CDPK3) | Dispensable | Possible | Possible | Possible |
| PBANKA_1305200 | serine/threonine protein kinase, putative | Dispensable | Possible | Possible | Not assigned |
| PBANKA_1414500 | glycogen synthase kinase-3 alpha, putative (GSK3alpha) | Dispensable | Possible | Refractory | Possible |
| PBANKA_0926600 | protein GCN20, putative | Dispensable | Possible | Possible | Possible |
| PBANKA_1352600 | serine/threonine protein kinase, putative | Dispensable | Possible | Refractory | Not assigned |
| PBANKA_1421600 | calcium/calmodulin-dependent protein kinase, putative | Dispensable | Possible | Possible | Possible |
| PBANKA_0308500 | tyrosine kinase-like protein, putative (TKL1) | Dispensable | Possible | Possible | Possible |
| PBANKA_0604400 | targeted glyoxalase II, putative (tGLO2) | Dispensable | Possible | Possible | Possible |
| PBANKA_1013300 | mitogen-activated protein kinase 1, MAPK1 | Dispensable | Possible | Possible | Possible |
| PBANKA_1037800 | secreted ookinete adhesive protein, SOAP | Dispensable | Possible | Possible | Possible |
| PBANKA_0515000 | ookinete surface protein P25 | Dispensable | Possible | Possible | Possible |
| PBANKA_1401600 | methyltransferase, putative | Slow | Possible | Refractory | Not assigned |
| PBANKA_1034400 | plasmepsin IV, PM IV | Slow | Possible | Not assigned | Not assigned |
| PBANKA_1104200 | 2-oxoisovalerate dehydrogenase subunit beta, mitochondrial, putative, BCKDHB | Slow | Possible | Possible | Not assigned |
| PBANKA_0103700 | conserved Plasmodium protein, unknown function | unstudied | | Refractory | Possible |
| PBANKA_0812900 | conserved Plasmodium protein, unknown function | unstudied | | Refractory | Possible |
| PBANKA_0603650 | conserved Plasmodium protein, unknown function | unstudied | | Not assigned | Possible |

| | | | | |
|---|---|---|---|---|
| PBANKA_0409000 | conserved Plasmodium protein, unknown function | unstudied | Refractory | Possible |
| PBANKA_0502900 | conserved Plasmodium protein, unknown function | unstudied | Possible | Possible |
| PBANKA_0111400 | conserved Plasmodium protein, unknown function | unstudied | Possible | Possible |
| PBANKA_0524100 | conserved Plasmodium protein, unknown function | unstudied | Not assigned | Possible |
| PBANKA_0519200 | conserved Plasmodium protein, unknown function | unstudied | Not assigned | Possible |
| PBANKA_0509600 | conserved Plasmodium protein, unknown function | unstudied | Refractory | Possible |
| PBANKA_0315700 | conserved Plasmodium protein, unknown function | unstudied | Possible | Possible |
| PBANKA_0519400 | conserved Plasmodium protein, unknown function | unstudied | Not assigned | Possible |
| PBANKA_0519100 | conserved Plasmodium protein, unknown function | unstudied | Not assigned | Possible |
| PBANKA_0821200 | conserved Plasmodium protein, unknown function | unstudied | Refractory | Possible |
| PBANKA_0111500 | conserved Plasmodium protein, unknown function | unstudied | Possible | Refractory |
| PBANKA_0817400 | conserved Plasmodium protein, unknown function | unstudied | Refractory | Refractory |
| PBANKA_0408800 | conserved Plasmodium protein, unknown function | unstudied | Refractory | Refractory |
